# Near-Physiological-Temperature Serial Femtosecond X-ray Crystallography Reveals Novel Conformations of SARS-CoV-2 Main Protease Active Site for Improved Drug Repurposing

**DOI:** 10.1101/2020.09.09.287987

**Authors:** Serdar Durdagi, Cagdas Dag, Berna Dogan, Merve Yigin, Timucin Avsar, Cengizhan Buyukdag, Ismail Erol, Betul Ertem, Seyma Calis, Gunseli Yildirim, Muge D. Orhan, Omur Guven, Busecan Aksoydan, Ebru Destan, Kader Sahin, Sabri O. Besler, Lalehan Oktay, Alaleh Shafiei, Ilayda Tolu, Esra Ayan, Busra Yuksel, Ayse B. Peksen, Oktay Gocenler, Ali D. Yucel, Ozgur Can, Serena Ozabrahamyan, Alpsu Olkan, Ece Erdemoglu, Fulya Aksit, Gokhan Tanisali, Oleksandr M. Yefanov, Anton Barty, Alexandra Tolstikova, Gihan K. Ketawala, Sabine Botha, E. Han Dao, Brandon Hayes, Mengning Liang, Matthew H. Seaberg, Mark S. Hunter, Alex Batyuk, Valerio Mariani, Zhen Su, Frederic Poitevin, Chun Hong Yoon, Christopher Kupitz, Raymond G. Sierra, Edward Snell, Hasan DeMirci

## Abstract

The COVID19 pandemic has resulted in 25+ million reported infections and nearly 850.000 deaths. Research to identify effective therapies for COVID19 includes: i) designing a vaccine as future protection; ii) structure-based drug design; and iii) identifying existing drugs to repurpose them as effective and immediate treatments. To assist in drug repurposing and design, we determined two apo structures of Severe Acute Respiratory Syndrome CoronaVirus-2 main protease at ambient-temperature by Serial Femtosecond X-ray crystallography. We employed detailed molecular simulations of selected known main protease inhibitors with the structures and compared binding modes and energies. The combined structural biology and molecular modeling studies not only reveal the dynamics of small molecules targeting main protease but will also provide invaluable opportunities for drug repurposing and structure-based drug design studies against SARS-CoV-2.

**One Sentence Summary:** Radiation-damage-free high-resolution SARS-CoV-2 main protease SFX structures obtained at near-physiological-temperature offer invaluable information for immediate drug-repurposing studies for the treatment of COVID19.

## INTRODUCTION

In late 2019, after the first patient was diagnosed with pneumonia of unknown etiology reported to the World Health Organization (WHO) from China, millions of cases followed in a short span of four months (WHO). On 11th of March, 2020; WHO declared COVID19 outbreak as a pandemic, which originated from Severe Acute Respiratory Syndrome Corona Virus-2 (SARS-CoV-2) infection. SARS-CoV-2 has a high spread rate (R_o_) value, deeming the pandemic difficult to control (Petersen et al., 2020). Moreover, the absence of a vaccine to provide immunity and the lack of effective treatments to control the infection in high comorbidity groups make this pandemic a major threat to global health (Ahn et al., 2020).

The first human coronavirus to cause a variety of human diseases, such as common cold, gastroenteritis, and respiratory tract diseases was identified in the 1960s (Tyrrell et al., 1965). In 2002, a deadly version of coronavirus responsible for SARS-CoV was identified in China (Heymann et al., 2004). SARS-CoV-2, the most recent member of the coronavirus family to be encountered, is a close relative of SARS-CoV and causes many systemic diseases (Andersen et al., 2020; Dutta and Sengupta., 2020; Braun et al., 2020). COVID19 patients exhibit (i) high C-reactive protein (CRP) and pro-inflammatory cytokine levels; (ii) macrophage and monocyte infiltration to the lung tissue; (iii) atrophy of spleen and lymph nodes which weakens the immune system; (iv) lymphopenia; and (v) vasculitis (Zhang et al., 2020; McGonagle et al., 2020). Release of a large amount of cytokines results in acute respiratory distress syndrome (ARDS) aggravation and widespread tissue injury leading to multi-organ failure and death. Therefore, mortality in many severe cases of COVID19 patients has been linked to the presence of the cytokine storm evoked by the virus (Ragab et al., 2020).

The SARS-CoV-2 genome encodes structural proteins including surface/spike glycoprotein (S), envelope (E), membrane (M), and nucleocapsid (N) proteins; and the main reading frames named ORF1a and ORF1b that contain 16 non-structural proteins (NSP) (Gordon et al., 2020; Chen, Liu & Guo., 2020). Among these, ORF1a/b encodes Papain-like protease (PLpro), main protease (Mpro), a chymotrypsin-like cysteine protease, along with polyproteins named polyprotein1a (pp1a) and polyprotein1b (pp1b) (Astel et al.,2005). Encoded polyproteins are then proteolyzed to NSPs by precise Mpro and PLpro cleavages of the internal scissile bonds. NSPs are vital for viral replication, such as RNA-dependent RNA polymerase (RdRp) and Nsp13, which are used for the expression of structural proteins of the virus (Dai et al., 2020; Thiel et al., 2003; Ullrich & Nitsche., 2020; Ziebuhr et al., 2000). SARS-CoV-2 Mpro has no homologous human protease that recognizes the same cleavage site (Pillaiyar et al., 2016). Therefore, drugs that target its active site are predicted to be less toxic and harmful to humans (L. Zhang et al., 2020). High sequence conservation of Mpro provides minimized mutation-caused drug resistance (Jin et al., 2020). Given its essential role in the viral life cycle, the SARS-CoV-2 Mpro presents a major drug target requiring a detailed structural study.

Drug repurposing is a rapid method of identifying potential therapies that could be effective against COVID19 compared to continuous investigative efforts (e.g identification of new drugs and development of preventive vaccine therapies). The well-studied properties of Food and Drug Administration (FDA) approved drugs means such molecules are better understood compared to their counterparts designed *de novo*. A putative drug candidate identified by drug-repurposing studies could make use of existing pharmaceutical supply chains for formulation and distribution, an advantage over developing new therapies (Chen et al., 2020; Huang et al., 2020; Pushpakom et al., 2018; Jarada et al., 2020). In typical drug repurposing studies, approved drug libraries are screened against the active site or an allosteric site of target protein structures obtained by methods that have limitations in revealing the enzyme structure, such as cryogenic temperature or radiation damage (Wang et al., 2020; Kneller et al., 2020) and investigations that only involve screening of drugs currently on the market (Zhou et al., 2020; Choudhary et al., 2020; Beck et al., 2020; Wang, 2020). Current structural biology-oriented studies that display a repurposing approach to SARS-CoV-2 research focused on target-driven drug design, virtual screening or a wide spectrum of inhibitor recommendations (Rathnayake et al., 2020; Chauhan and Kalra, 2020; Chen et al., 2020; Muralidharan et al., 2020; Khan et al., 2020; Kumar et al., 2020; Joshi et al., 2020). It is critical to interfere with the ongoing pandemic, instead of time and resource-consuming drug design studies, drug repurposing studies with the existing drugs are crucial in short terms should be a priority.

The main purpose of our study is to reveal the conformational dynamics of Mpro, which plays a central role in the viral life-cycle of SARS-CoV-2. Linac Coherent Light Source (LCLS) with its ultrafast and ultrabright pulses enables outrunning secondary radiation damage. The updated high-throughput Macromolecular Femtosecond Crystallography (MFX) instrument of LCLS that is equipped with the new autoranging epix10k 2-megapixel (ePix10k2M) detector that provides a dynamic range of eleven thousand 8keV photons in fixed low gain mode to collect secondary radiation-damage-free structural data from Mpro microcrystals remotely at ambient-temperature (Sierra et al., 2019; Blaj et al., 2019; Van Driel et al., 2020).

Here we present two SFX structures of SARS-CoV-2 Mpro which provide structural dynamics information of its active site and their deep *in silico* analysis. Our results emphasize the importance of structure-based drug design and drug repurposing by using the XFEL structures. We obtained more relevant results of flexible areas over two different structures. Radiation-damage-free SFX method which enables obtaining the novel high-resolution ambient-temperature structures of the binding pocket of Mpro provides an unprecedented opportunity for identification of highly effective inhibitors for drug repurposing by using a hybrid approach that combines structural and *in silico* methods. Besides, structure-based drug design studies will be more accurate based on these novel atomic details on the enzyme’s active site.

## RESULTS

### Ambient-temperature XFEL crystal structures of SARS-CoV-2 Mpro reveal alternate conformations of the drug-binding pocket

We determined two radiation-damage-free SFX crystal structures of SARS-CoV-2 Mpro in two crystal forms at 1.9 Å and 2.1 Å resolutions with the following PDB IDs: 7CWB and 7CWC, respectively (**Fig. 1A, B**) (**Supplementary Table 1&2 and fig. S27**). The diffraction data collected remotely at the MFX instrument of the LCLS at SLAC National Laboratory, Menlo Park, CA (Sierra et al., 2019) (***Supplementary Methods: Data Collection and Analysis for SFX studies at LCLS***). We used an Mpro structure determined at ambient-temperature using a rotating anode home X-ray source (PDB ID: 6WQF; Kneller et al., 2020) as our initial molecular replacement search model for structure determination. Two high-resolution SFX structures obtained in different space groups were superposed with an overall RMSD of 1.0 Å **(fig. S1)**. They reveal novel active site residue conformations and dynamics at atomic level, revealing several differences compared to the prior ambient-temperature structure of SARS-CoV-2 Mpro that was obtained at a home X-ray source (Kneller et al., 2020) **(Fig. 1A, B)**.

**Fig. 1.**
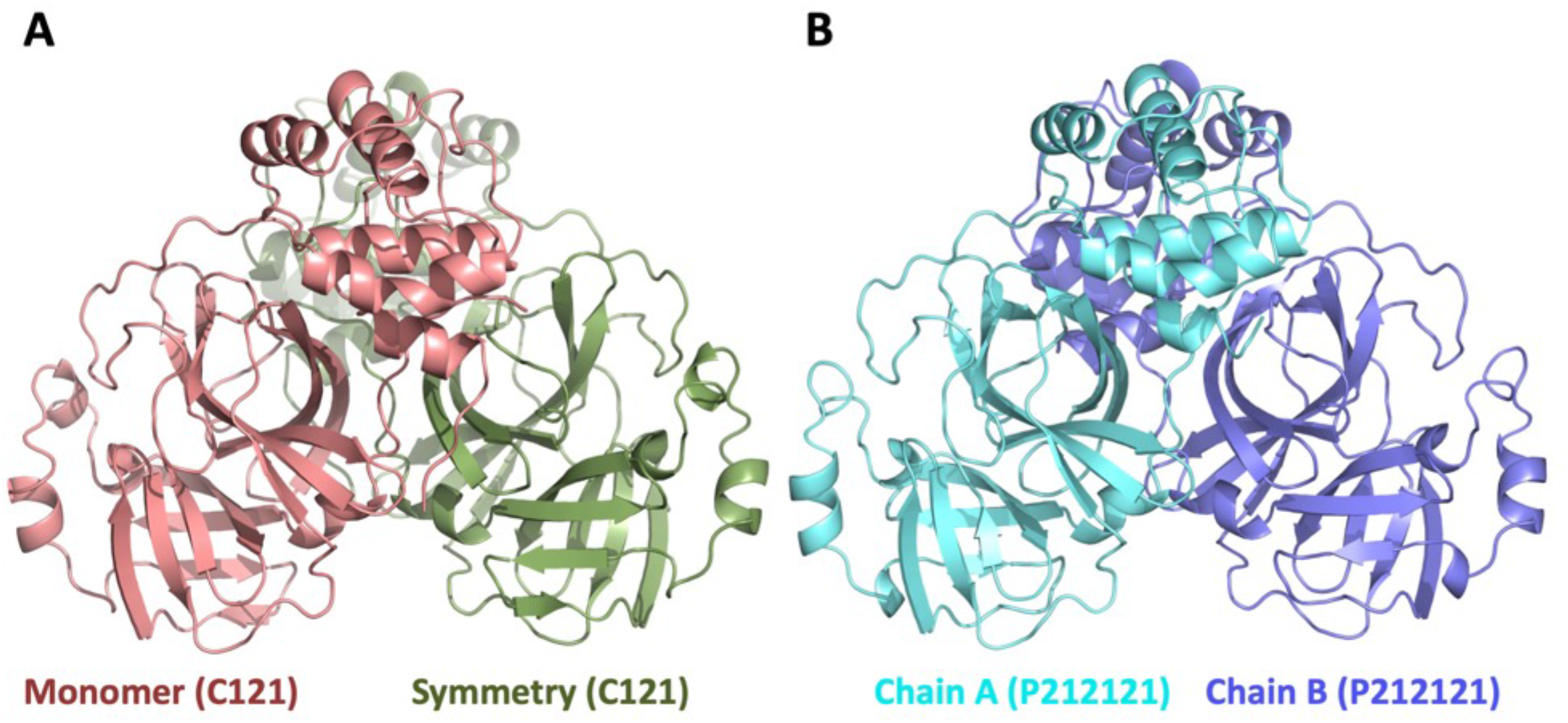
Overview of Mpro crystal structures. **A)** Cartoon representation of Mpro in space group C121. There is one molecule in the asymmetric unit cell colored in dark-salmon (left). Biologically-relevant dimeric form is generated by application of a symmetry operator which is colored in green. **B)** Cartoon representation of the Mpro in space group P2_1_2_1_2_1_ (right). Two chains of the dimer are colored in cyan and slate.

Mpro has a unique N-terminal sequence that affects enzyme’s catalytic activity (Chang, 2010; L. Zhang et al., 2020). Besides the native monoclinic form of Mpro at 1.9 Å (**Fig. 1A**, PDB ID: 7CWB) we also determined the structure of the Mpro with additional 4 extra N-terminal amino acids (generated by thrombin specific N-terminal cleavage) at 2.1 Å resolution (**Fig. 1B**, PDB ID: 7CWC). The structure obtained from this modified version of Mpro reveals how the minor changes introduced at the N-terminus affects both the three-dimensional structure of the Mpro and promotes the formation of a new orthorhombic crystal form. Biologically relevant dimeric structure of native monomeric Mpro can be generated by adding the symmetry-related chain B (**Fig. 1A**).

Each protomer of SARS-CoV-2 Mpro is formed by three major domains (Jin et al., 2020) **(fig. S2)**. Domain I starts from the N-terminal of protein and includes anti-parallel beta-sheet structure. This beta-sheet forms a beta-barrel fold which ends at residue 100. The domain II of Mpro resides between residues 101 through 180 and mostly consists of anti-parallel beta-sheets. The third domain of the Mpro is located between residues 181 through 306 and consists of mostly alpha helices and has a more globular tertiary structure **(fig. S2)**. The intersection between domain I, domain II and loop region of domain III is the key region of the enzyme **(fig. S2)** which forms the catalytic site and substrate-binding pocket of the enzyme. The biologically active form of SARS-CoV-2 Mpro is a homodimer (L. Zhang et al., 2020). Previous biochemical studies of Mpro suggested there is a competition for dimerization surface between domains I and III. In the absence of domain I, Mpro undergoes a new type of dimerization through the domain III (Zhong et al., 2008). During our purifications, we repeatedly observed a combination of monomeric and dimeric forms of Mpro on the size exclusion chromatography steps which may be caused by this dynamic compositional and conformational equilibrium.

The two SARS-CoV-2 Mpro SFX crystal structures reveal a non-flexible core active site and the catalytic amino acid Cys145 **fig. S3**. Temperature factor analysis revealed that the active site is surrounded by mobile regions **(fig. S3)**. The presence of these mobile regions was observed in both SFX crystal structures, suggesting an intrinsic plasticity rather than an artifactual finding that could have arisen based on the crystal lattice contacts **(fig. S3**; indicated with red circles). Further, this plasticity suggests that molecules that interact via non-covalent bonds to the Mpro binding pocket would do so weakly. Our investigation of PDB structures with available electron densities identified that the majority of the Mpro inhibitors formed covalent bonds with the active site residue Cys145. Additionally, few non-covalent inhibitors were identified and exhibited weak electron densities **(fig. S4.1 – S4.37).**

We compared our radiation-damage-free ambient-temperature SFX structure (PDB ID: 7CWB) with ambient-temperature X-ray structure (PDB ID: 6WQF). Structures were similar with an RMSD value of 0.404 Å **(Fig. 2A)**. However, we observed significant conformational differences especially in the side chains of Thr24, Ser46, Glu47, Leu50, Asn142, Cys145, Met165, and Gln189 residues **(Fig. 2B & fig. S5)**. The calculated bias-free composite omit map that covers the active region has been shown **(Fig. 3B & fig. S6)**. This structure offers previously unobserved new insights on the active site of Mpro in addition to 6WQF structure which is important for the future *in silico* modeling studies. All three domains contribute to the formation of the active site of the protein. The intersection part of the domain I residue His41, domain II residues Cys145 and His164 interact via a coordinated water molecule with Asp187 located at the N-terminal loop region of domain III to form the active site, that is an important drug target area **(Fig. 2C & fig. S7)**. The N-terminal loop of domain III is suggested to be involved in enzyme activity (Ma et al., 2020). The distance between Cys145 S*γ* and His41 Nε2 is 3.7 Å, very similar to the ambient-temperature structure (6WQF) (Kneller et al., 2020). Oδ1-Oδ2 atoms of Asp187 and NH2-Nε atoms of Arg40 contribute to a salt bridge between these two residues and stabilize the positions of each other. The W5 water molecule (indicated with a red sphere at the figure) in the active site plays crucial roles for catalysis. W5 forms triple H-bonds with His41, His164, and Asp187 side chains with the distances of 2.8 Å, 2.9 Å, and 2.8 Å, respectively **(Fig. 2C & fig. S7)**.

**Fig. 2.**
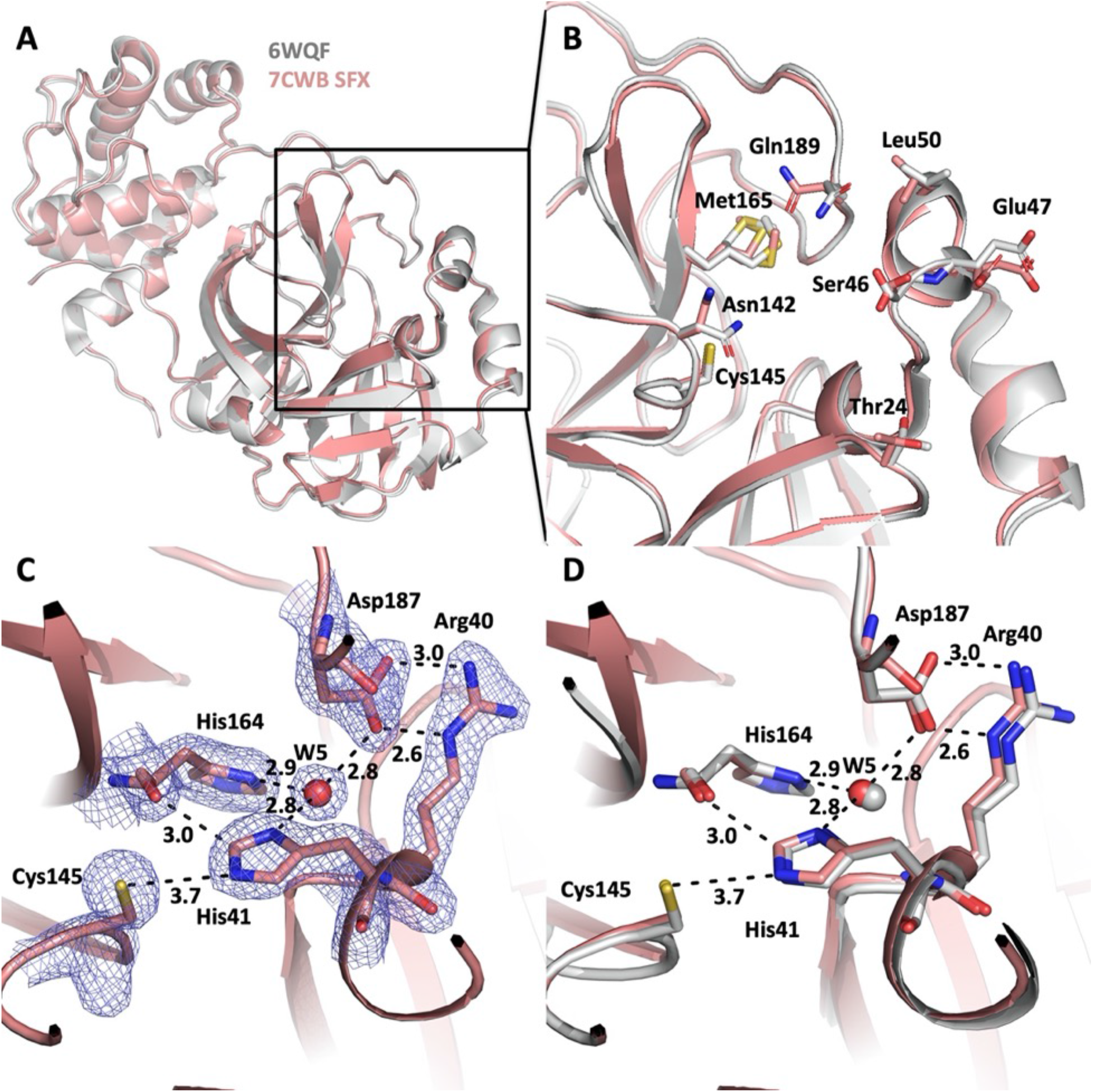
Comparison of SFX structure (PDB ID:7CWB) with ambient temperature structure (PDB ID: 6WQF). **A)** The two structures align very well with overall RMSD = 0.404 Å SFX structure shown in gray, ambient-temperature structure shown in pink. Same coloring scheme used as in all panels. **B)** Superposition of the predicted drug binding pocket reveals the significant conformational states. Residues with altered conformations were labeled and their positions were indicated. **C)** 2Fo-Fc electron density belonging to the active site residues are contoured in 1 sigma level and colored in slate. H-bonds and other interactions are indicated by dashed lines and distances are given by Angstrom. **D)** Superposition of the 7CWB (SFX) and 6WQF active site reveals very similar states.

When compared to the room temperature structure of Mpro (6WQF) our structure displays additional active site residue dynamics while it has an overall high similarity **(Fig. 2D)**. Canonical chymotrypsin-like proteases contain a catalytic triad composed of Ser(Cys)-His-Asp(Glu) in their catalytic region, however, SARS-CoV-2 Mpro possesses a Cys145 and His41 catalytic dyad which distinguishes the SARS-CoV-2 Mpro from canonical chymotrypsin-like enzymes (Gorbalenya & Snijder., 1996; Kneller et al., 2020). During the catalysis, the thiol group of Cys145 is deprotonated by the imidazole of His41 and the resulting anionic sulfur nucleophilically attacks the carbonyl carbon of the substrate. After this initial attack, an N-terminal peptide product is released by abstracting the proton from the His41, resulting in the His41 to become deprotonated again and a thioester is formed as a result. In the final step, the thioester is hydrolyzed which results in a release of a carboxylic acid and the free enzyme; therefore, restoring the catalytic dyad (Pillaiyar et al., 2016; Ullrich & Nitsche., 2020). Catalytic residue conformations of SFX structures are consistent with 6WQF and support the proposed Mpro catalytic mechanism **(Fig. 2C, D)**.

The crystal contact of the symmetry-related molecule with the N-terminal region of the Mpro is essential for the formation of the crystal lattice in the C121 space group (7CWB) **(Fig. 3A & fig. S8)**. There is an extensive network of H bonding interactions. The Ser1 amino group and carbonyl O atom engage in two H bonding interactions with carbonyl O and backbone amide N of Phe140. The carbonyl group of Gly2 forms a H bond with the carbonyl Oγ atom of Ser139. The backbone amide group of Phe3 forms a second H bonding interaction with the carbonyl Oγ atom of Ser139 in the C121 space group. The side chain amine and amino group of Arg4 form a H bond with carbonyl O atom of Lys137 **(fig. S8)**. These interactions are crucial in terms of whether the enzyme switches to the secondary crystal conformation in the P2_1_2_1_2_1_ space group. The elimination of these H bond networks by the addition of 4 N-terminal amino acids induced the formation of a second orthorhombic crystal form of SARS-CoV-2 Mpro. In the new crystal form in the P2_1_2_1_2_1_ space group, the binding pocket has no closeby crystal lattice contacts, as a result, has a wider binding pocket. This offers more possibilities for obtaining larger size drug soaks for future structural studies of Mpro drug complexes.

**Fig. 3.**
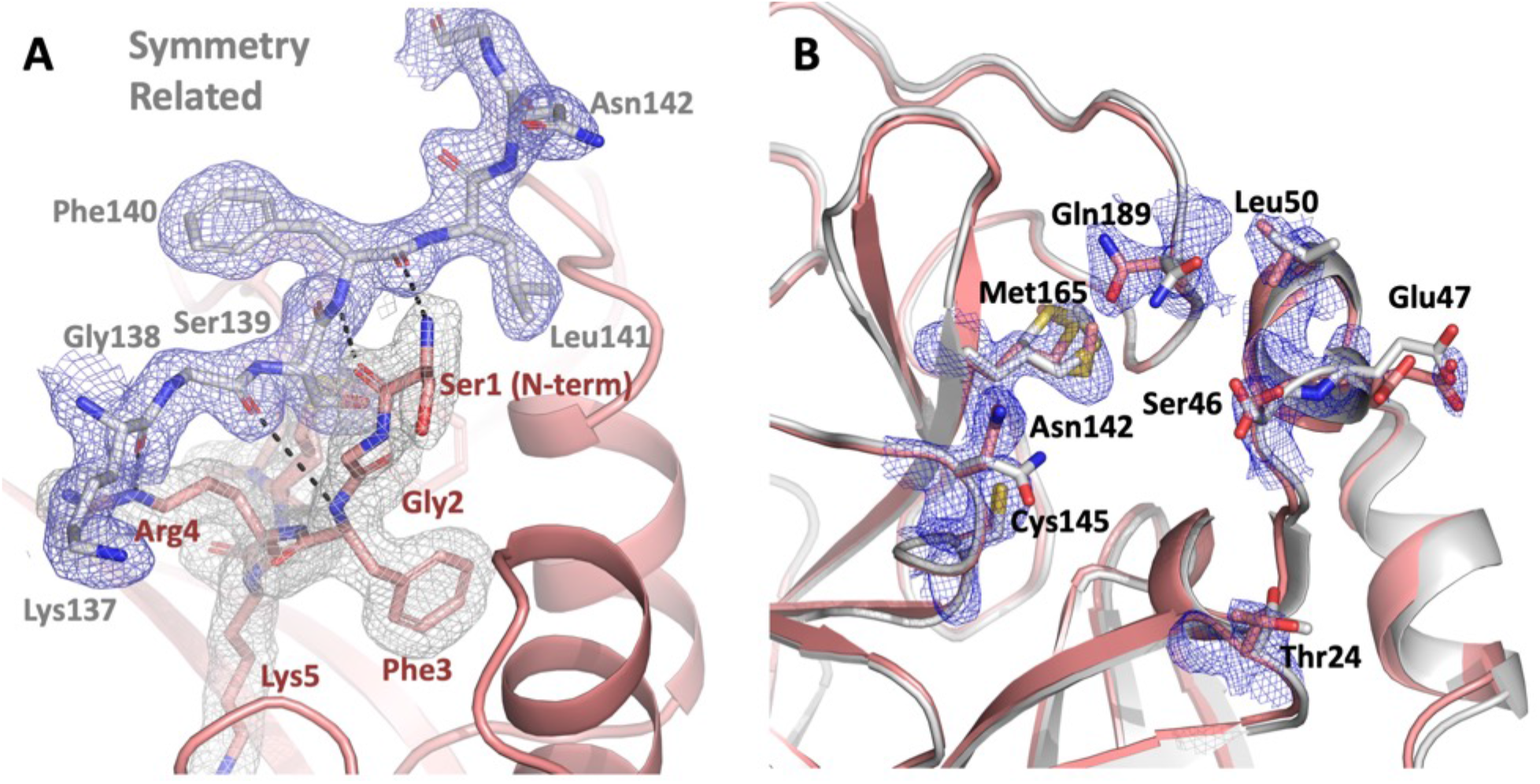
Key crystal contacts of C121 crystal form. **A)** The 2Fo-Fc electron density belongs to the N-terminal region of SARS-CoV-2 main protease contoured at 1 sigma level and colored in gray. Regions of the symmetry related molecule colored in gray, 2Fo-Fc electron density contoured in 1 sigma level and colored in slate. H-bonds are indicated by black dashed lines. Elimination of these H-bonds were essential to obtain the second crystal form in P2_1_2_1_2_1_ space group with more open inhibitor binding pocket for future soaking studies of the putative main protease inhibitors. **B)** Composite omit map of 7CWB active site.

### All-atom molecular dynamics (MD) simulations of apo forms of Mpro reveals protomers display asymmetric behavior

To better understand the structural and dynamical properties of SFX crystal structures in this study, MD simulations were performed using these newly determined crystal structures. As the catalytically active form of Mpro is a homodimer, we have used the dimer form of crystal structures. For the SFX structure obtained in space group C121 with a monomeric asymmetric unit, the relevant dimeric form is generated by symmetry operation. The MD simulations were run for 200 ns to elucidate the dynamic effects on the structures, especially the active site region. The evolutionary changes of atomic coordinates over time were monitored by calculating the RMSD for both chains, i.e. protomers (**fig. S9 & S10** for 7CWB and 7CWC, respectively). The root-mean-square fluctuations (RMSF) were also computed based on Cα atoms for both protomers separately (**fig. S11 & S12** for 7CWB and 7CWC, respectively) to determine the flexible regions. In simulations for both 7CWB and 7CWC, the protomers exhibit non-identical behavior as also observed by others in apo form of Mpro (Suarez and Diaz., 2020; Amamuddy et al., 2020) though as expected, higher RMSF values were observed for the loop regions of protomers. Additionally, the protomers of 7CWC display higher fluctuations compared to 7CWB around the loops covering the active site (such as loops containing residues 44-52 and 185-190) which could affect the accessibility of the active site by inhibitor compounds.

### Principal component analysis (PCA) correlates the inter-domain motions and its impact on the drug-binding pocket dynamics

The trajectory frames obtained from MD simulations were used to perform PCA to determine the variations of conformers of protein structures, i.e. to observe the slowest motions during MD simulations. As PCA and PCA-based methods are useful to reveal intrinsically accessible movements such as domain motions (Bahar et al., 2010), we have performed PCA for backbone atoms of dimeric units for the structures belonging to different space groups. We focused on the first three PCs, that show around 40% of the total variance in MD trajectories to determine the regions of protein structures that display the highest variation (**fig. S13 & S14** for 7CWB and 7CWC, respectively). The first three PCs were projected onto the protein structures to determine the contributions of each residue to specified PCs which displays the motions of specific regions with blue regions with higher thickness representing to more mobile structural parts of the protein along the specified PCs. It can be seen that both in C121 and P2_1_2_1_2_1_ space group structures, again protomers A and B display asymmetric behavior and domain movements along all considered PCs (**Fig. 4A-C** for 7CWB and **Fig. 4D-F** for 7CWC) which was also observed in residue dynamics for the same domain between alternate chains (Amamuddy et al., 2020). When the motions displayed by two dimeric forms are compared to each other 7CWB (**Supplementary movies M1-3** for 7CWB and **M4M6** for 7CWC), we observe that domain III is more mobile in 7CWB (the crystals in the C121 space group) compared to 7CWB (the crystals in the P2_1_2_1_2_1_ space group). However, the loop region containing residues 45-53 around the catalytic site is more mobile for both protomers in 7CWC while it is only mobile for the chain B of 7CWB. Interestingly, protomer A of 7CWC is more mobile compared to protomer A of 7CWB which could reflect the differences in dimeric interface due to the additional 4 N-terminal amino acids and missing interactions with chain B N-terminus residues in the dimer interface. However, the loop regions surrounding the binding pockets with residues 166-172 and 185-195 display higher flexibility in chain A of 7CWB while its mobility is somewhat restricted in 7CWC.

**Fig. 4.**
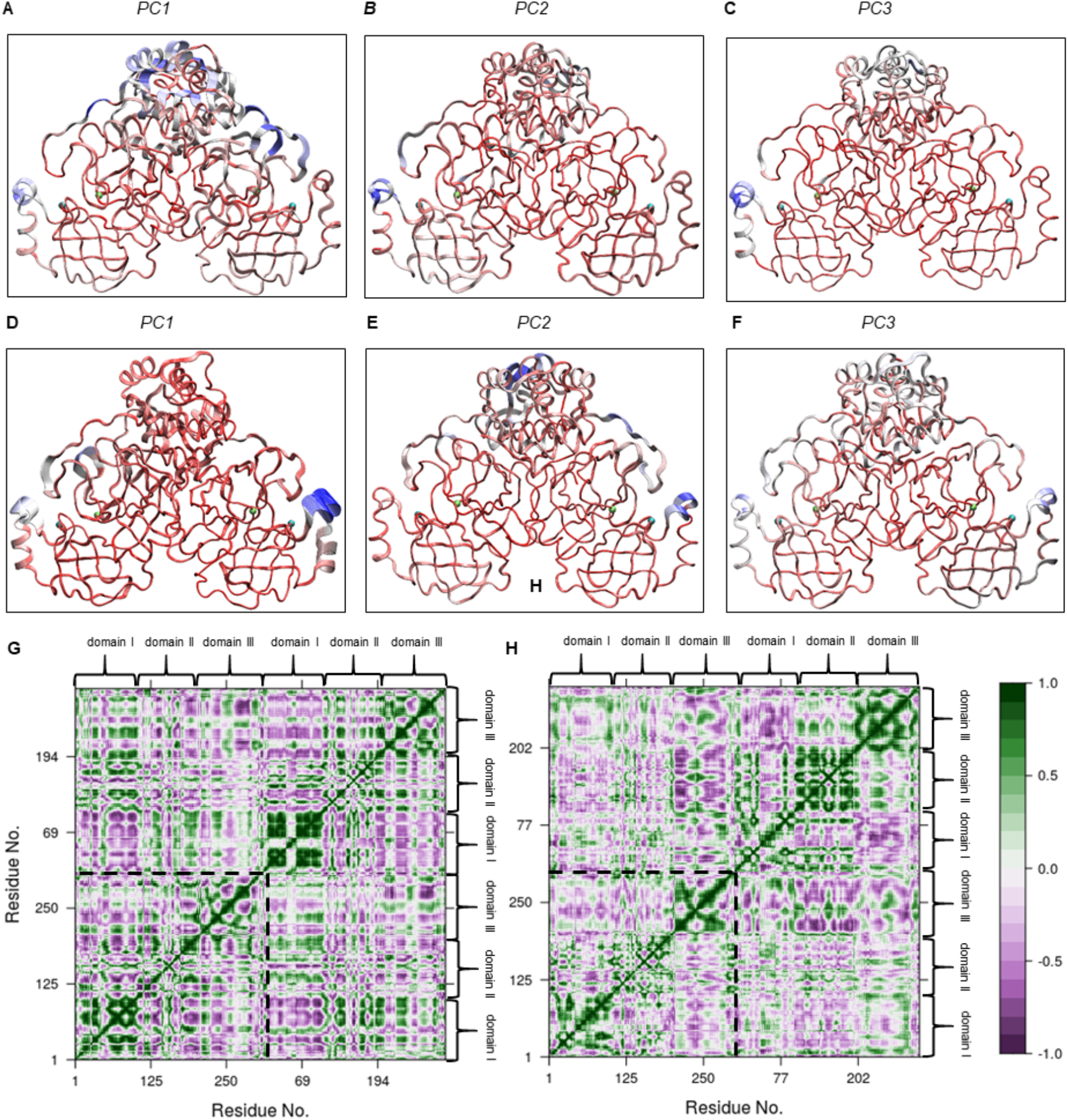
Contributions of residues to the first three PCs for **A-C)** 7CWB and **D-F)** 7CWC with chain A displayed on the right and chain B on the left side. The aligned trajectory frames were generated to interpolate between the most dissimilar structures in the distribution along specified PCs. Color scale from red to blue represents low to high displacements along specified PCs with broadening of the tubes depict the trajectory movements. The Cα atoms of catalytic residues His41 and Cys145 are displayed as spheres with cyan and lime colors, respectively. **G-H)** Dynamic cross correlation matrix generated from the motions observed in PC space with values ranging from −1 (complete anticorrelation) to +1 (complete correlation) **G)** for 7CWB and **H)** for 7CWC. The boundary between chain A and B is denoted by dashed lines.

We have also performed cross correlation analysis of residues along the PC space to understand the correlation between motions especially for different domains. The correlation between the motions along the first three PCs were plotted as dynamical cross correlation maps displayed in **Fig. 4G & 4H** for 7CWB and 7CWC, respectively. Focusing on the correlation between motions within protomers initially, we see that motions of domain I and III as well as domain I and II are mostly anti-correlated as shown by the purple regions in **Fig. 4G** for 7CWB and **Fig. 4H** for 7CWC. While we observed similar behavior for protomer A and B of 7CWC, the directions of domain I and III movements are different in protomers A and B of 7CWB, specifically for residues 189-192 that link are at the linkage points of domain II and III and residues 45-50 of domain I bordering the catalytic site. While in protomer A, these loops move in the same directions, their movements are mostly anti-correlated in protomer B for the slowest motion along PC1 **(Supplementary movies M1, Fig. 4G)**. This could cause differences in the accessibility of catalytic sites in a time dependent manner in protomers of 7CWB. On the other hand, domain II and III within protomers have mixed correlations with each other in which motions of some residues are along the same directions, others are in opposite directions within both protomers A and B of 7CWB while domain II has limited mobility as being more buried than other domains (Suarez and Diaz., 2020) except for the β-strand segment of residues 166-172 of protomer A. Surprisingly, this segment also has some mobility albeit limited in protomer B of 7CWC instead of protomer A **(Supplementary movies M4, Fig. 4H)**. Glu166 of this segment is actually an important residue that plays a role in stabilizing the substrate binding site S1 by interacting with Ser1 of the alternate protomer along with Phe140 (Ghahremanpour et al., 2020; Jin et al., 2020) and we observed that along PC1, for protomer A of 7CWB and protomer B of 7CWC, the motions of these residues are correlated. This could have implications about information transfer from one protomer to the other though other Mpro structures of SARS-CoV-2 needs to be studied in atomistic details. In addition, we observe that correlations between domains of protomers A and B for 7CWB and 7CWC are dissimilar. For instance, domains I of 7CWB have defined anti-correlated motions as can be seen from the purple areas in **Fig. 4H** while in 7CWC, the movements of domains I are more ambiguous. There are also differences in the behavior of domain I of protomer A and domain II of protomer B in the dimer structures in which for 7CWB the motions of these domains are positively correlated while they are anti-correlated in 7CWC.

### Different non-covalent interactions stabilize the dimer interfaces in different space groups

The structures that were obtained in this study, compared with four other Mpro structures crystallized recently. Selected Mpro structures were cryogenic apo form (PDB IDs: 6Y2E; Zhang et al., 2020 & 7C2Y), holoprotein with non-covalent inhibitor, X77 (PDB ID: 6W63) and room temperature structure (PDB ID: 6WQF; Kneller et al., 2020). Space groups of these structures are C121, P2_1_2_1_2_1_, P2_1_2_1_2 and I121 respectively. The completeness of the structures was also evaluated, 6WQF, and 6Y2E crystallized in full-length sequence (1-306), however, 6W63 only lacks Gln306 at both C-terminal ends of its chains. Among compared structures, 7C2Y, which has the same space group as 7CWC has, and lacks 24 amino acids (chain A, 1-3, 141-142, 281-283, and 298; chain B, 1-2, 45-50, 139-142, and 277-279). In our structures 7CWB has a full-length sequence, however, 7CWC lacks its last 6 amino acid residues from C-terminal in chain A, and starts with Phe3 (lacks Ser1 and Gly2), and ends at Ser301, lacks the last 5 residues. When six structures (6W63, 6Y2E, 6WQF, 7C2Y, 7CWB, and 7CWC) were compared based on hydrogen bonding interactions at the dimerization interfaces, more similarities were observed for the latter three (7C2Y, 7CWB, and 7CWC). The only difference between our structures and the other four is the hydrogen bond between Ser139 (chain A) and Gln299 (chain B) **(fig. S15)**. Although this hydrogen bond is not observed in other structures, the corresponding residues are close to each other, however they are not within hydrogen bonding distance **(fig. S16)**.

We also monitored all interface interactions throughout the MD simulations and compared 7CWB and 7CWC. Interestingly, the hydrogen bond between Ser139 (chain A) and Gln299 (chain B) was lost and the interaction was turned into a van der Waals interaction in 7CWB, and conserved as by 64% of the simulation time. In 7CWC structure, Ser139 (chain A) was not in the vicinity of Gln299 (chain B) **(fig. S17)**. More interestingly, the same interaction but this time between Gln299 (chain A) and Ser139 (chain B), that is present at the 7CWB crystal structure, was only observed 16% of the simulation time (**fig. S18**). In the 7CWC crystal structure, there was no hydrogen bonding interaction between Gln299 (chain A) and Ser139 (chain B). However during simulation, this bond was formed and retained in 65% of the simulation time **(fig. S18)**.

When static structures were compared based on hydrogen bond formation analysis, 7C2Y, 7CWB, and 7CWC clustered together, having the same hydrogen bonding network at the dimerization interface. The explanation for the hydrogen bond differences observed at the interface is the lack of amino acids at the N-terminal and C-terminal ends of the 7C2Y and 7CWC structures, compared to the 7CWB.

Simulation trajectories of 7CWB and 7CWC were compared based on hydrogen bond occupancies. The major differences were Lys137 (chain A) and Arg4 (chain B), Asn142 (chain A) and Ser301(chain B), and Arg298 (chain A) and Tyr118 (chain B). The first interaction was not observed in 7CWC, however the latter two interactions were not presented in 7CWB. Lys137 (chain A) and Arg4 (chain B) was conserved 82% of the simulation time; Asn142 (chain A) and Ser301 (chain B); and Arg298 (chain A) and Tyr118 (chain B) retained 67% and 72% of the obtained trajectory frames, respectively **(figs. S18 and S19)**. We also observed that during MD simulations, in the case of 7CWC, Gln299 (chain A) and Phe140 (chain B) come closer to each other, and this van der Waals contact was conserved 64% of the simulation time and not observed in the case of 7CWB **(fig. S19)**.

### Non-covalent bound inhibitors do not mainly form structurally stable complexes with Mpro

We used three well-known SARS-CoV-2 Mpro inhibitors (i.e., Ebselen, Tideglusib, and Carmofur) at the investigation of ligand-target interactions. These compounds were docked to the 7CWB and 7CWC structures and all-atom MD simulations were performed for the top-docking poses. These three compounds were also docked to structures with the following PDB IDs: 6W63 and 6Y2E, for comparison. MD simulations were performed using the same MD protocol. Results showed that especially for apo form dimer targets, Ebselen and Carmofur are quite flexible at the binding pocket throughout the simulations and in most of the cases, they do not form a stable complex structure **(fig. S20)**. Tideglusib has a more stable structure at the binding pocket of Mpro, however, its binding modes are different **(Fig. 5)**. In the following section, details of MD simulations results of Tideglusib at the binding site of the Mpro will be discussed.

**Fig. 5.**
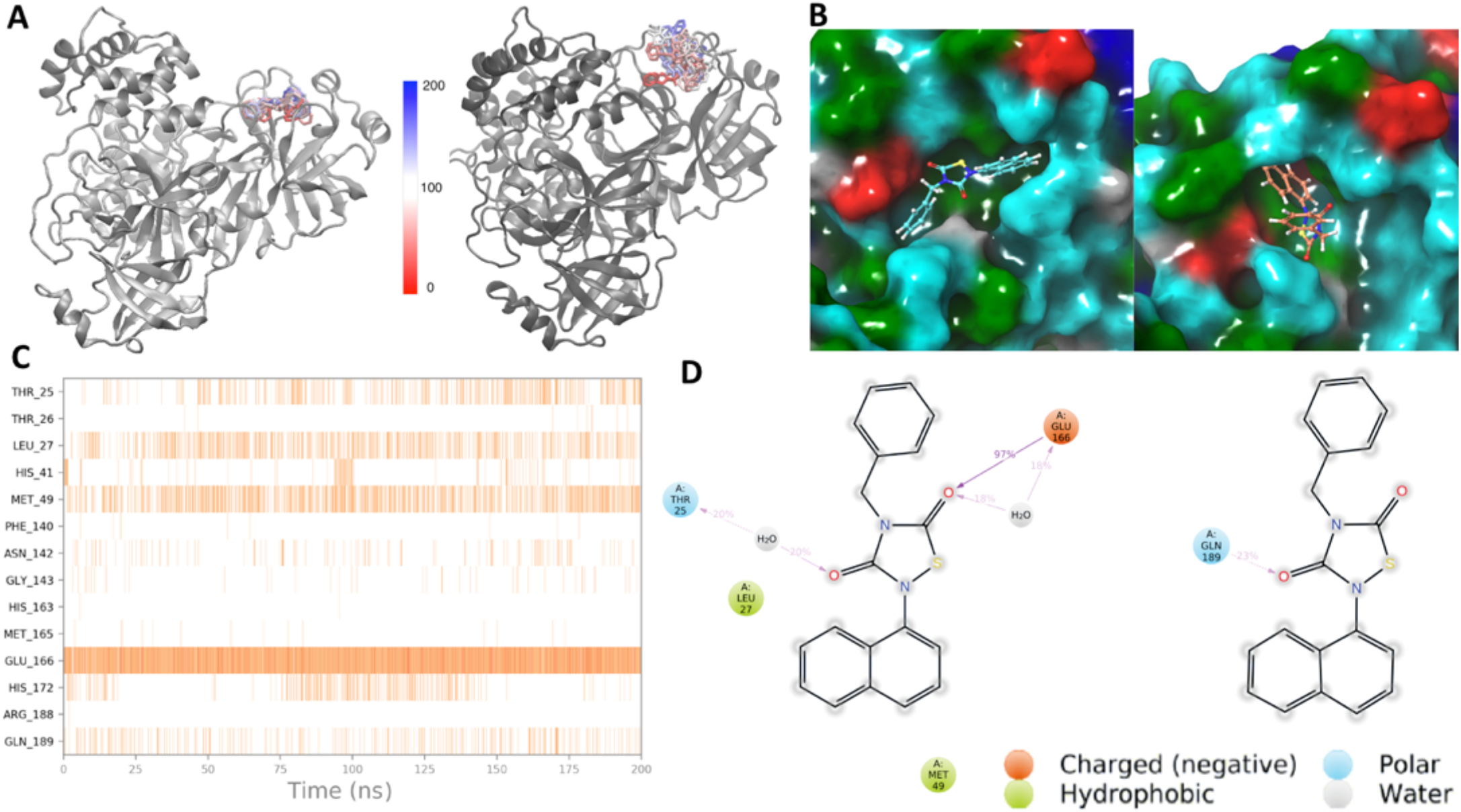
Trajectory analyses of Tideglusib bound 7CWC and 6Y2E PDB coded structures. **A)** Conformational changes of Tideglusib at the binding pockets of 7CWC (left) and 6Y2E (right) throughout 200 ns MD simulations. **B)** Representative binding poses of Tideglusib at the 7CWC (left) and 6Y2E (right). Residues are colored with amino acid types: red, negative charged; blue, positive-charged; green, hydrophobic; cyan, hydrophilic; **C)** Protein-ligand contacts throughout the MD simulations, 7CWC; **D)** 2D ligand interactions diagrams of Tideglusib, 7CWC (left); 6Y2E (right).

Comparison of binding pocket volumes of 7CWC and 6YE2 shows that latter has a bigger average binding pocket volume. Average binding pocket volumes were 174 and 142 Å3, for the Tideglusib bound 6YE2 and 7CWC structures, respectively. Corresponding solvent accessible surface area (SASA) values throughout the MD simulations also support this result. The SASA, which is the surface area of a molecule accessible by a water molecule, of 7CWC is smaller than 6Y2E **(fig. S21).**

MD simulations were performed for one of the well-known SARS-CoV-2 Mpro inhibitor Tideglusib. Tideglusib initially docked to the binding pockets of 6Y2E and 7CWC structures using induced-fit docking (IFD) approach. Top-docking poses of these compounds were then used in allatom MD simulations using the same MD protocols. While Tideglusib was structurally very stable at the binding pocket of the 7CWC during the simulations, it was not so stable at the 6Y2E the binding site **(Fig. 5A)**. Representative trajectory frames (i.e., the frame that has the lowest RMSD to the average structures) were used in the comparison of binding modes **(Fig. 5B)**. Results showed that while the binding mode of Tideglusib forms hydrogen bonds and pi-pi stacking interactions with Glu166, Gln189, and His172, respectively at the 7CWC; its corresponding binding mode at the 6Y2E only forms van der Waals type interactions with hydrophobic moieties. A timeline protein-ligand contacts were visualized throughout the simulations **(Fig. 5C).** Results showed that Thr25, Leu27, Met49, Glu166, and Gln189 form stable interactions with the ligand. 2D ligand atom interactions with protein residues are also represented **(Fig. 5D)**. Interactions that occur more than 15% of the simulation time are shown. However, corresponding interactions of the ligand at the 6Y2E were not stable **(fig. S22).**

### MD Simulations of Tideglusib with the 7CWB and 6WQF

For the comparison of monomer forms of Mpro, we applied IFD protocol to predict the binding mode of Tideglusib at the binding pocket of 6WQF and 7CWB. Carbonyl oxygens of the thiadiazolidine ring of the Tideglusib formed hydrogen bonds between Asn142, Gly143, and Glu166 from their backbone atoms. The naphthalene ring of the ligand formed a pi-pi stacking interaction with the His41 in the 7CWB structure. A similar binding mode of Tideglusib was observed when 6WQF was used. However, in this binding mode, His41 and Gly143 are found to be important. A backbone hydrogen bond was observed between one of the carbonyl oxygen of the thiadiazolidine ring and Gly143. His41 formed two pi-pi interactions between the benzyl and the naphthalene rings of the Tideglusib **(fig. S23A, B)**. These two binding modes predicted by IFD were used in 200 ns classical all-atom MD simulations. Each Mpro-Tideglusib system is evaluated based on RMSD changes from the average structure and we obtained representative structures from corresponding trajectories. In the representative structure of 7CWB-Tideglusib complex, we observed van der Waals interactions between surrounding residues, Thr25, His41, Cys44, Thr45, Ser46, Met49, Asn142, Cys145, His163, His164, Met165, Glu166, and Gln189. In the representative structure of 6WQF-Tideglusib complex, in addition to the similar van der Waals interactions, Asn142 and Gln189 formed two hydrogen bonds with the carbonyl oxygens of the thiadiazolidine ring from their backbone atoms **(fig. S23C, D)**. The alignment of the representative frames from 6WQF and 7CWB yielded an RMSD value of 0.97 Å **(fig. S24)**. Translational and rotational motions of the Tideglusib were also examined by the trajectory analyses. Conformational spaces that were explored by the Tideglusib were given in **fig. S25**. We also monitored the binding cavity volume throughout the MD simulations in both complexes and obtained the average values as 154 and 127 Å^3^ for the Tideglusib-bound 7CWB and 6WQF structures, respectively **(fig. S26)**. In the IFD poses, corresponding binding cavity volumes were calculated as 251 and 238 Å^3^, respectively. Decreased binding cavity volume during the MD simulations may be correlated with the unstable conformations of the Tideglusib in both structures. Overall, from the MD simulations of Tideglusib with the monomer forms of Mpro, we observed more stable conformations of Tideglusib at the binding pocket of 6WQF compared to the 7CWB.

## Discussion

SFX utilizes micro-focused, ultrabright and ultrafast X-ray pulses to probe small crystals in a serial fashion. Structural information is obtained from individual snapshots; capturing Bragg diffraction of single crystals in random orientations (Martin-Garcia et al., 2016). The main advantages of SFX over its counterparts are the capability of working with micron to nanometer sized crystals which does not necessitate the lengthy and laborious optimization steps and enables working with multiple crystal forms and space groups. It enables obtaining high resolution structures at physiologically meaningful temperature and confirms the dynamic regions of the active site without secondary radiation damage. SFX offers great potential and provides much needed critical information for future high-throughput structural drug screening and computational modeling studies with sensitivity to dynamics, insensitivity to potential radiation-induced structural artifacts resulting in production of detailed structural information.

SARS-CoV-2 Mpro catalyzes the precise cleavage events responsible for activation of viral replication and structural protein expression (Jin et al., 2020). It has been the focus of several structural and biochemical studies, many of which have been performed at cryogenic temperatures, aiming to provide better understanding of the active site dynamics and reveal an inhibitor that affects the enzyme based on structural information. Two crystal forms of Mpro, native and modified, were determined at ambient-temperature with resolutions of 1.9 Å and 2.1 Å respectively. The two forms produced are optimal for co-crystallization and soaking respectively. Co-crystallization experiments provide efficient interaction in the binding pocket as both drug and protein are stabilized before the formation of crystals. Due to the close crystal lattice contacts, co-crystallization seems to be the preferred method for the native form of Mpro (C121). The N-terminal of Mpro which plays a critical role in crystal packing. Elimination of the H-bonding network produced orthorhombic crystals in the P2_1_2_1_2_1_ space group yielding a wider binding pocket, increasing the probability of capturing an expanded number of protein-drug complexes by soaking. An added advantage of two different crystal forms was the elimination of the artifacts introduced by specific lattice packing restraints of each crystal form in the dynamics analysis.

The high-resolution Mpro SFX structures presented here in two different crystal forms collectively revealed the intrinsic plasticity and dynamics around the enzyme’s active site. Due to the anionic nature of Cys145, it seems challenging to design molecules that interact with the active site only through non-covalent bonds as it has a very flexible environment. These findings provide a structural basis for and is consistent with studies claiming that the majority of inhibitors form covalent bonds with the active site of Mpro (Dai et al., 2020; Jin et al., 2020), mainly through Cys145 **(figs. S4.4–S4.37)**. Especially, unlike some studies, ebselen does not form a stable complex structure (Sies et al., 2020; Menendez et al., 2020; Jin et al., 2020; Zmudzinski et al., 2020; Węglarz-Tomczak et al., 2020). In addition to the importance of Cys145 residue in our two different structures, a coordinated W5 molecule, regulating the catalytic reaction via triple hydrogen-bonding interactions with the His164, and the Asp187 that stabilizing the positive charge of His41 residue (Kneller et al., 2020). This active site residue conformations of SFX structures are consistent with previous ambient-temperature structure (6WQF) **(Fig. 2C, D)**.

Gln189 contributes to the stability of the ligands (Dai et al., 2020; Kneller et al., 2020), along with Asn142 and Ser46, are the active site residues forming the flank of the cavity **(Fig. 6)**. In a recent study, Asn142 and Gln189 have been indicated to interact with 11a and 11b inhibitors that have proven *in vitro* effectiveness (Dai et al., 2020). Along with these amino acids, Thr24, which makes van der Waals interaction with the N3 inhibitor, and Ser46 which undergoes conformational changes in the presence of this inhibitor (PDB ID: 6LU7, Jin et al., 2020) and Leu50 binding to N3, gave different side-chain conformations with SFX, according to 6WQF structure (Kneller et al., 2020) **(Fig. 2B)**. There were differences in the crystallization conditions between the SFX and 6WQF studies, the former making use of charge effects and the latter molecular crowding. The SFX study is also secondary radiation damage-free, eliminating potential artifacts to examine crucial amino acids at the atomic level, especially in terms of catalytic and inhibitor binding sites **(figs. S5–S7)**. These two high-resolution SFX structures in different space groups reveal new active site residue conformations and intra- and inter-domain network and their dynamics at the atomic level, which helps us to better understand any related structural allosteric transitions of Mpro structure interacting with the inhibitors **(Fig. 5 & figs. S23–25)**. Therefore, considering the importance of the required sensitivity in drug design or the use of natural compounds studies, these active site residue conformations reveal the critical importance of our study more clearly.

**Fig. 6.**
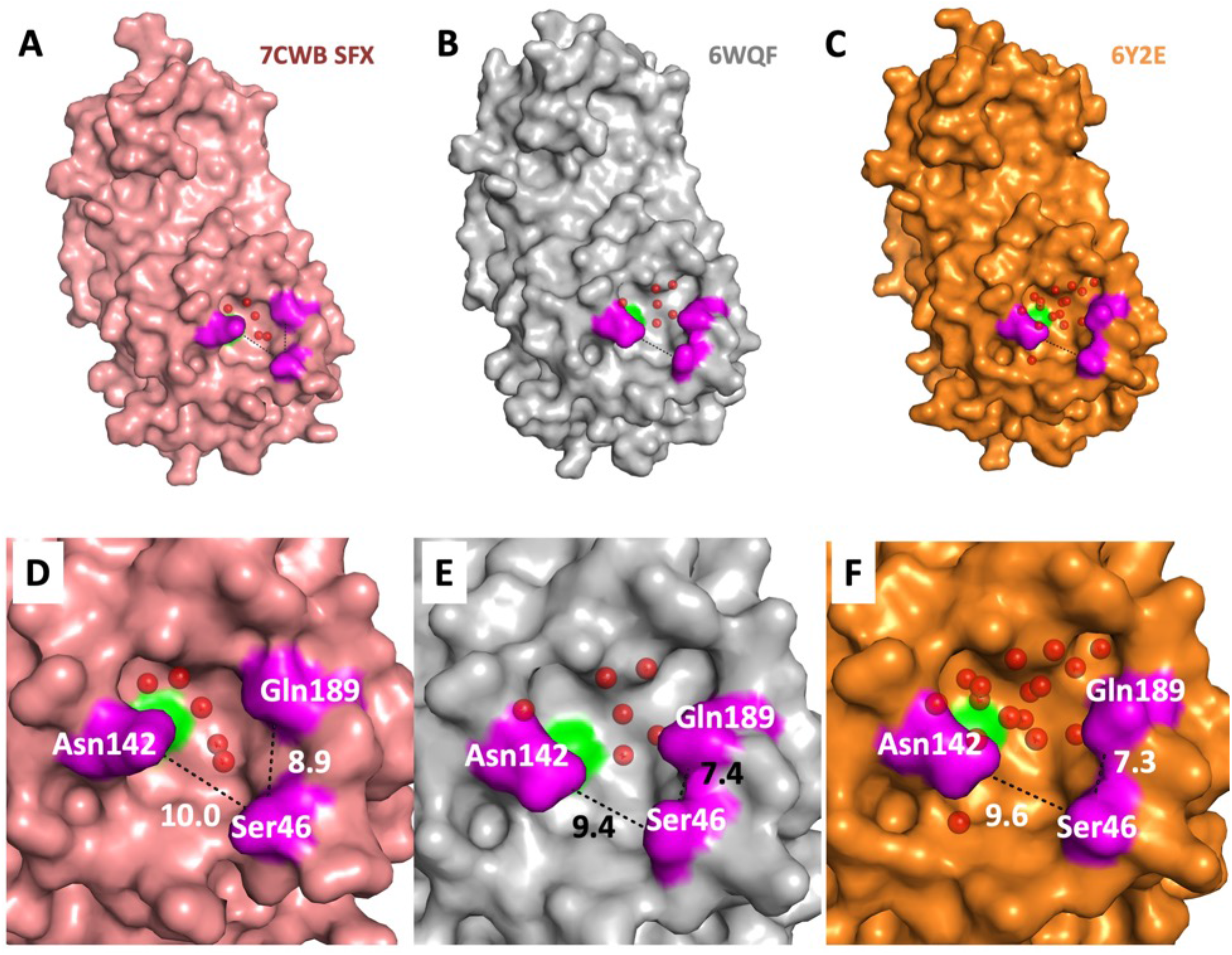
Different monoclinic crystal forms of the SARS-CoV-2 Mpro with catalytic site cavities. **A)** C121 SFX, **B)** 6WQF, and **C)** 6Y2E, respectively. Water molecules in the catalytic cavity are shown with red spheres. The catalytic residue (Cys145) is highlighted in green and the flank cavity residues (Asn142, Ser46, Gln189) are highlighted in purple in the catalytic cavity of each crystal form **D)** C121 SFX, **E)** 6WQF, and **F)** 6Y2E, respectively. Distance between C (Asn142 Cγ-Ser46 Cβ-Gln189 Cδ) atoms of flank cavity residues is shown as a dashed line. All distances are given in Angstrom (Å).

There are many *in silico* docking studies performed based on cryogenic Mpro protein structures of SARS-CoV-2 (Jin et al., 2020; Ton et al., 2020; Dai et al., 2020; Pillaiyar et al., 2020; Durdagi et al., 2020). Although potent antiviral drug candidates are identified and a vaccine research is still ongoing, they have not yielded a desirable final treatment/cure yet. In pursuit of effective drugs against COVID19, the key role of Mpro in viral replication of SARS-CoV-2, highly conserved structure, and the low toxicity of the antiviral molecules targeting this protein due to the absence of homolog of this protease in humans made Mpro the target of our study. Having access to the alternative ambient-temperature structures of Mpro and observed conformational changes on active site residues will be a significant boon for the development of therapeutics and provide better understanding on ligand and inhibitor binding. At this point, our work has two original aspects. Firstly, we used a comprehensive platform, SFX, which helps to deeply understand the complexity of SARS-CoV-2 to gain access to the high-resolution and radiation damage-free structure and the structural dynamics of the target protein Mpro at an unprecedented level at near-physiological-temperatures. Secondly, we determined two high-resolution SFX structures of SARS-CoV-2 Mpro in two different space groups due to the new high-throughput data collection setup offered by the MFX instrument of the LCLS-II.

Drug repurposing has been the preferred area of research in the insufficiency of time and resources in emergency cases such as a novel pandemic. The most important advantage of the repurposing research is the bypass of several lengthy early stages in drug development. Also, knowing the beneficial and detrimental effects of targeted drugs, along with well-established precautions will help with time limitations. This procedure has emerged as a fundamental and very strategic approach not only for prospective cohort design but also for many types of clinical trials, particularly crosssectional studies because as the molecules considered in repurposing studies passed through several stages, have well-defined profiles, they would not require prolonged pre-clinical studies, and hence they are important candidates to consider in case of disease emergencies or outbreaks (Su et al., 2017; Cavalla et al., 2013; Choo et al., 2019; Aguila et al., 2020). Therefore, enormous contribution to clinical studies, repurposing is a rapid step towards the conclusion of not only randomized controlled trials but also critical structural biology investigations (Bumb et al., 2015; Pihan et al., 2012; Choudhary et al., 2020). Considering all this, adopting the drug repurposing approach and using known inhibitors ebselen, tideglusib, and carmofur to carry out Mpro-based *in silico* molecular docking, MD simulations and post-MD analyses make our combined study more specific.

A grave issue with drug research is the variability of the target proteins as the mutations and modifications may render the found drugs ineffective (Dinesh et al., 2020). For this reason, it is only sensible to work on a protein, whose biochemical properties are conserved over time and among different strains. Evolutionarily, viruses try to hide by mimicking the proteins involved in the functioning of the host organism. Cytomegalovirus (HCMV) can mimic a common host protein to hijack normal cell growth machinery or human immunodeficiency virus (HIV) can mimic a high percentage of human T cell receptors (Bernstein, 2017; Robertson, 2003). SARS-CoV-2 virus contains the PLpro enzyme, which is highly similar to the deubiquitinating enzymes (USP7, USP14) in human metabolism (Ratia et al., 2014). That’s why targeting this enzyme carries a high risk. In addition, Spike protein, one of the other targeted proteins, also has a high mutation rate (Jia et al., 2020). Also, Spike protein includes a similar restriction site with epithelial channel protein (Anand P et al., 2020). Besides that, inhibitors which target Spike protein may act only for preventive aims (i.e., before the virus infection), if the virus already infected the host cell, targeting this region may not be useful. There is no Mpro homolog in the human genome, targeting this protease is therefore safer and harmless to humans with reduced cross reactivity and side effects making it an ideal candidate for drug therapy.

## Acknowledgements

HD acknowledges support from National Science Foundation (NSF) Science and Technology Centers grant NSF-1231306 (Biology with X-ray Lasers, BioXFEL) and The Scientific and Technological Research Council of Turkey (TUBITAK) grant (118C270). HD would like to thank Michelle Young, Ritu Khurana, Lori Anne Love and Tracy Chou for their invaluable support and discussions. Use of the Linac Coherent Light Source (LCLS), SLAC National Accelerator Laboratory, is supported by the U.S. Department of Energy, Office of Science, Office of Basic Energy Sciences under Contract No. DE-AC02-76SF00515. The HERA system for in helium experiments at MFX was developed by Bruce Doak and funded by the Max-Planck Institute for Medical Research. Research was supported by the DOE Office of Science through the National Virtual Biotechnology Laboratory, a consortium of DOE national laboratories focused on response to COVID-19, with funding provided by the Coronavirus CARES Act. The numerical calculations reported in this paper were partially performed at TUBITAK ULAKBIM, High Performance and Grid Computing Center (TRUBA resources). SD and IE acknowledge TRUBA for the computational resources. SD acknowledges support from Bahcesehir University (BAU) Scientific Research Projects (BAP) grant (BAU2020-0101).

## Materials and Methods

### Gene Construct Design and Cloning

#### Construct-1 with Native N- & C-terminals

The published Severe Acute Respiratory Syndrome CoronaVirus-2 main protease (SARS-CoV-2 Mpro) crystal structure revealed that the N-terminal serine residue is involved in the critical C121 space group crystal lattice contact (PDB ID 6LU7 Jin et al., 2020). We designed the native Mpro construct with the following amino acid sequence to obtain the native enzyme with no modifications at the N- & C-terminal ends. MSAVLQ(native_Mpro_cleavage_site)SGFRKMAFPSGKVEGCMVQVTCGTTTLNGLWLDDV VYCPRHVICTSEDMLNPNYEDLLIRKSNHNFLVQAGNVQLRVIGHSMQNCVLKLKVDTANPKTPKYKFVRIQPGQTFSVLACYNGSPSGVYQCAMRPNFTIKGSFLNGSCGSVGFNIDYDCVSFCYMHHMELPTGVHAGTDLEGNFYGPFVDRQTAQAAGTDTTITVNVLAWLYAAVINGDRWFLNRFTTTLNDFNLVAMKYNYEPLTQDHVDILGPLSAQTGIAVLDMCASLKELLQNGMNGRTILGSALLEDEFTPFDVVRQCSGVTFQ(PreScission_protease_cleavage_site)GPHHHHHH* (* is stop). The corresponding gene is synthesized by Genscript, USA and cloned into pET28a(+) bacterial vector by using NdeI and BamHI restriction cleavage sites at 5’ and 3’ ends respectively. N-terminal canonical SARS-CoV-2 Mpro autocleavage cut site is indicated by green and purple which generates the native N-terminus. C-terminus has the PreScision™ restriction site shown in red which is used to generate the native C-terminus after Ni-NTA hexa-histidine affinity purification chromatography. In-frame hexa-histidine tag and stop codon is shown in blue color.

#### Construct-2 with Modified N-terminus and Native C-terminus

To eliminate the critical N-terminus crystal contact to obtain a new apo crystal form we designed the modified Mpro construct with the following amino acid sequence inserted in th *E. coli* vector pET28a(+) MGSSHHHHHHSSGLVPR(thrombin_cleavage_site)GSHMSGFRKMAFPSGKVEGCMVQVTCG TTTLNGLWLDDVVYCPRHVICTSEDMLNPNYEDLLIRKSNHNFLVQAGNVQLRVIGHSMQNCVLKLKVDTANPKTPKYKFVRIQPGQTFSVLACYNGSPSGVYQCAMRPNFTIKGSFLNGSCGSVGFNIDYDCVSFCYMHHMELPTGVHAGTDLEGNFYGPFVDRQTAQAAGTDTTITVNVLAWLYAAVINGDRWFLNRFTTTLNDFNLVAMKYNYEPLTQDHVDILGPLSAQTGIAVLDMCASLKELLQNGMNGRTILGSALLEDEFTPFDVVRQCSGVTFQ* (* is stop). The gene is synthesized by Genscript, USA and cloned into pET28a(+) bacterial overexpression vector by using NdeI and BamHI restriction sites at 5’ and 3’ ends respectively. N-terminal hexa-histidine tag (labeled in purple) and modified SARS-CoV-2 Mpro thrombin cleavage site which is part of the pET28a(+) vector indicated blue sequence which generates the modified N-terminus with four extra residues as follows (GSHM) shown in blue and underlined. C-terminus has the in-frame stop codon that is used to generate the native C-terminus by ribosome during bacterial overexpression shown in green asterisk.

#### Protein expression

Both constructs were transformed into *E. Coli* BL21 Rosetta-2 strain. 12 liters of six independent bacterial cell cultures containing target Mpro protein genes were grown in either regular LB-Miller media or Terrific Broth (TB) supplemented with 35 μg/ml chloramphenicol and 50 μl/ml kanamycin at 37°C. Cultures are incubated by using New Brunswick Innova 4430R shaker at 110 rpm until they reach OD600 about 0.8-1.2 for each culture. Recombinant protein expression was induced by IPTG with a final concentration of 0.4 mM. Incubation for protein production was performed at 18°C for minimum of 24 hours and maximum of 7 days. Cells were harvested at 4°C by using Beckman Allegra 15R desktop centrifuge at 3500 rpm for 20 minutes. Protein expression was confirmed by precast TGX-mini protean gradient SDS-PAGE from BioRad.

### Protein purification

Standard chromatography purification methods were applied to both constructs with slight modifications as described below. Soluble Mpro proteins were purified by first dissolving the bacterial cells in the lysis buffer containing 50 mM Tris pH 7.5, 300 mM NaCl, 5% v/v Glycerol supplemented with 0.01% Triton X-100 followed by sonication (Branson W250 sonifier, USA). After sonication step, cell lysate was centrifuged by using The Beckman Optima™ L-80 XP Ultracentrifuge at 40000 rpm for 30 minutes at 4°C by using Ti45 rotor (Beckman, USA). After ultracentrifugation the pellet which contains membranes and insoluble debris was discarded and clear supernatant applied to nickel affinity chromatography by using a Ni-NTA agarose resin (QIAGEN, USA). To purify the Mpro protein, first the chromatography column was equilibrated by flowing 3 column volume of the loading buffer containing 20 mM Tris-HAc pH 7.5, 5 mM Imidazole, 150 mM NaCl. After equilibration, the supernatant containing the overexpressed Mpro protein was loaded into the Ni-NTA agarose column at 2 ml/minute flow rate. Unbound proteins were removed by washing with 5 column volumes of the loading buffer to clear the non-specific binding. After washing, hexa-histidine tagged Mpro proteins were eluted from the column with the elution buffer containing 20 mM Tris-HAc pH 7.5, 150 mM NaCl, 250 mM Imidazole in 35 ml of total volume. After elution, purified protein was placed in a 3 kDa cut off dialysis membrane and dialyzed against the buffer containing 20 mM Tris-HAc pH 7.5, 150 mM NaCl overnight to get rid of the excess Imidazole. After the dialysis step we applied 1:100 stoichiometric molar ratio 3C protease (PreScission protease, GenScript, USA) to cleave the C-terminal hexa-histidine tag of construct-1 with native N- & C-terminals. For construct-2 with modified N-terminus we used thrombin protease (Sigma, USA) to get rid of the N-terminal hexa-histidine tag. Both PreCision and thrombin cleavage have been performed overnight at 4°C. In the final purification step, to remove the cleaved hexa-histidine tag and other non-specific binding proteins we applied the solution to reverse Ni-NTA chromatography and collected the unbound fractions containing the untagged Mpro protein. The pure Mpro concentrated by ultrafiltration columns from Millipore to a final concentration of 25 mg/ml and added 1 mM final concentration of DTT and stored at −80°C until crystallization trials.

### Crystallization of Mpro protein for SFX crystallography at XFEL

For initial crystallization screening, we employed sitting-drop microbatch under oil screening method by using 72 well Terasaki crystallization plates (Greiner-Bio, Germany). Purified Mpro protein at 25 mg/ml mixed with 1:1 volumetric ratio with ~3500 commercially available sparse matrix crystallization screening conditions. The sitting drop solutions were then covered with 20 μl of 100% paraffin oil (Tekkim Kimya, Turkey). All the crystallization experiments were performed at ambient-temperature. For our native construct-1 we were able to obtain multiple hit conditions and among them the best crystals were obtained at *Pact Premier*™ crystallization screen 1 condition #39 from Molecular Dimensions, UK. The best crystallization condition has contained 100 mM MMT buffer pH 6.0 and 25 % w/v PEG 1500 [MMT buffer; DL-Malic acid, 4-Morpholine Ethane Sulfonicacid (MES) monohydrate, 2-Amino-2-(hydroxymethyl)-1,3-propanediol (TRIS)-HCl]. For the modified construct only one crystallization condition yield the macrocrystals. After multiple optimization of the seeding protocol by using crystals obtained by microbatch under oil, we scaled up the batch crystallization volume to total of 14 ml for native construct-1 and total volume of 50 ml for modified construct-2. Microcrystals 1-5 × 5-10 × 10-20 μm^3^ in size were passed through 100 micron plastic mesh filters (Millipore, USA) in the same mother liquor composition to eliminate the large single crystals and other impurities before the data collection. Crystal concentration was approximated to be 10^10^-10^11^ particles per ml based on light microscopy. Due to COVID19 travel restrictions none of the initial crystals or the batched crystalline slurry were able to be pretested for their diffraction quality before the scheduled XFEL beamtime.

### Transport of Mpro microcrystal for SFX studies at MFX instrument at the LCLS

1.9 ml total volume of crystal slurry was transferred to 2 ml screw top cryovial (Wuxi NEST biotechnology, China cat#607001). To absorb the mechanical shocks during transport from Istanbul to Menlo Park, CA these vials were wrapped loosely by Kimwipes (Kimberly-Clark, USA) and placed in 20 ml screw top glass vials and tightly closed to provide insulation during transport via air. The vials were wrapped with excess amounts of cotton (Ipek, Turkey) and placed in a Ziploc™ bag (SC Johnson, USA) to provide both added layer of insulation and mechanical shock absorption. The Ziploc™ bags were placed in a styrofoam box that was padded with ~1 kg of cotton to provide more insulation and mechanical shock absorption during the transport. The styrofoam box was sealed and wrapped with an additional layer of 1 cm thick loose cotton layer and duck taped all around to further insulate the delicate Mpro crystals during ambient-temperature transport. All these packing materials and techniques provided us with crystals diffracting to 1.9 Å - 2.1 Å resolution as described below.

### MESH Sample injection for MPro crystals

The 1.6 ml sample reservoir was loaded with Mpro crystal slurry in their unaltered mother liquor as described above. We used standard Microfluidic Electrokinetic Sample Holder (MESH) (Sierra et al., 2012; Sierra et al., 2016) injector for our sample injection. The sample capillary was a 200 μm ID × 360 μm OD × 1.0 m long fused silica capillary. The applied voltage on the sample liquid was typically 2500-3000 V, and the counter electrode was grounded. The sample ran typically between 2.5 and 8 μl/min.

### Data Collection and Analysis for SFX studies at LCLS

The SFX experiments with native Mpro microcrystals were carried out at the LCLS beamtime ID: mfx17318 at the SLAC National Accelerator Laboratory (Menlo Park, CA). The LCLS X-ray beam with a vertically polarized pulse with duration of 30 fs was focused using compound refractive beryllium lenses to a beam size of ~6 × 6 μm full width at half maximum (FWHM) at a pulse energy of 0.8 mJ, a photon energy of 9.8 keV (1.25 Å) and a repetition rate of 120 Hz. OM monitor (Mariani et al., 2016) and *Psocake* (Damiani et al., 2016, Thayer et al., 2017) were used to monitor crystal hit rates, analyze the gain switching modes and determine the initial diffraction geometry of the new ePix10k2M detector (van Driel et al., 2020). A total of 1,163,413 detector frames were collected in 2h47m7s continuously with the new from native (construct-1) Mpro microcrystals. A total of 686,808 detector frames were collected in 1h36m20s continuously with the new ePix10k2M Pixel Array Detector from modified (construct-2) Mpro microcrystals. The total beamtime needed for native (construct-1) and modified (construct-2) datasets were 2h59m17s and 1h50m59s respectively, which shows the efficiency of the MFX beamline installed with the new ePix10k 2M detector and robust injector system, as due to lack of blockages no dead time was accumulated. Individual diffraction pattern hits were defined as frames containing more than 30 Bragg peaks with a minimum signal-to-noise ratio larger than 4.5, which were a total of 208,839 and 214,355 images for native and modified respectively. The detector distance was set to 118 mm, with an achievable resolution of 2.1 Å at the edge of the detector (1.64 Å in the corner). An example diffraction pattern is shown in **fig. S27**.

### Data processing; hit finding, indexing and scaling

The diffraction patterns were collected at the MFX instrument at the LCLS using the ePix10k2M detector (Van Driel et al., 2020). The raw data images were subjected to detector corrections with cheetah (Barty et al., 2014), as well as for hit finding based on Bragg reflections. The hitfinding parameters for all datasets classifying a hit were as follows (using peakfinder8): a minimum pixel count of 2 above an adc-threshold of 500 with a minimum signal to noise ratio of 7 was considered a peak, and an image containing at least 20 peaks was classified as a crystal hit. The crystal hits were then indexed using the software package *CrystFEL* (White et al., 2012; White et al., 2019) version 9.0 (White et al., 2020) using the peaks found by *CHEETAH*. Indexing was attempted using the indexing algorithms from *XGANDALF* (Gevorkov et al., 2019), *DIRAX* (Duisenberg et al., 1992), *MOSFLM* (Powell et al., 2013) and *XDS* (Kabsch et al., 2010), in this order. After an approximate cell was found, the data was indexed using cell axis tolerances of 5 Å and angle tolerances of 5° (--tolerance option in *CrystFEL*). The integration radii were set to 2, 3, 5 and the “multi” option was switched on to enable indexing of multiple crystal lattices in a single image. The indexed reflections were subsequently integrated and merged using partialator (White et al., 2016) applying the unity model over 3 iterations and the max-ADU set to 7500. The complete reflection intensity list from *CrystFEL* was then scaled and cut using the *TRUNCATE* program from the *CCP4* suite (Winn et al., 2011) prior to further processing.

For the native Mpro protein crystals the final set of indexed patterns, containing 168,655 frames (80.7% indexing rate), was merged into a final dataset (Overall CC* = 0.999; 1.8 Å cutoff) for further analysis (C121, unit cell: a = 114.0 Å, b = 53.5 Å, c = 45.0 Å; α = 90°, β = 102°, γ = 90°). The final resolution cutoff was estimated to be 1.9 Å using a combination of CC* (Karplus & Diederichs, 2012) and other refinement parameters. The final dataset had overall Rsplit = 6.31%, and CC* = 0.865 in the highest resolution shell. For the N-terminally modified Mpro protein crystals the final set of indexed patterns, containing 157,976 frames (73.6% indexing rate), was merged into a final dataset (Overall CC* = 0.999; 2.1 Å cutoff) for further analysis (P2_1_2_1_2_1_, unit cell: a = 69.2 Å, b = 104.3 Å, c = 105.6 Å; α = β = γ = 90°). The final resolution cutoff was estimated to be 2.1 Å using a combination of CC* and other refinement parameters. The final dataset had overall Rsplit = 5.91%, and CC* = 0.678 in the highest resolution shell.

### Structure determination and refinement of Apo Mpro structures

We determined two ambient-temperature Mpro structures by using two crystal forms in space group C121 and P2_1_2_1_2_1_ structures using the automated molecular replacement program *PHASER* (McCoy et al., 2007) implemented in *PHENIX* (Adams et al., 2010) with the previously published ambient-temperature structure as a search model (PDB ID: 6WQF) (Kneller et al., 2020). This choice of starting search model minimized experimental temperature variations between the two structures. Coordinates of the 6WQF were used for initial rigid body refinement with the *PHENIX* software package. After simulated-annealing refinement, individual coordinates and TLS parameters were refined. We also performed composite omit map refinement implemented in *PHENIX* to identify potential positions of altered side chains and water molecules were checked in program *COOT* (Emsley & Cowtan, 2004), and positions with strong difference density were retained. Water molecules located outside of significant electron density were manually removed. The Ramachandran statistics for native monoclinic Mpro structure (PDB ID: 7CWB) (most favored / additionally allowed / disallowed) are 96.7 / 3.0 / 0.3 % respectively. Ramachandran statistics for orthorhombic Mpro structure (PDB ID: 7CWC) (most favored / additionally allowed / disallowed) are 96.5 / 2.4 / 0.1 % respectively. The structure refinement statistics are summarized in **Supplementary Table S1**. Structure alignments were performed using the alignment algorithm of *PyMOL* (www.schrodinger.com/pymol) with the default 2σ rejection criterion and five iterative alignment cycles. All X-ray crystal structure figures were generated with *PyMOL*.

### Temperature factor analysis and generation of ellipsoids

The two ambient-temperature Mpro in space group C121 and P2_1_2_1_2_1_ were examined to generate ellipsoid structures based on b-factor with *PyMOL* and these two structures were compared with the Mpro structures at 100 K (PDB ID: 6XKH) to provide better understanding on the flexibility of atoms, side chains and domains. The all ellipsoid structures were colored with rainbow selection on *PyMOL*.

### Molecular Modeling Studies

We have used different crystal structures of Mpro available in literature as well as the obtained crystal structures in this study (PDB IDs: 7CWB and 7CWC) as target structures for molecular docking and MD simulations. The biologically-relevant dimeric form of 7CWB is generated by application of a symmetry operator. As the crystal structures in this study were obtained at ambient-temperature, for comparison another ambient-temperature structure of Mpro (PDB ID: 6WQF) in apo form was also selected as target structure. Another apo form structure of Mpro (PDB ID: 6Y2E) in dimeric form was also chosen for comparison. Additionally, Mpro structure bound to a non-covalent inhibitor (PDB ID: 6W63) in both monomeric and dimeric forms was utilized as target structure. For ligands, we have considered three compounds that have shown promising inhibitory activity based on the high-throughput screening of over 10,000 compounds by Jin et al., namely; Ebselen (IC50 = 0.67 ± 0.09 μM), Tideglusib (IC50 = 1.55 ± 0.30 μM) and Carmofur (IC50 = 1.82 ± 0.06 μM) (Jin et al., 2020).

All the target structures considered in this study were firstly prepared using Protein Preparation module of Maestro modeling program in which missing atoms were added, water molecules not in the vicinity of co-crystallized ligands were removed and bond orders were assigned (Madhavi Sastry et al., 2013). The protonation states of amino acids at physiological pH were adjusted using PROPKA (Bas et al., 2008) to optimize the hydrogen binding and charge interactions. As a final step of preparation, a restrained minimization was performed with OPLS3e force field parameters (Harder et al., 2016).

The structures of three compounds were taken from PubChem; Ebselen (PubChem ID, 3194), Tideglusib (PubChem ID: 11313622) and Carmofur (PubChem ID: 2577). The compounds also needed preparation hence, LigPrep module (Schrödinger Release 2018-4, 2018) of Maestro modeling program was employed with OPLS3e force field parameters (Harder et al., 2016). The ionization states of the molecules were predicted by Epik module (Shelley et al., 2007) at physiological pH of 7.4.

The prepared target protein and ligand structures were used for molecular docking studies. We have employed a grid-based docking method, Induced Fit Docking (IFD) protocol of Maestro (Sherman et al., 2006a,b) which uses Glide (Friesner et al., 2004; Halgren et al., 2004; Friesner et al., 2006) and Prime (Jacobson et al., 2004) to induce adjustments in receptor structures with flexible ligand sampling options. For target structures in apo form (structures with PDB IDs: 7CWB, 7CWC, 6WQF and 6Y2E), binding sites for docking studies were defined by centering grids at the centroid of a set of residues, namely His41, Cys145 and Glu166. On the other hand, for target structures in holo form (structures with PDB ID: 6W63 both monomeric and dimeric form) binding sites were defined by centering the grids at the centroid of the co-crystallized ligand molecule. In the dimeric form of Mpro, only one chain was considered for docking studies. IFD protocol involves subsequent phases, including (i) initial docking of the compounds with the rigid receptor in which 20 poses per ligand are retained; (ii) refining the residues (within 5 Å of the ligand) in complex using Prime module (Jacobson et al., 2004); and (iii) redocking of each protein/ligand complex structure within 30 kcal/mol of the lowest-energy structure and within the top 20 structures overall. During Glide redocking, standard precision (SP) option was chosen. The docking poses were scored and ranked based on GlideScore and poses with the lowest scores, i.e. top-docking poses were selected for further studies at each target protein.

The selected docking poses at each considered target structure of Mpro with the three compounds were subjected to MD studies. The apo form structures obtained in this study were also subjected to MD simulations. For comparison reasons, MD simulations were also performed for the holo form structure (PDB ID: 6W63) with its co-crystallized ligand, X77. The target protein-ligand complexes were placed in simulation boxes with orthorhombic shape in which box sizes were calculated based on buffer distance of 10.0 Å along all three dimensions and solvated with explicit water molecules of SPC (Berendsen et al., 1987) model. The simulation systems were neutralized by the addition of counter ions (Na+ or Cl-depending on the charge of the systems) and 0.15 M NaCl solution was added to adjust concentration of the solvent systems. All atom MD simulations package Desmond (Bowers et al., 2006) was employed. Proceeding the production MD simulations, the systems were equilibrated using relaxation protocols of Desmond package in which a series of minimizations and short MD simulations which are performed with small time-steps at lower temperature and restrains on the nonhydrogen solute atoms in the initial stages and slowly time-steps are increased as well as simulation temperature and restrains on solute atoms are released. The production simulations were performed under constant pressure and temperature conditions, i.e. *NPT* ensemble. Temperature was set as 310 K while being controlled by Nose–Hoover thermostat (Nosé, 1984; Hoover, 1985). The pressure was set as atmospheric pressure of 1.01325 bar with isotropic pressure coupling and controlled by Martyna–Tobias–Klein barostat (Martyna et al., 1994). Smooth particle mesh Ewald method (Essmann et al., 1995) was utilized to calculate long range electrostatic interactions with periodic boundary conditions (PBC). For short range electrostatics and Lennard-Jones interactions, the cut-off distance was set as 9.0 Å. The multi-step integrator RESPA was employed in which the time steps were varied for interaction types as followed in fs for: *bonded*, 2.0; *near* 2.0 and *far* 6.0.

Principal components analysis (PCA), a statistical data processing method, were performed to reduce the large-dimensional data by extracting large amplitude motions onto collective sets. A covariance matrix were generated from MD trajectory data for backbone atoms of protein structures as follow

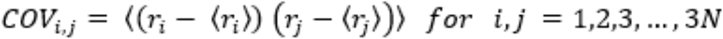

Here, *i* and *j* represent the backbone atom number, i.e. residue numbers of proteins while *N* is the number of backbone atoms considered in analysis. The Cartesian coordinates of atoms are denoted by and for *i*th and *j*th atom, respectively with and representing the time-averaged values over MD simulations. By diagonalization of covariance matrix, a collection of eigenvectors and corresponding eigenvalues were obtained. The eigenvectors of the diagonalized matrix are referred as principal components (PCs) and constitute a linear basis set that matches the distribution of observed structures. The corresponding eigenvalues of the diagonalized matrix display the variance of the distribution along each PCs. In this study, we have utilized the Bio3d package (Grant et al., 2006; Yao et al., 2013), a platform independent R package to perform PCA for considered simulation systems. The trajectories obtained from independent MD simulations were concatenated and frames were aligned with respect to the initial (reference) frame before PCA.

The correlation of atomic displacements is evaluated by cross-correlation analysis to appreciate the coupling of motions. The magnitudes of all pairwise cross-correlation coefficients were investigated to assess the extent of atomic displacement correlations for each simulation system in principal component space. The normalized covariance matrix of atomic fluctuations was calculated as

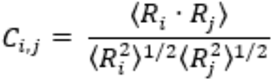

where and are the displacements of residues *i* and *j*, i.e., mean square atomic fluctuations. The values of varies between −1 to 1 with representing completely correlated motions (same period and same phase), representing completely anticorrelated motions (same period and opposite phase) while 0 value indicates motions are uncorrelated. (Ichiye et al., 1991; McCammon et al., 1988). Bio3d package (Grant et al., 2006; Yao et al., 2013) in R environment was employed to generate atom-wise crosscorrelations of motions observed in PCs 1 to 3 and dynamical cross-correlation map, or DCCM were generated and displayed as a graphical representation of cross-correlation coefficients.

### Interface Analysis

Interface analysis of crystal structures and MD trajectories were carried out with the GetContacts python scripts (https://getcontacts.github.io/). Two different approaches were followed, in the first one only hydrogen bonds at the dimerization interface were taken into account, and in the second approach all possible interactions, namely; salt bridges, pi-cation, pi-pi stacking, T-stacking, van der Waals (vdW), and hydrogen bonds were calculated. If the distance between the acceptor and the donor atoms is <3.5 Å and the angle <70 ° hydrogen bond is defined between atom groups. Salt bridges were defined between atoms of negatively charged [ASP (OD1, OD2) and GLU (OE1, OE2)] and positively charged [LYS (NZ) and ARG (NH1, NH2)], where distances were <4.0 Å. T-stacking, pi-cation, and pi-stacking distance criteria were 5.0, 6.0, and 7.0 Å, respectively. Hydrophobic and vdW interactions were calculated based on atom R (radii), if the distance between atoms is less than the sum of R of atom A, R of atom B, and 0.5 Å.

One frame and trajectory-based calculations were performed. One frame calculations were applied for the crystal structures, and 1 means the corresponding residues are in contact, and 0 means no interaction. However, in trajectory-based calculations, we used all available 2000 frames from our MD trajectories. For the interaction frequencies, we applied a 0.6 threshold to only take into account the contacts that are occurred at least 60% of the simulations.

## Supplementary Figures and Tables

**Fig. S1.**
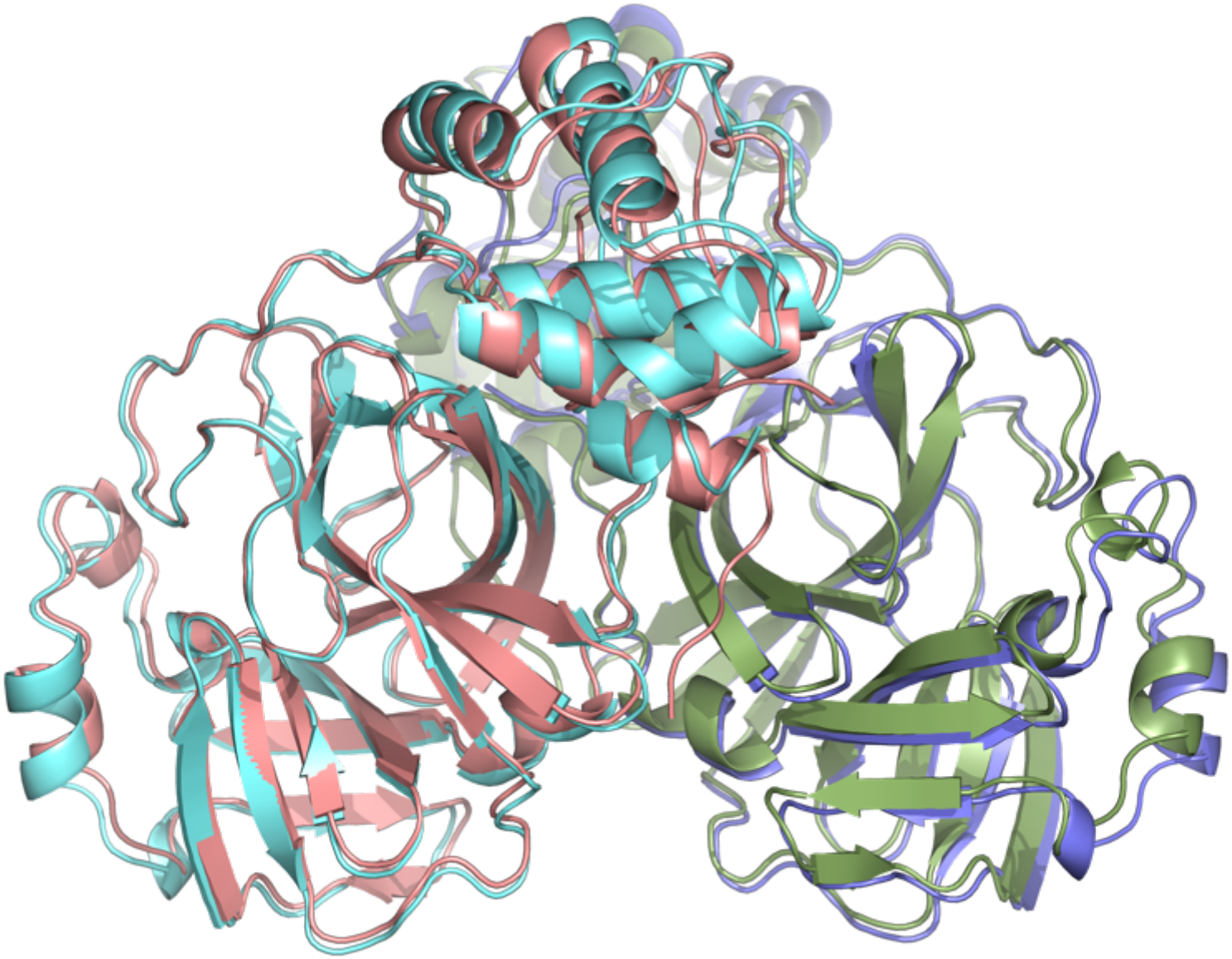
Superposition of two crystal forms. Native Mpro in space group C121 colored in darksalmon and its symmetry mate in green. Modified Mpro in space group P2_1_2_1_2_1_ is colored in palecyan and light blue. Two crystal structures align with an overall RMSD of 1.00 Å.

**Fig. S2.**
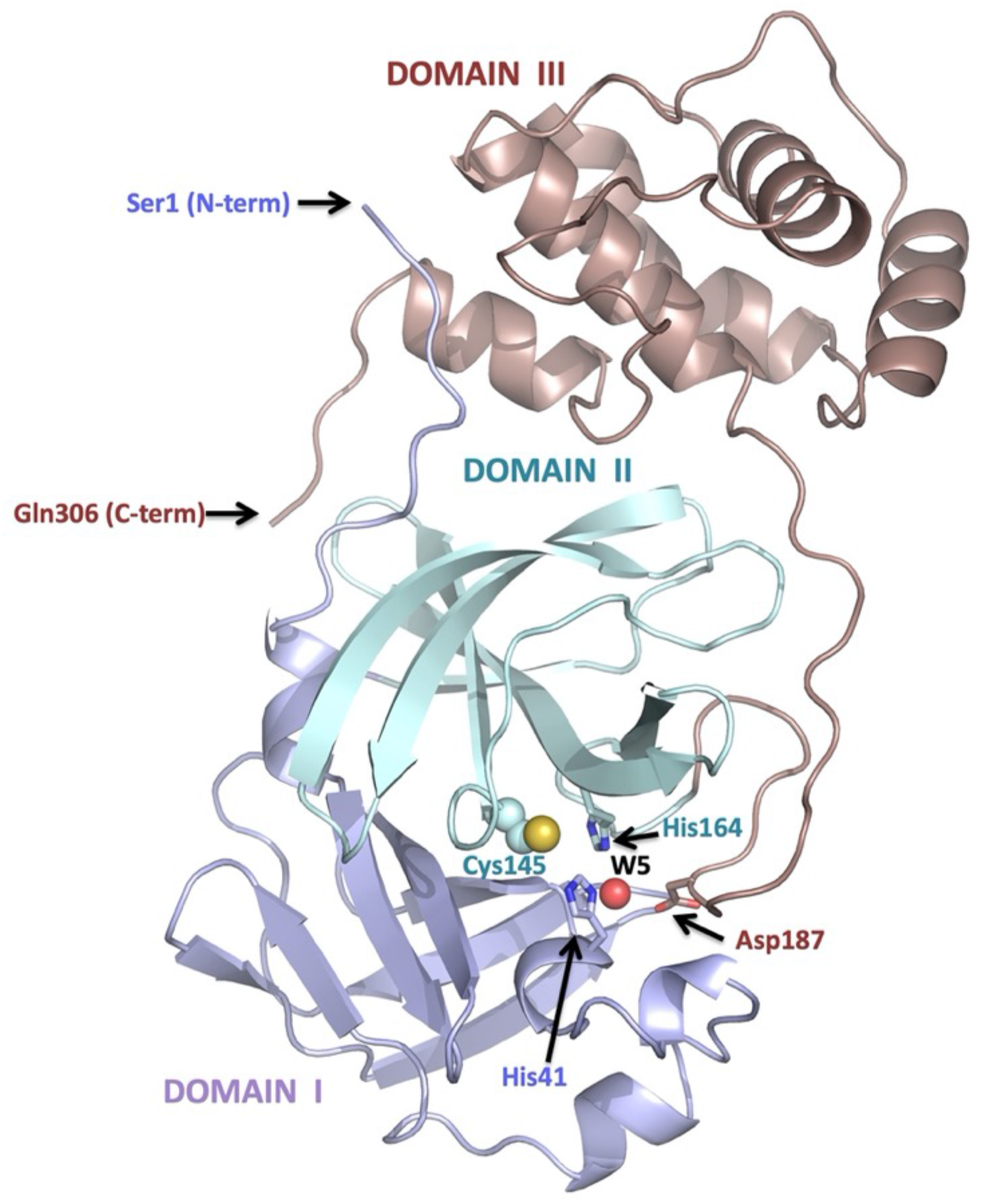
Three major domains of SARS-CoV-2 Mpro. Domain I is colored in light blue, domain II is colored in pale cyan and domain III is colored in dark salmon. Active site formed by residues from all three subdomains such as critical Cys145 is shown as spheres, His41, His164 and Asp187 shown in sticks. Red sphere labeled as W5 represents the water molecule in the binding pocket.

**Fig. S3.**
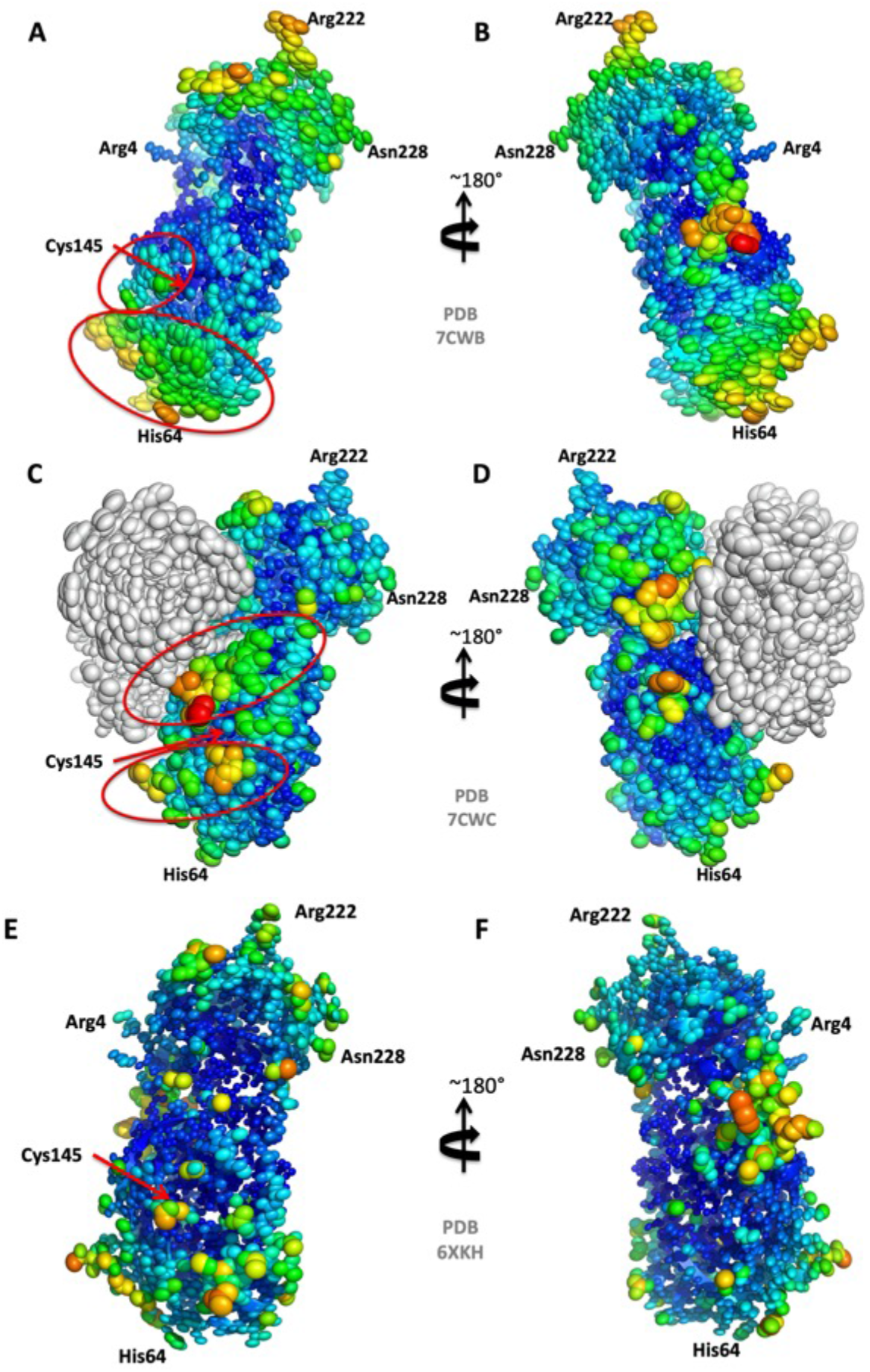
Representation of ellipsoids for SFX and cryogenic apo Mpro crystal structures (PDB IDs: 7CWB, 7CWC and 6XKH) based on B-factor. **A)** Mpro structure in space group C121 at 1.9 Å is examined to indicate the magnitude of thermal vibration. **B)** The 180 degrees rotated view of panel a) around y axis **C)** The dimer crystal form of Mpro structure in space group P2_1_2_1_2_1_ at 2.1 Å is examined and chain A is colored by rainbow selection while chain B is colored by grey. **D)** The 180 degrees rotated panel c) around y axis. **E)** Mpro structure at 1.28 Å (PDB ID: 6XKH) is examined to compare our ambient-temperature crystal structures with the X-ray structure at 100 K. **F)** The 180 degrees rotated view of panel a) around y axis. All the ellipsoid structures are produced with PyMOL (www.pymol.org).

**Fig. S4.1.**
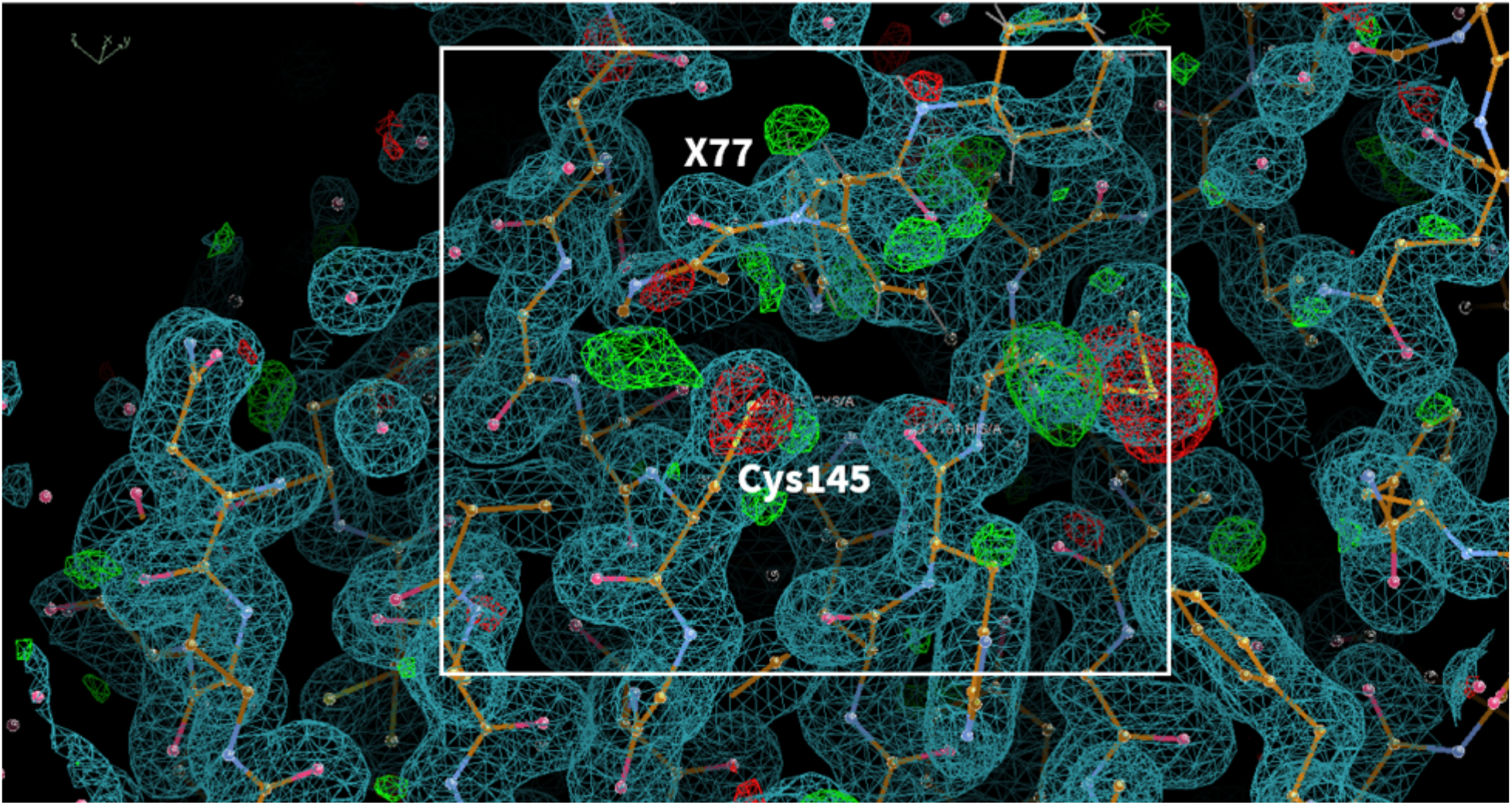
(non-covalent inhibitor) **PDB ID:** 6W79 **Ligand:** X77 **Resolution:** 1.46 Å **Author(s):** Mesecar et al. **doi:** 10.2210/pdb6W79/pdb **Deposited:** 2020 March 18 **Released:** 2020 August 26

**Fig. S4.2.**
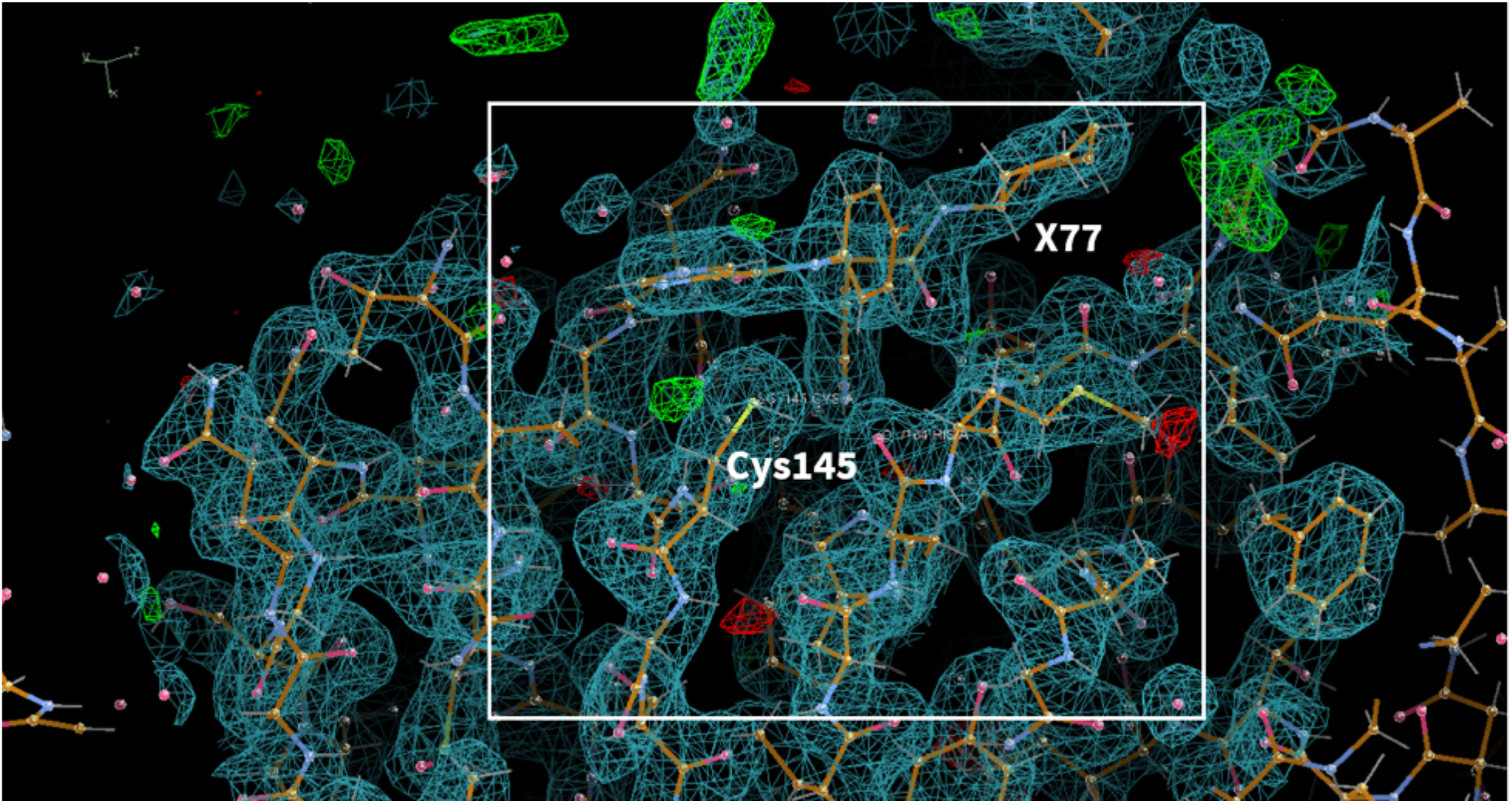
(non-covalent inhibitor) **PDB ID:** 6W63 **Ligand:** X77 **Resolution:** 1.46 Å, **Author(s):** Mesecar et. al. **doi**: 10.2210/pdb6W63/pdb **Deposited:** 2020 March 16 **Released:** 2020 March 25

**Fig. S4.3.**
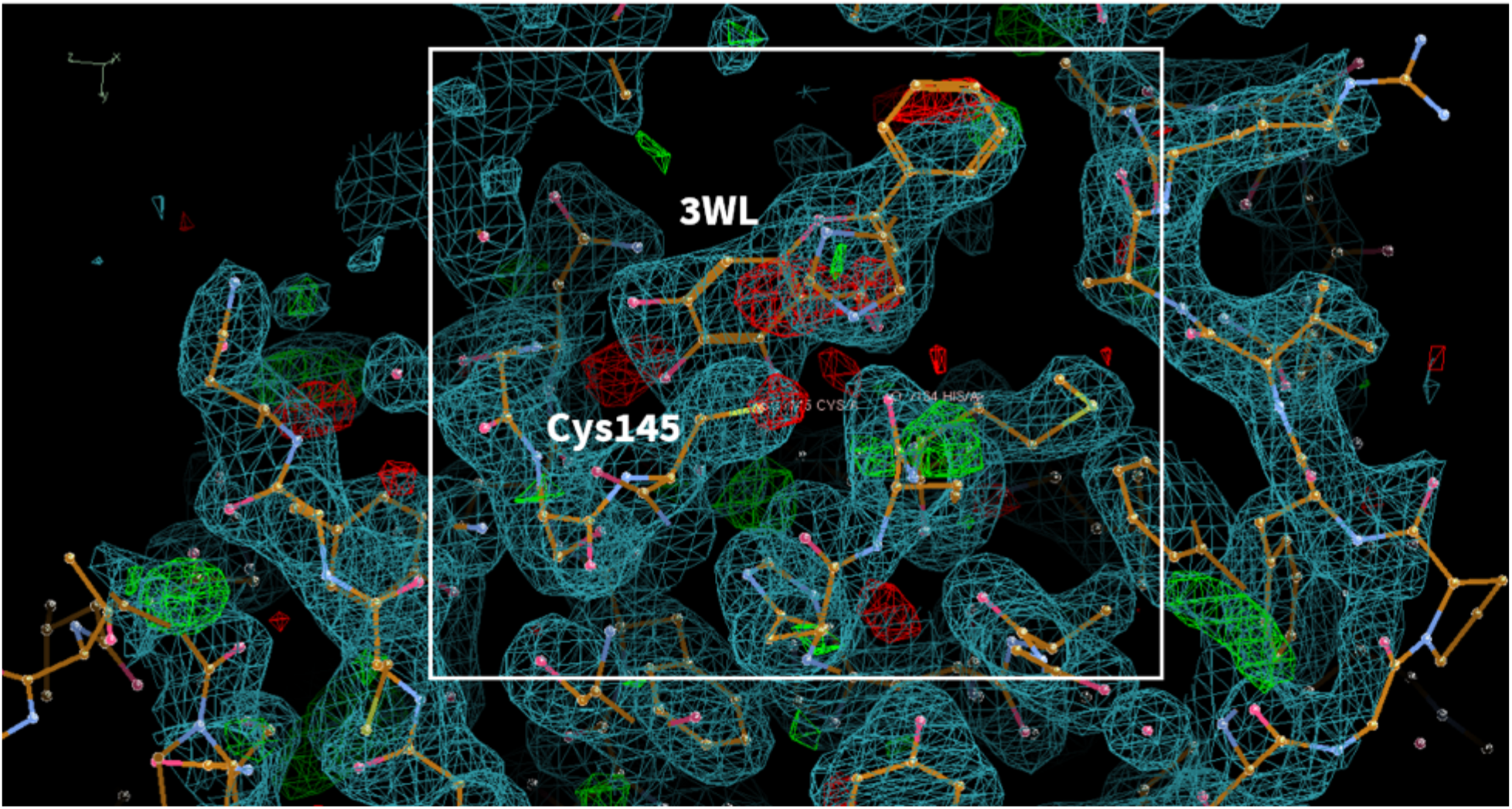
(non-covalent inhibitor) **PDB ID:** 6M2N **Ligand:** 3WL **Resolution:** 2.20 Å **Author(s):** Su et al. **doi:** 10.2210/pdb6M2N/pdb **Deposited:** 2020 February 28 **Released:** 2020 April 15

**Fig. S4.4.**
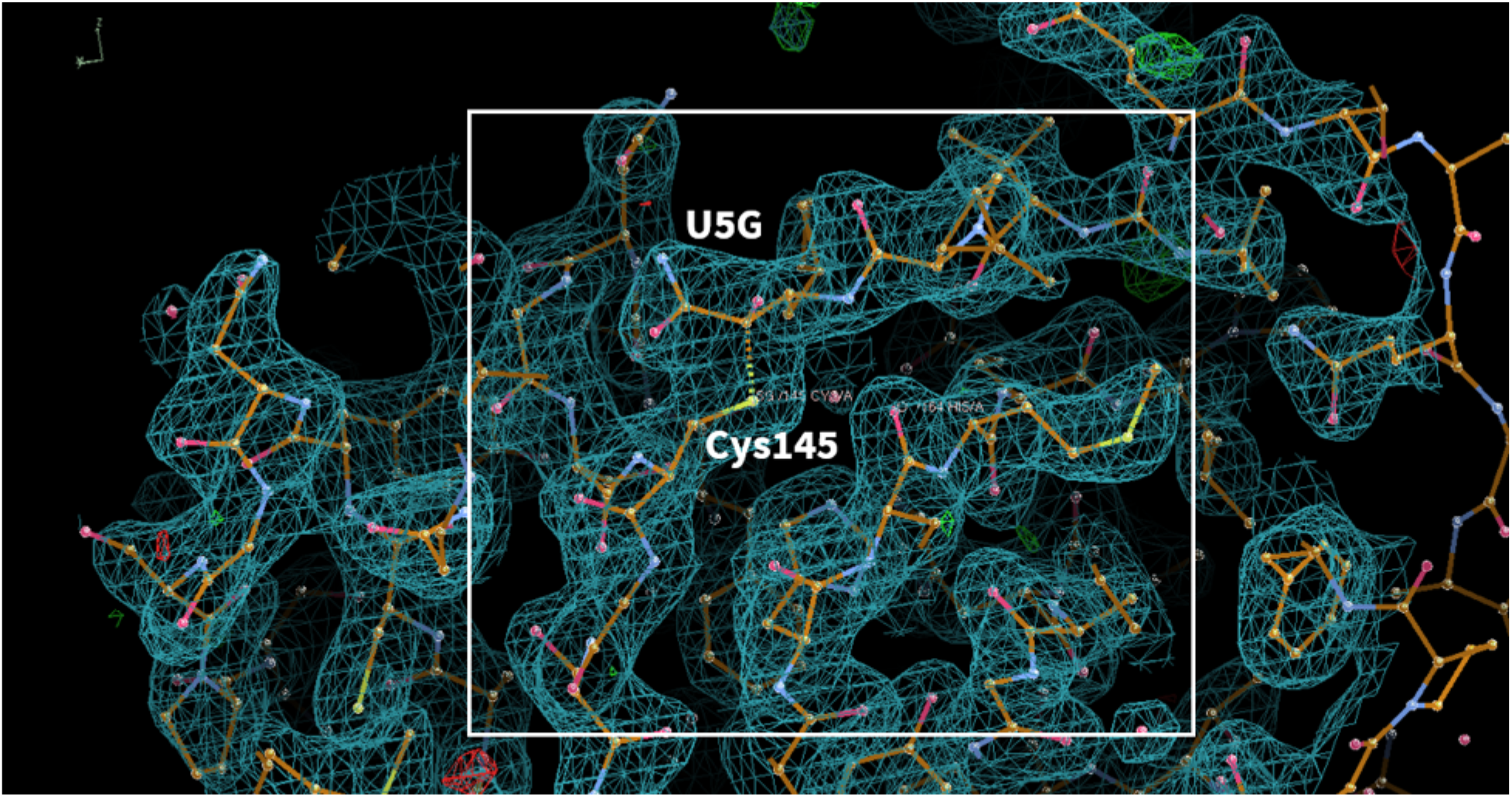
(covalent inhibitor) **PDB ID:** 6ZRU **Ligand:** Boceprevir(U5G), **Resolution** 2.10 Å **Author(s)**: Oerlemans et al. **doi:** 10.2210/pdb6ZRU/pdb **Deposited:** 2020 July 14 **Released:** 2020 August 12

**Fig. S4.5.**
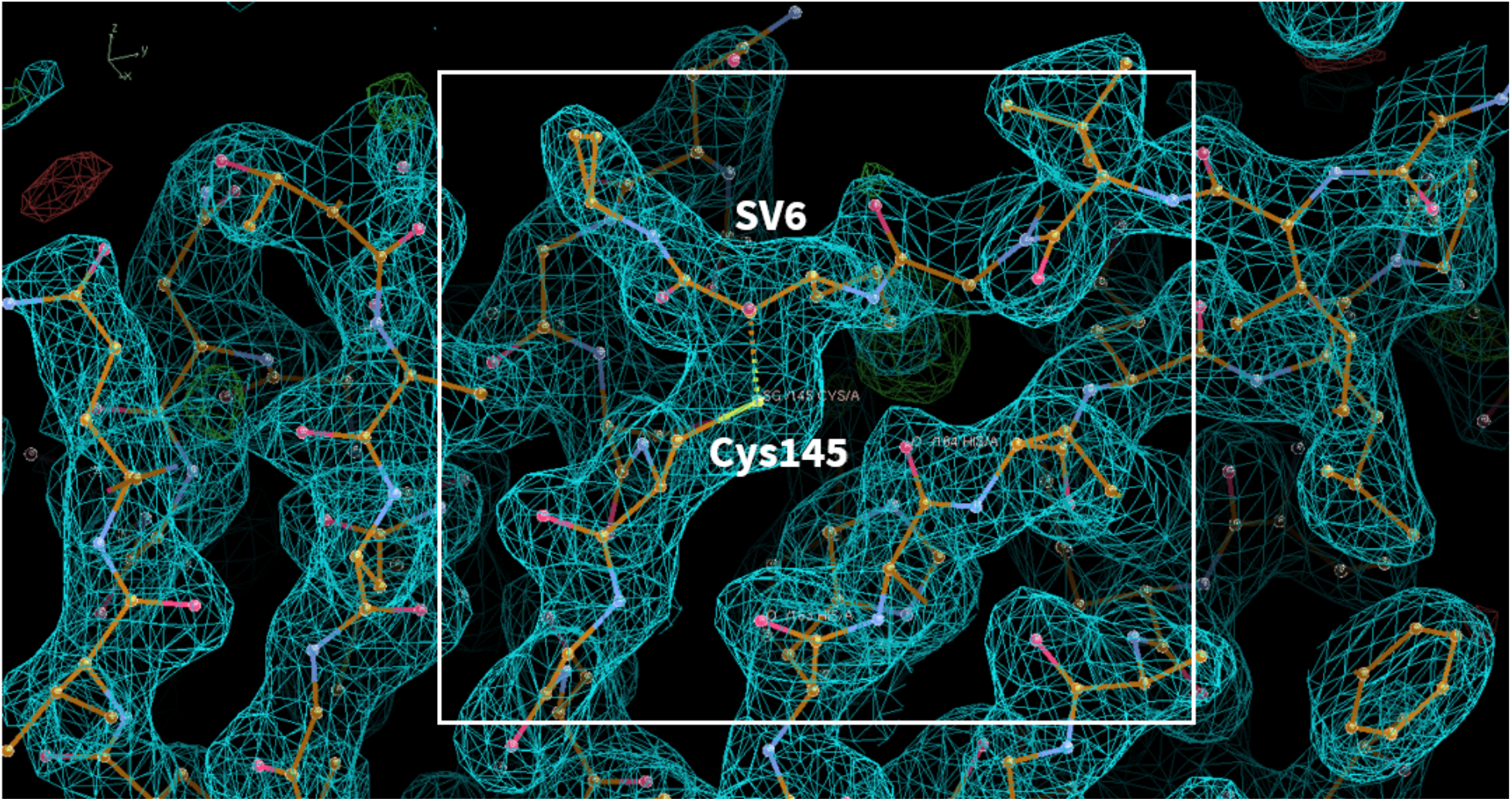
(covalent inhibitor) **PDB ID:** 6ZRT **Ligand:** Telaprevir(SV6) **Resolution**: 2.10 Å **doi:** 10.2210/pdb6ZRT/pdb **Author(s)**: Oerlemans et al. **Deposited:** 2020 July 14 **Released:** 2020 August 12

**Fig. S4.6.**
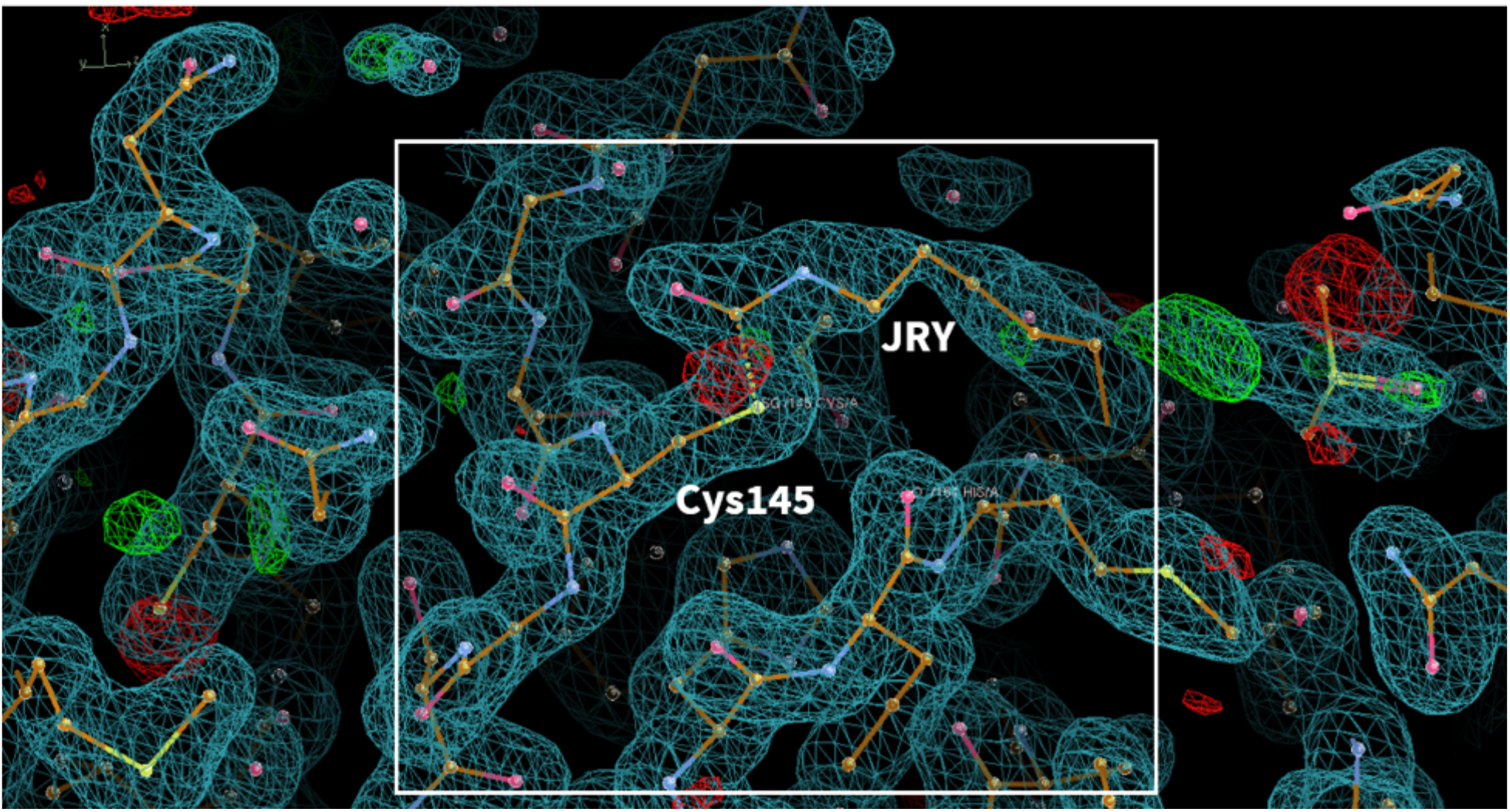
(covalent inhibitor) **PDB ID:** 7BUY **Ligand:** Carmofur(JRY) **Resolution**: 1.60 Å **doi:** 10.2210/pdb7BUY/pdb **Author(s)**: Jin et al. **Deposited:** 2020 April 08 **Released:** 2020 April 29

**Fig. S4.7.**
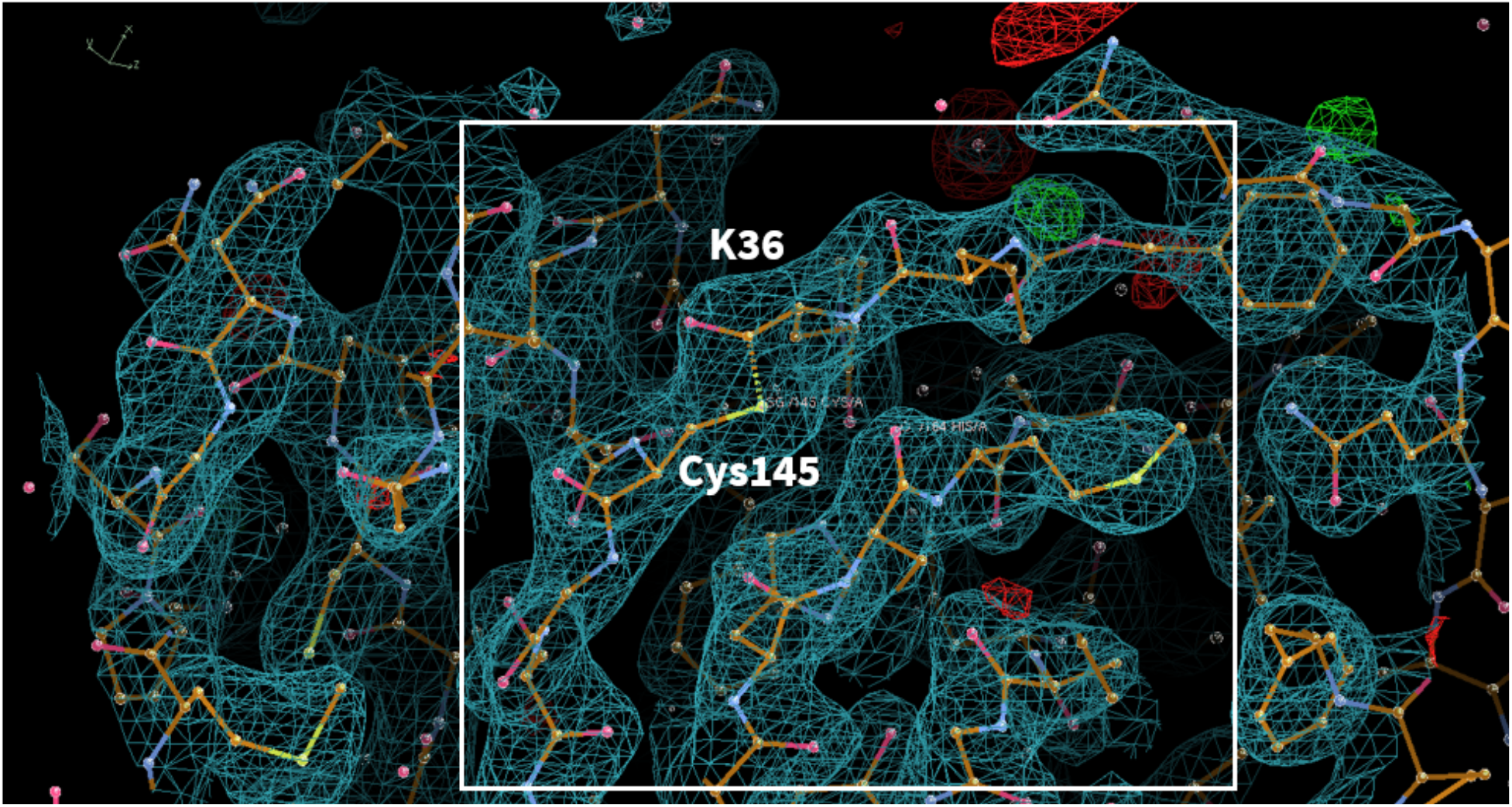
(covalent inhibitor) **PDB ID:** 6WTT **Ligand:** GC-376(K36) **Resolution**: 2.15 Å **doi:** 10.1101/2020.04.20.051581 **Author(s)**: Ma et al. **Deposited:** 2020 May 03 **Released:** 2020 May 20

**Fig. S4.8.**
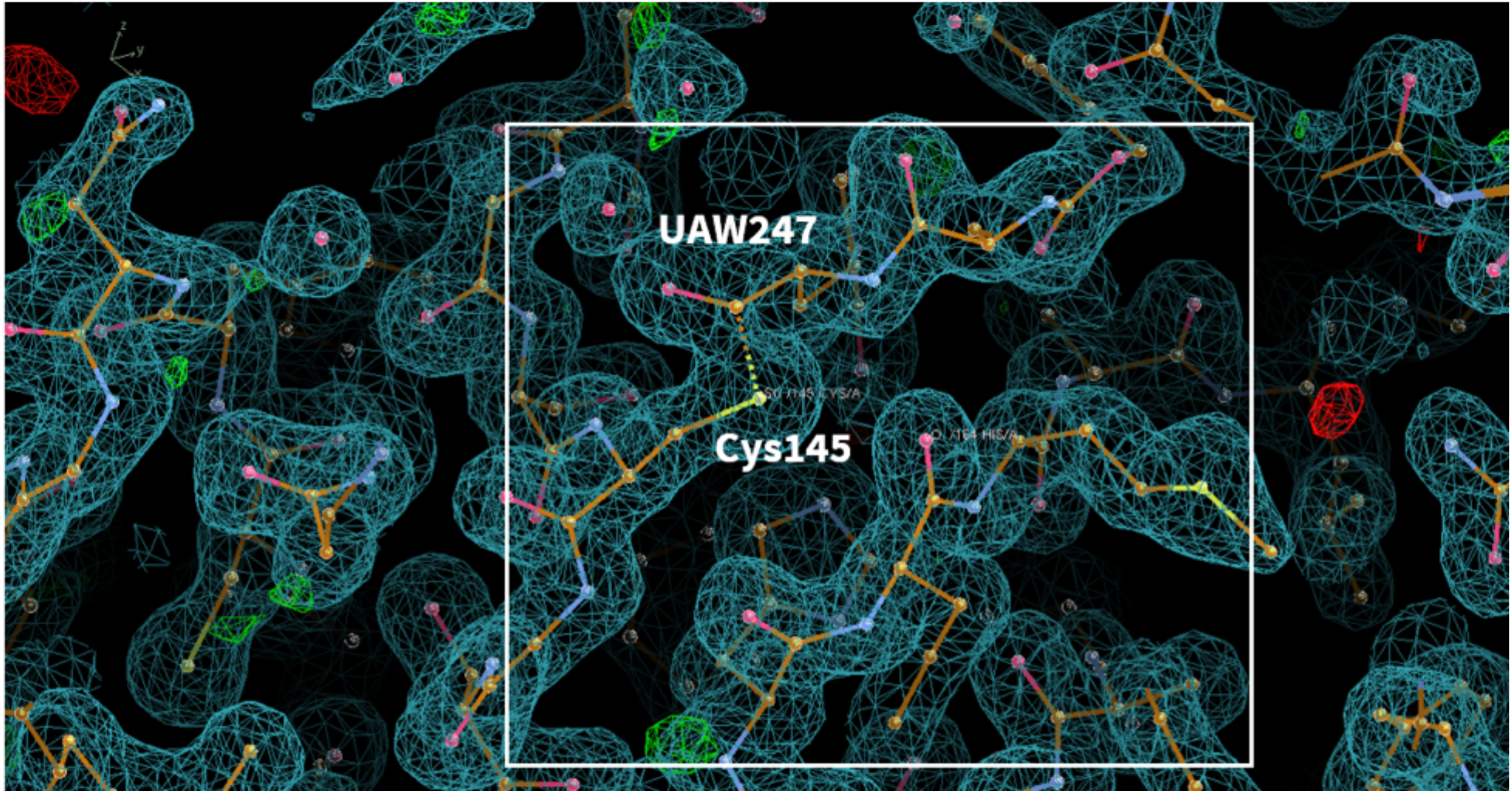
(covalent inhibitor) **PDB ID:** 6XBH **Ligand:** UAW247 **Resolution**: 1.60 Å **doi:** 10.2210/pdb6XBH/pdb **Author(s)**: Ma et al. **Deposited:** 2020 June 06 **Released:** 2020 June 17

**Fig. S4.9.**
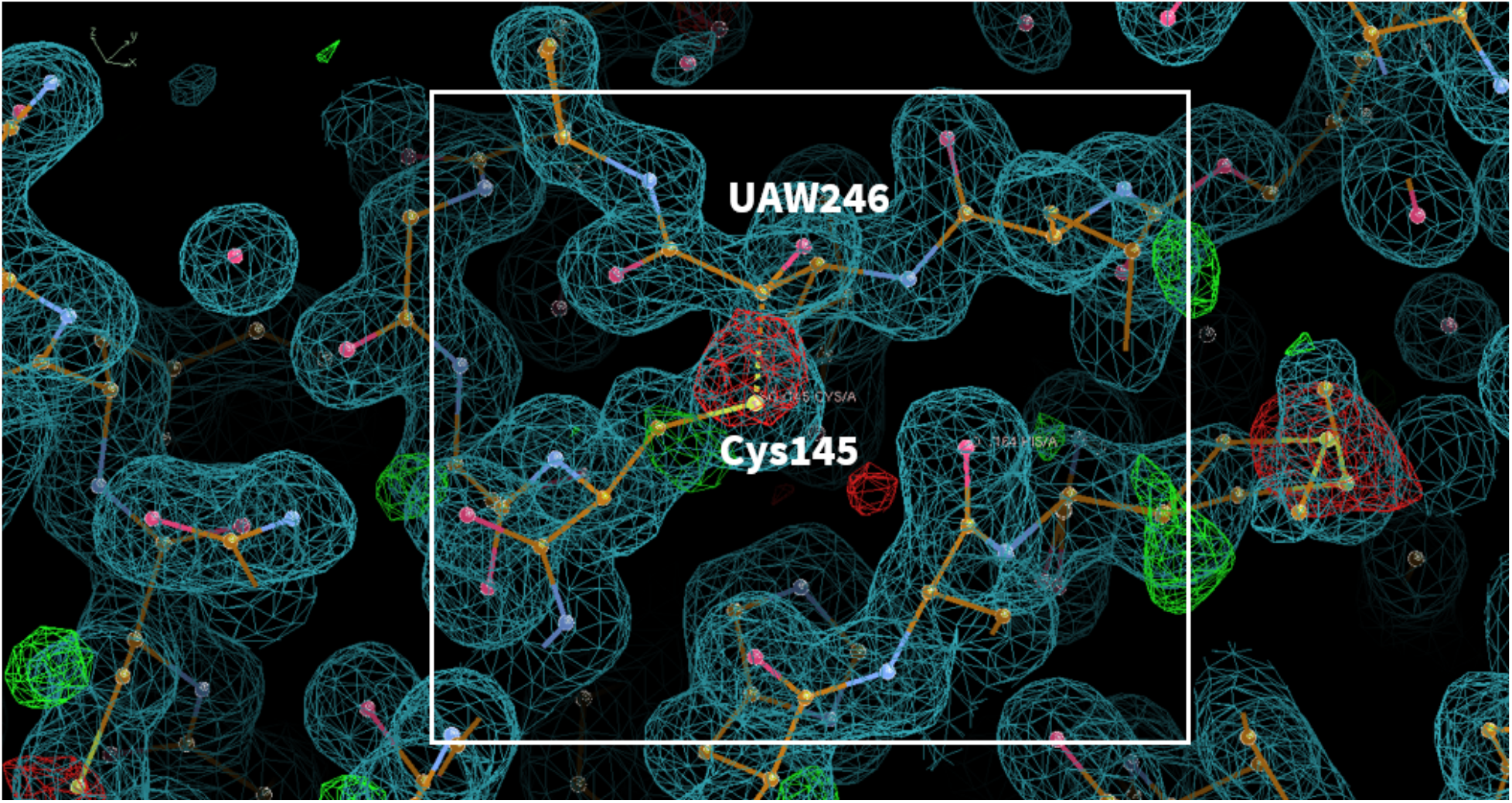
(covalent inhibitor) **PDB ID:** 6XBG **Ligand:** UAW246 **Resolution**: 1.60 Å **doi:** 10.2210/pdb6XBH/pdb **Author(s)**: Ma et al. **Deposited:** 2020 June 06 **Released:** 2020 June 17

**Fig. S4.10.**
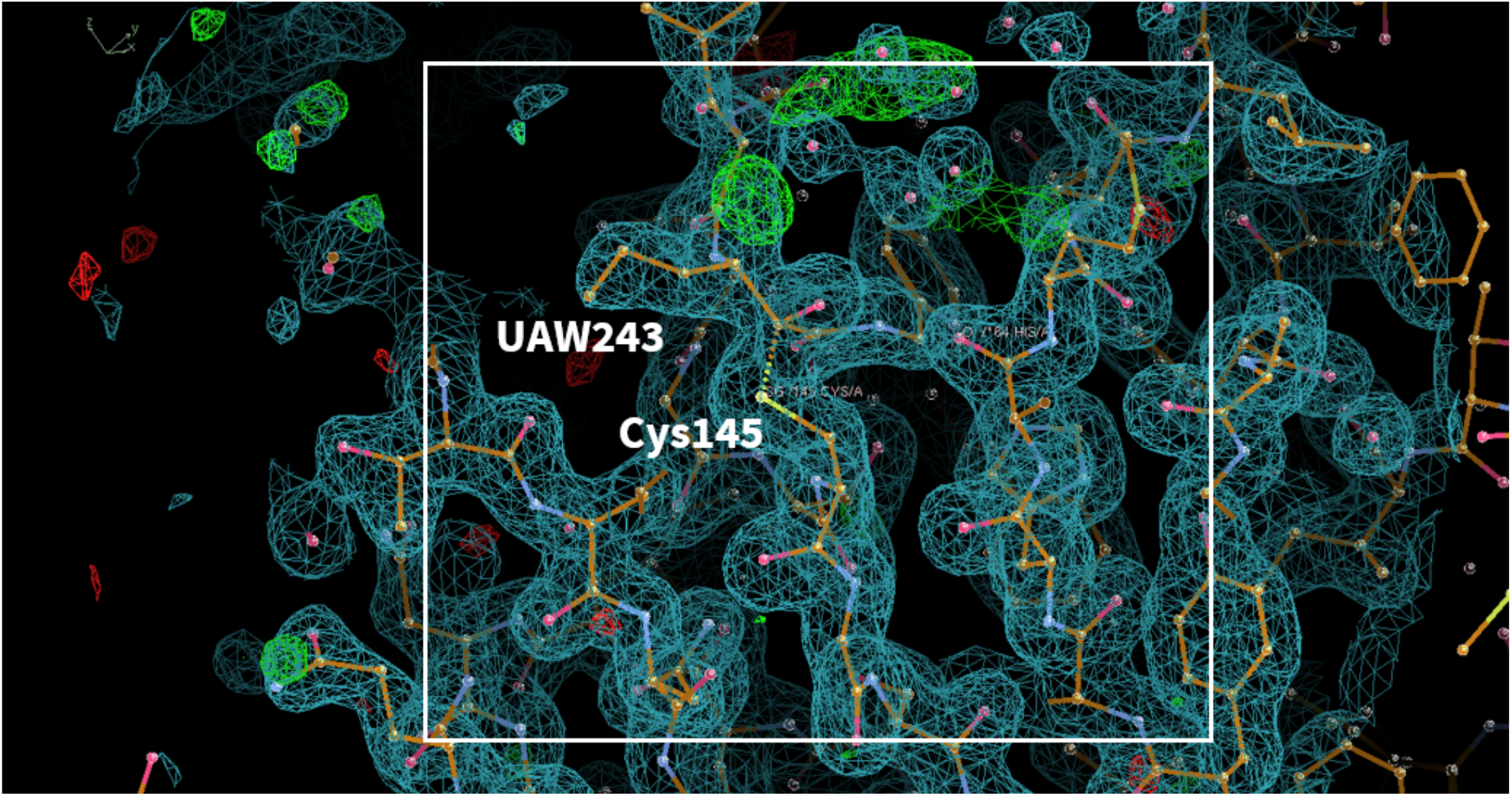
(covalent inhibitor) **PDB ID:** 6XFN **Ligand:** UAW243 **Resolution**: 1.70 Å **doi:** 10.2210/pdb6XFN/pdb **Author(s)**: Sacco et al. **Deposited:** 2020 June 15 **Released:** 2020 June 24

**Fig. S4.11.**
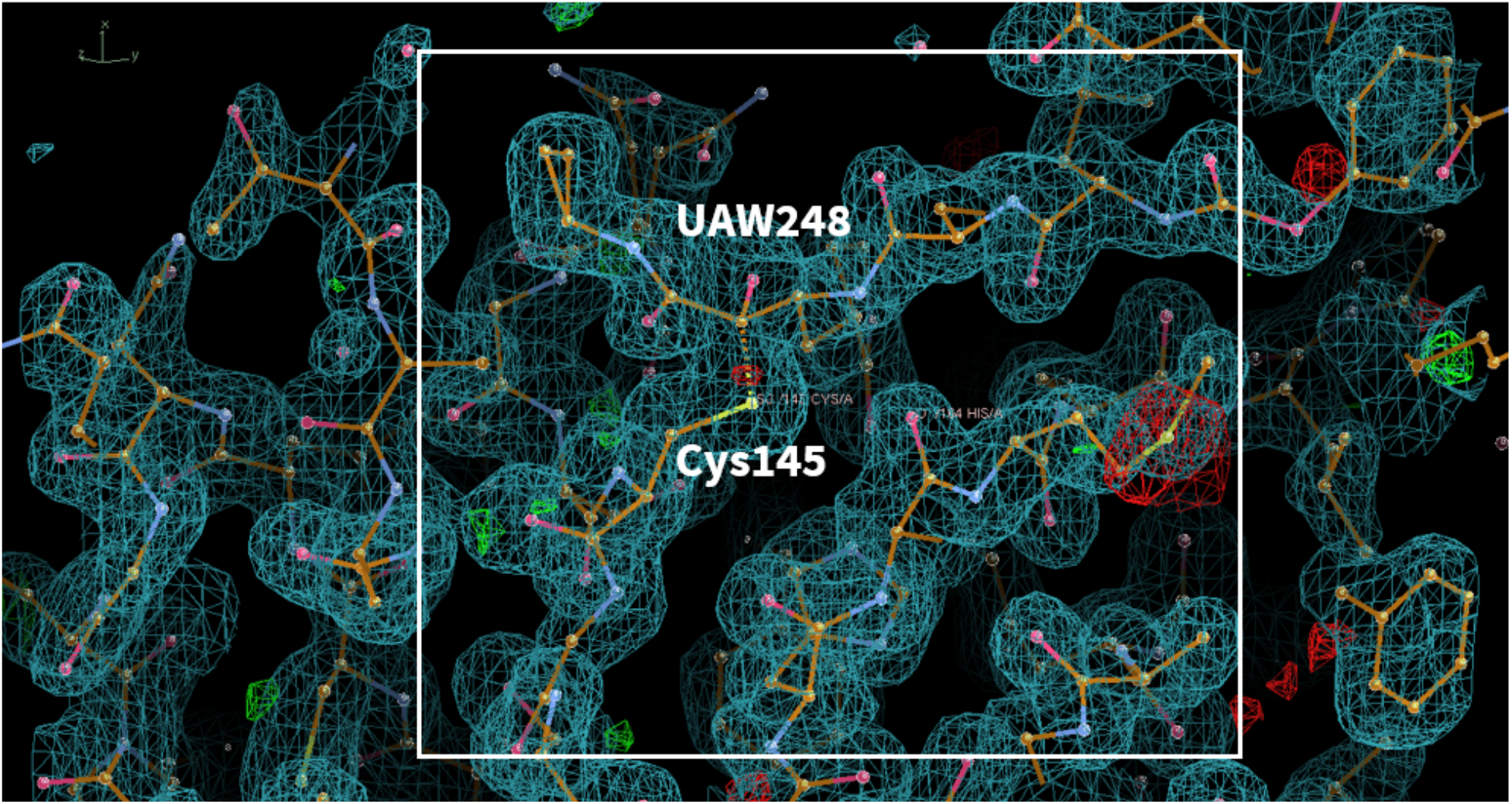
(covalent inhibitor) **PDB ID:** 6XBI **Ligand:** UAW248 **Resolution**: 1.70 Å **doi:** 10.2210/pdb6XBI/pdb **Author(s)**: Ma et al. **Deposited:** 2020 June 06 **Released:** 2020 June 17

**Fig. S4.12.**
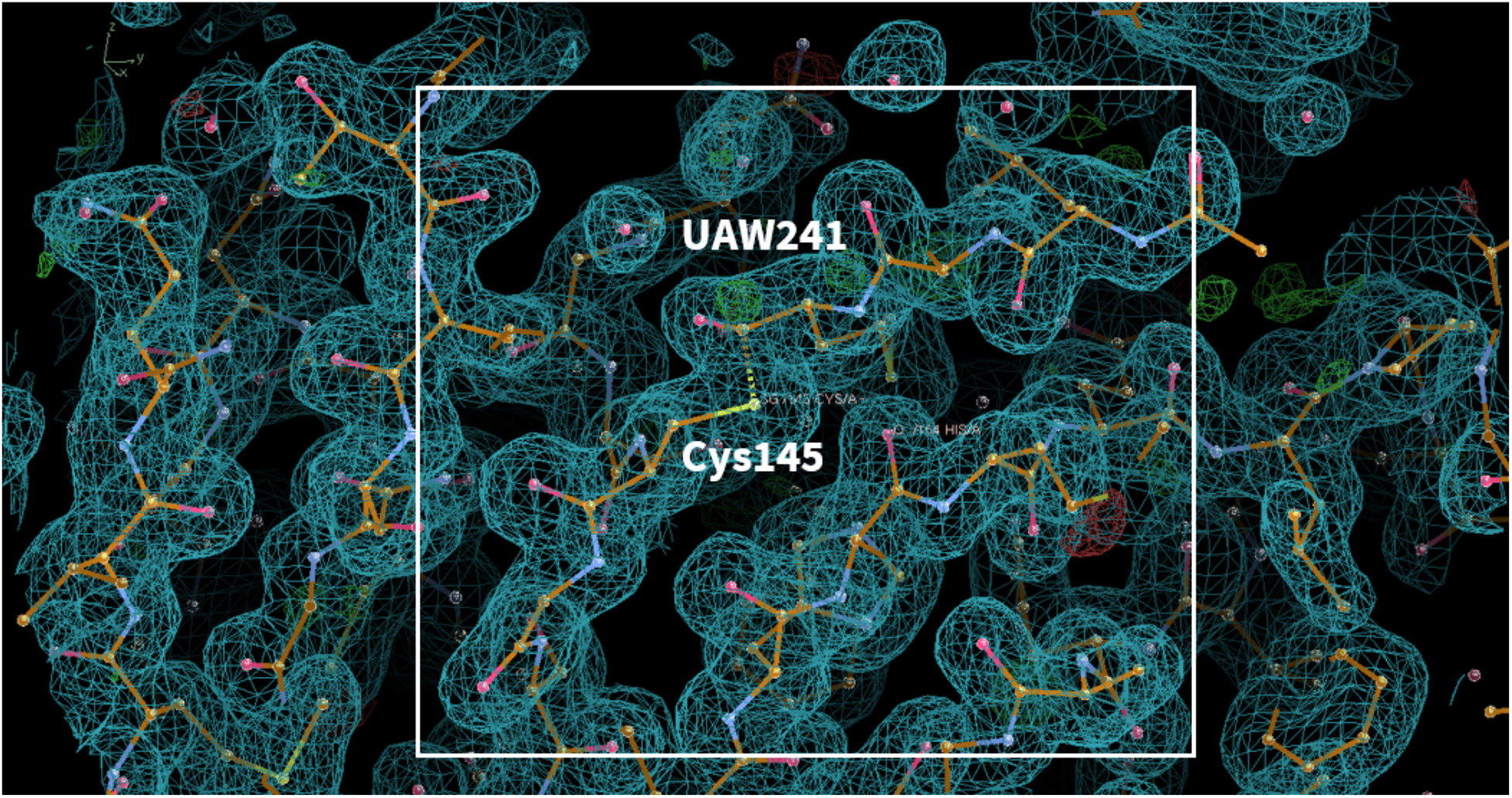
(covalent inhibitor) **PDB ID:** 6XA4 **Ligand:** UAW241 **Resolution**: 1.65 Å **doi:** 10.2210/pdb6XA4/pdb **Author(s)**: Sacco et al. **Deposited:** 2020 June 03 **Released:** 2020 June 17

**Fig. S4.13.**
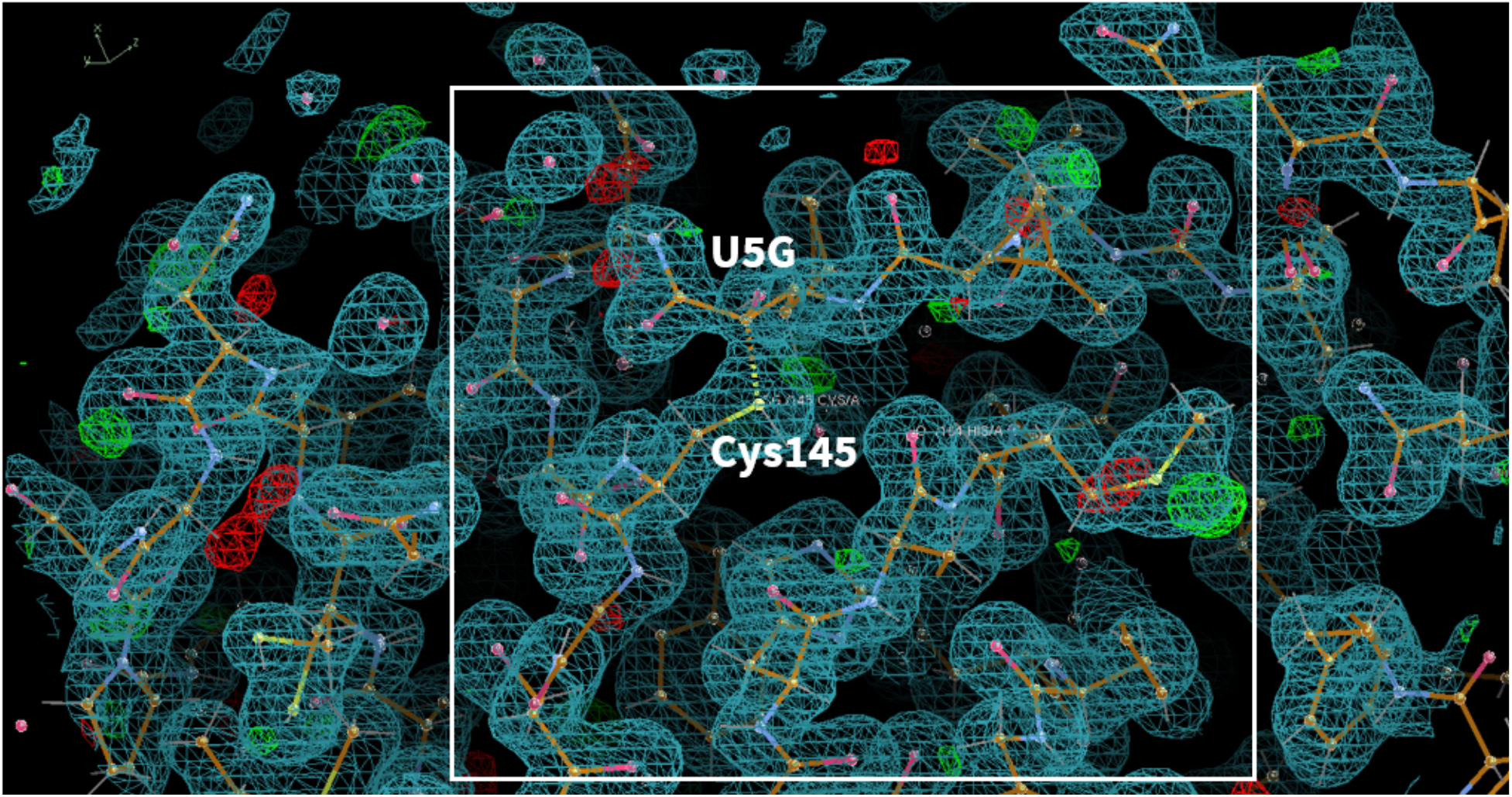
(covalent inhibitor) **PDB ID:** 6WNP **Ligand:** Boceprevir(U5G) **Resolution**: 1.44 Å **doi:** 10.2210/pdb6WNP/pdb **Author(s)**: Anson et al. **Deposited:** 2020 April 23 **Released:** 2020 May 06

**Fig. S4.14.**
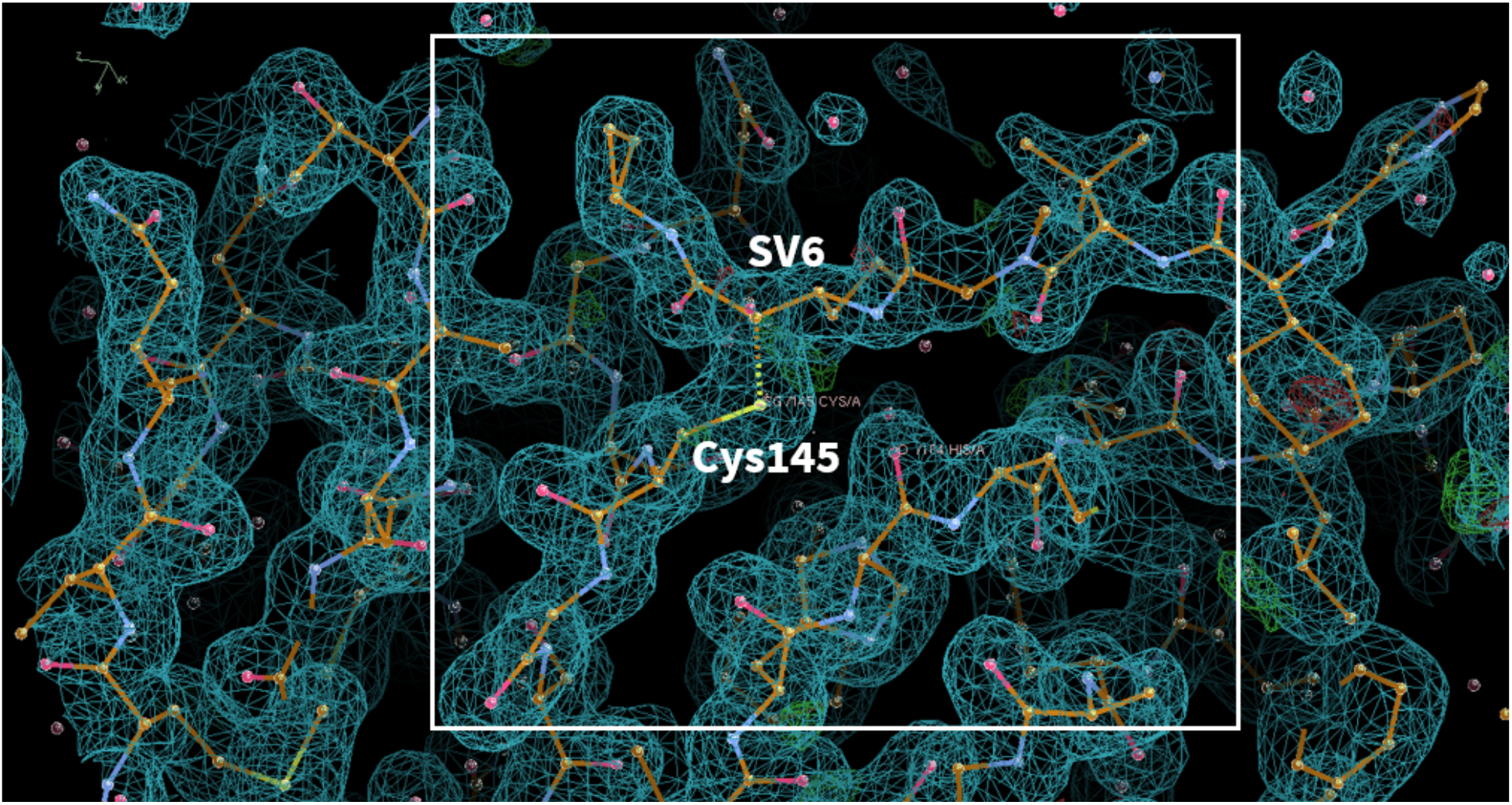
(covalent inhibitor) **PDB ID:** 7C7P **Ligand:** Telaprevir(SV6) **Resolution**: 1.74 Å **doi:** 10.2210/pdb7C7P/pdb **Author(s)**: Qiao et al. **Deposited:** 2020 May 26 **Released:** 2020 July 29

**Fig. S4.15.**
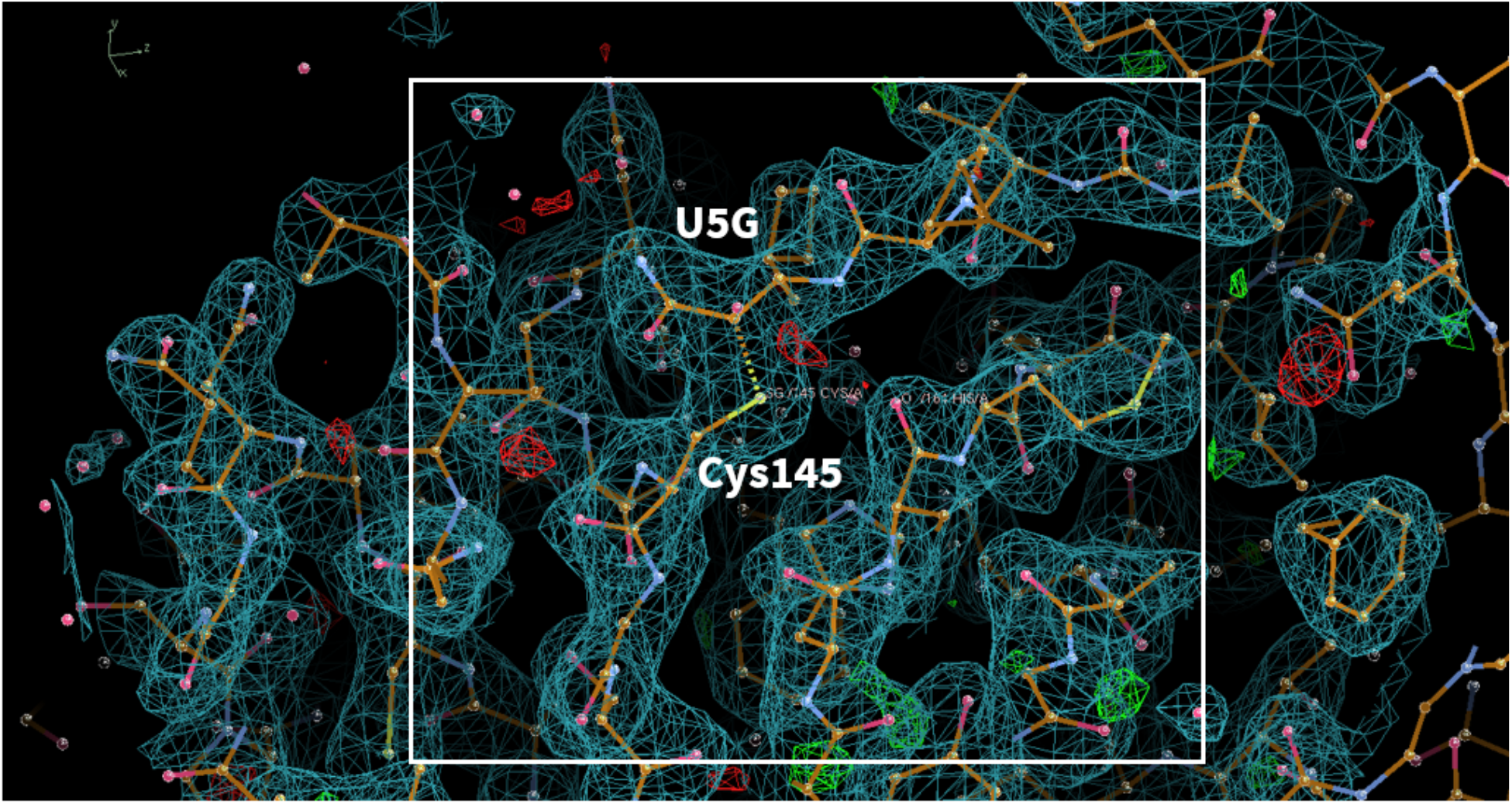
(covalent inhibitor) **PDB ID:** 7COM **Ligand:** Boceprevir(U5G) **Resolution**: 2.25 Å **doi:** 10.2210/pdb7COM/pdb **Author(s)**: Qiao et al. **Deposited:** 2020 August 04 **Released:** 2020 August 19

**Fig. S4.16.**
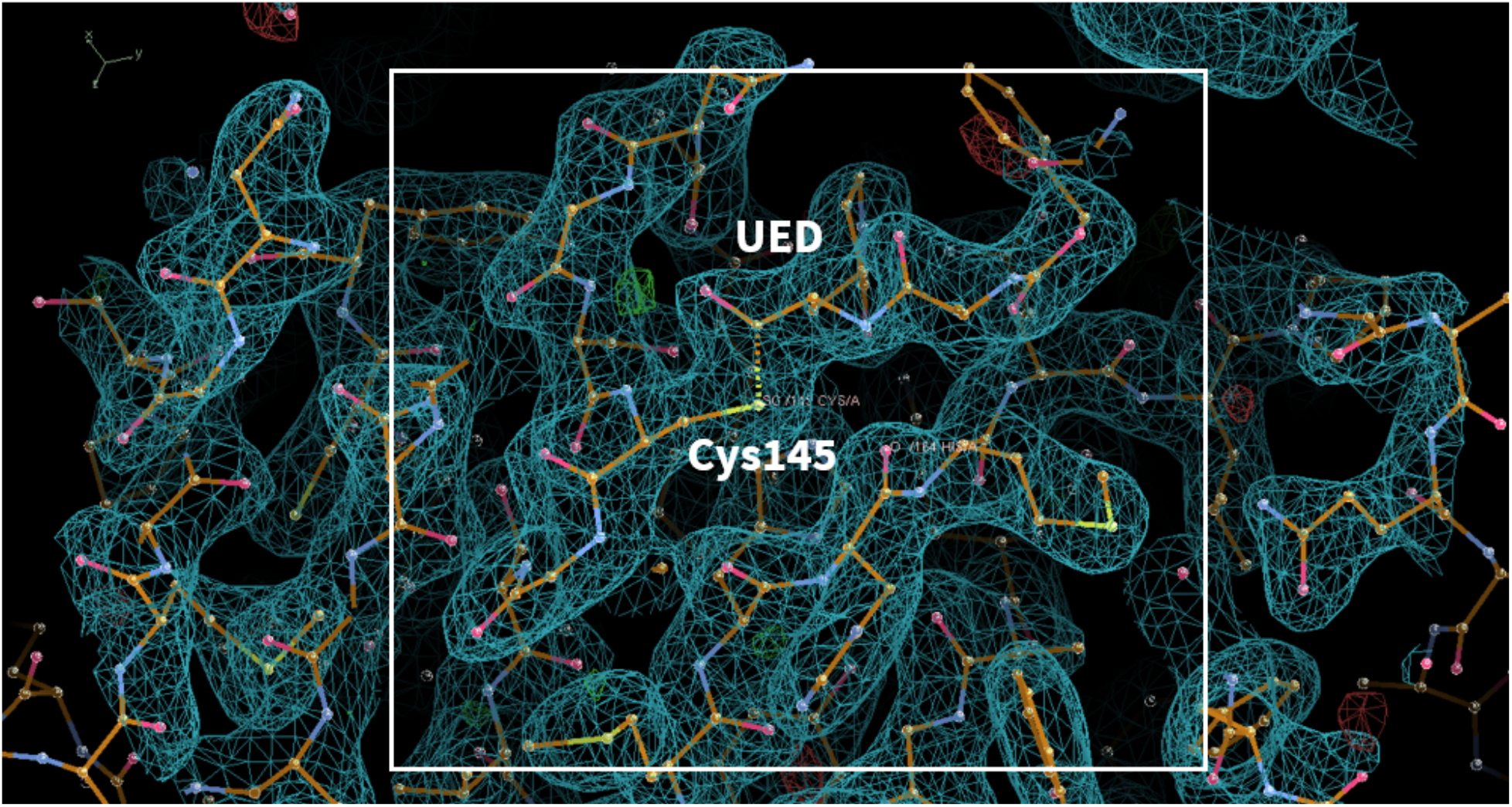
(covalent inhibitor) **PDB ID:** 6WTK **Ligand:** Feline(UED) **Resolution**: 2.00 Å **doi:** 10.2210/pdb6WTK/pdb **Author(s)**: Khan et al. **Deposited:** 2020-May-02 **Released:** 2020-May-20

**Fig. S4.17.**
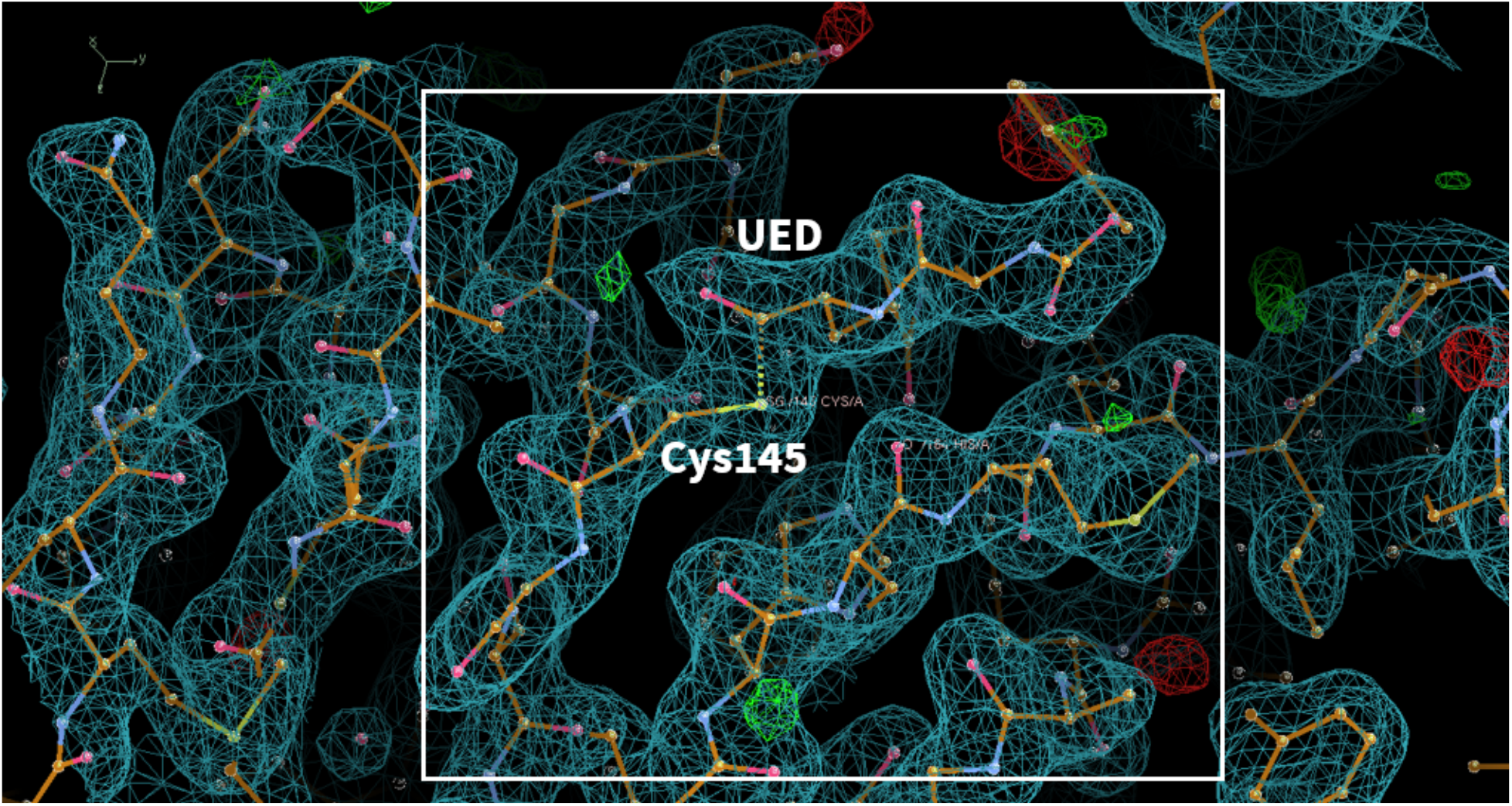
(covalent inhibitor) **PDB ID:** 6WTJ **Ligand:** Feline(UED) **Resolution**: 1.90 Å **doi:** 10.2210/pdb6WTJ/pdb **Author(s)**: Khan et al. **Deposited:** 2020 May 02 **Released:** 2020 May 20

**Fig. S4.18.**
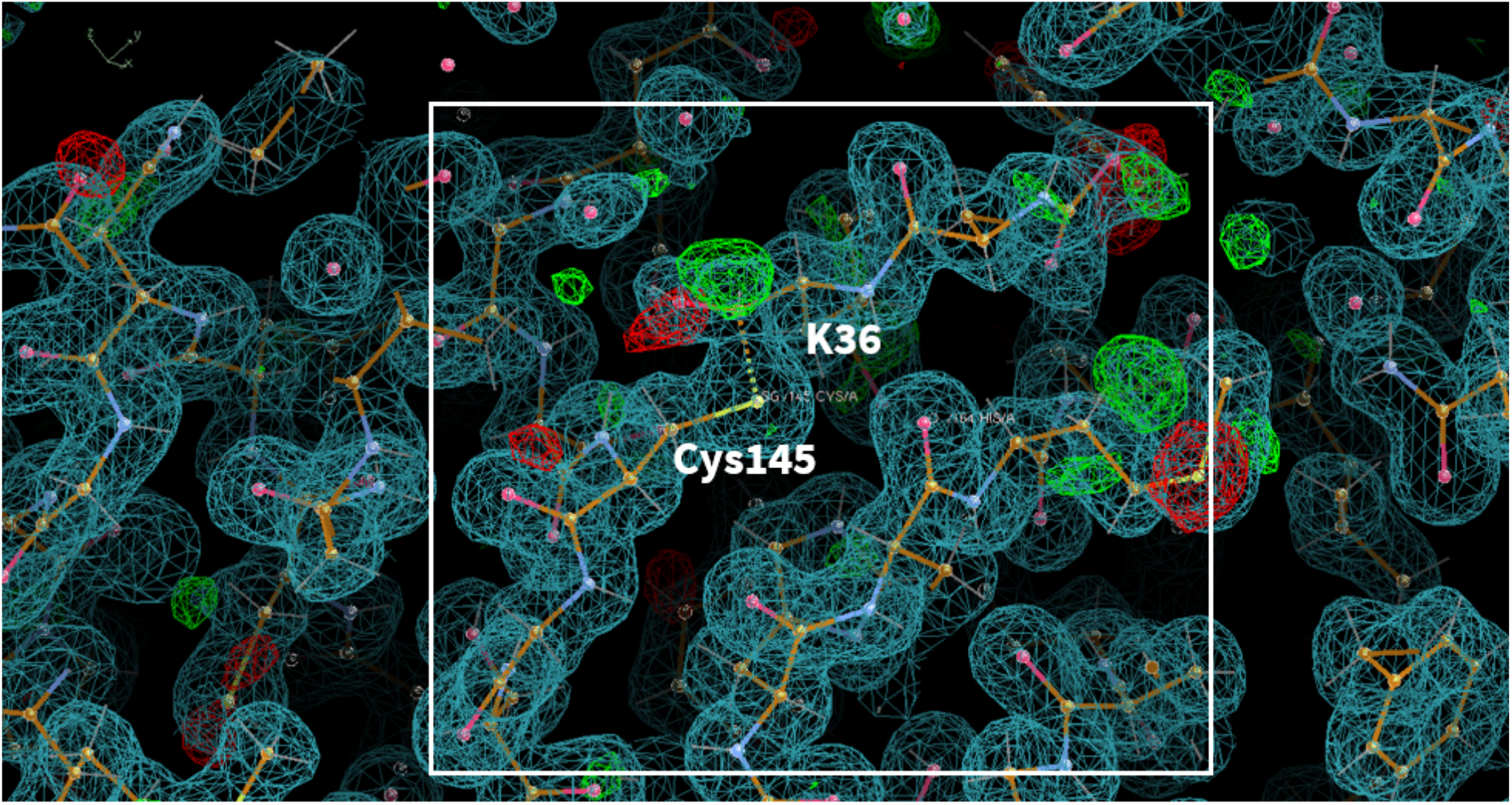
(covalent inhibitor) **PDB ID:** 7BRR **Ligand:** GC-376(K36) **Resolution**: 1.40 Å **doi:** 10.2210/pdb7BRR/pdb **Author(s)**: Fu et al. **Deposited:** 2020 March 29 **Released:** 2020 May 13

**Fig. S4.19.**
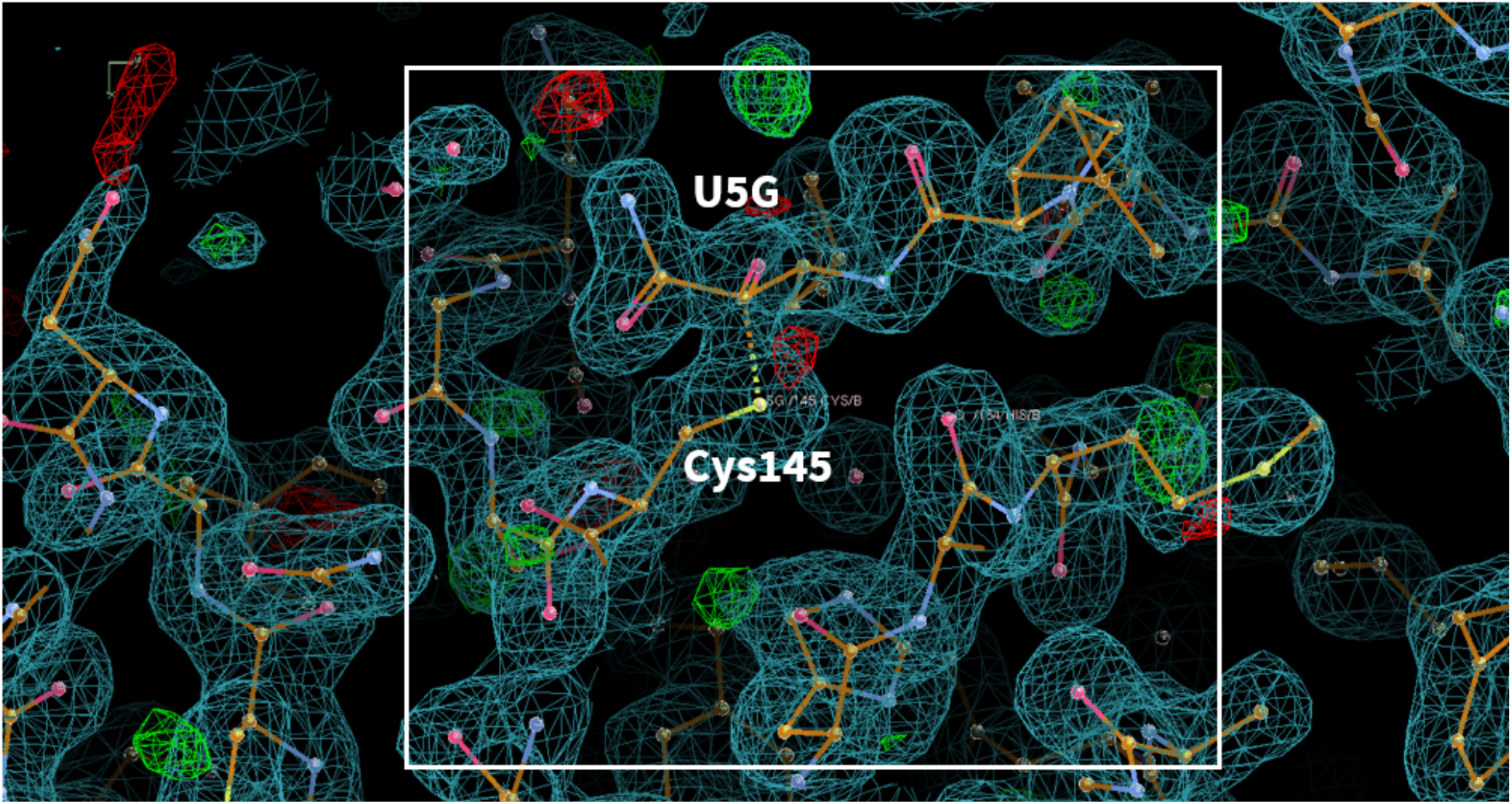
(covalent inhibitor) **PDB ID:** 7BRP **Ligand:** Boceprevir(U5G) **Resolution**: 1.95 Å **doi:** 10.2210/pdb7BPR/pdb **Author(s)**: Masutani et al. **Deposited:** 2020 March 23 **Released:** 2020 July 15

**Fig. S4.20.**
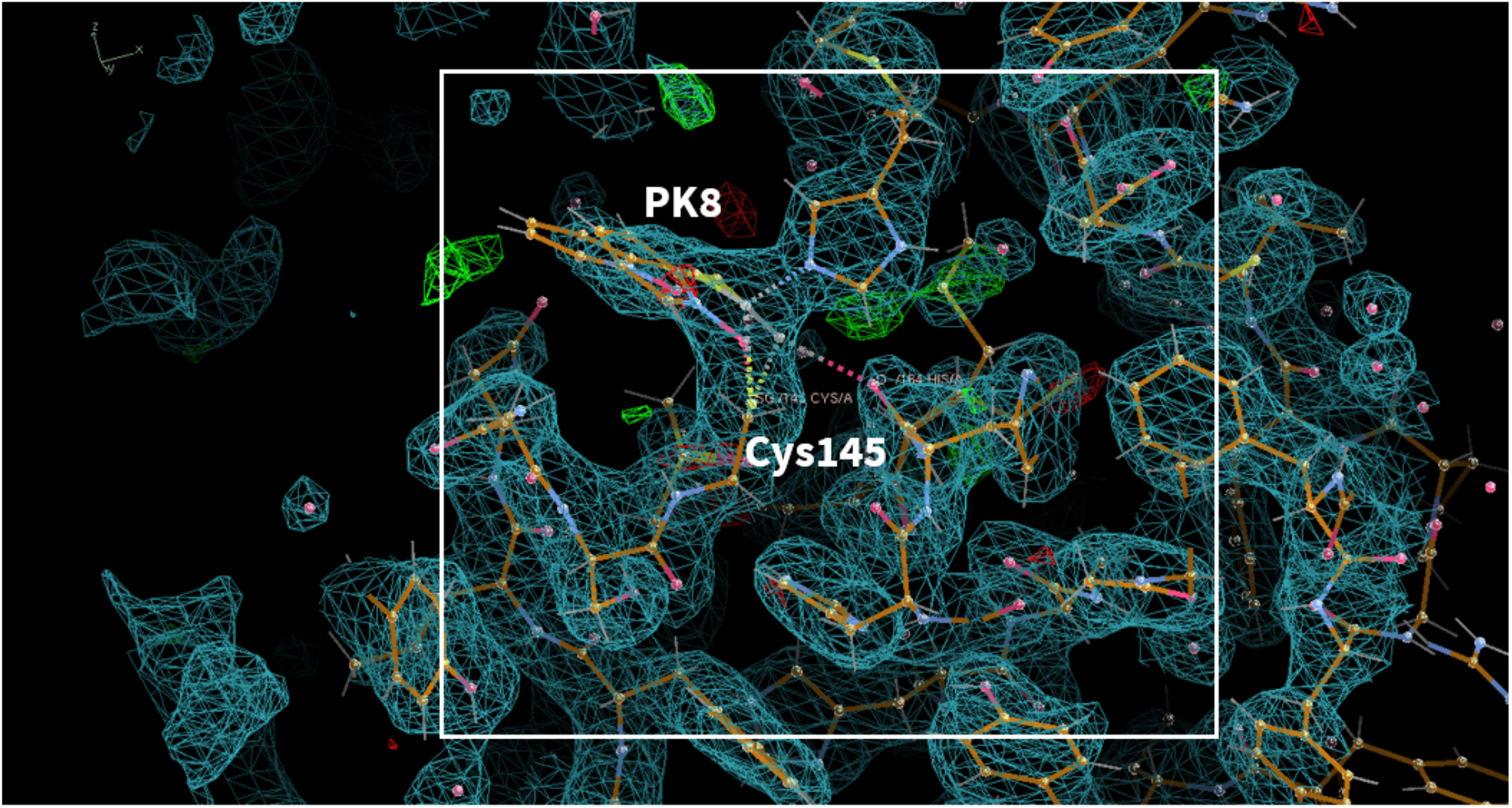
(covalent inhibitor) **PDB ID:** 6YT8 **Ligand:** pyrithione zinc(PK8) **Resolution**: 2.50 Å **doi:** 10.2210/pdb6YT8/pdb **Author(s)**: Guenther et al. **Deposited:** 2020 April 24 **Released:** 2020 May 06

**Fig. S4.21.**
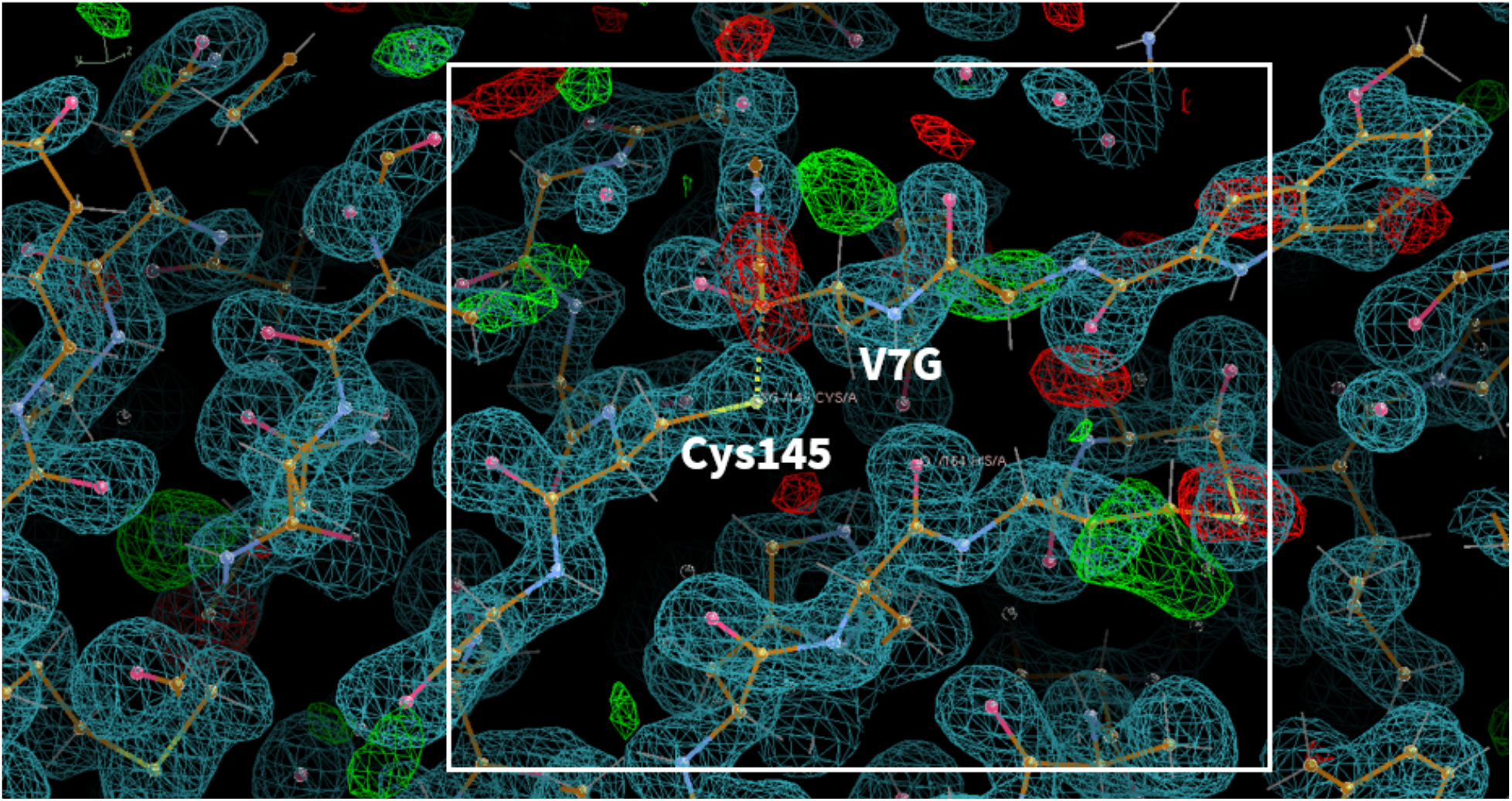
(covalent inhibitor) **PDB ID:** 6XR3 **Ligand:** GRL-024-20(V7G) **Resolution**: 1.45 Å **doi:** 10.2210/pdb6XR3/pdb **Author(s)**: Anson et al. **Deposited:** 2020 July 10 **Released:** 2020 August 19

**Fig. S4.22.**
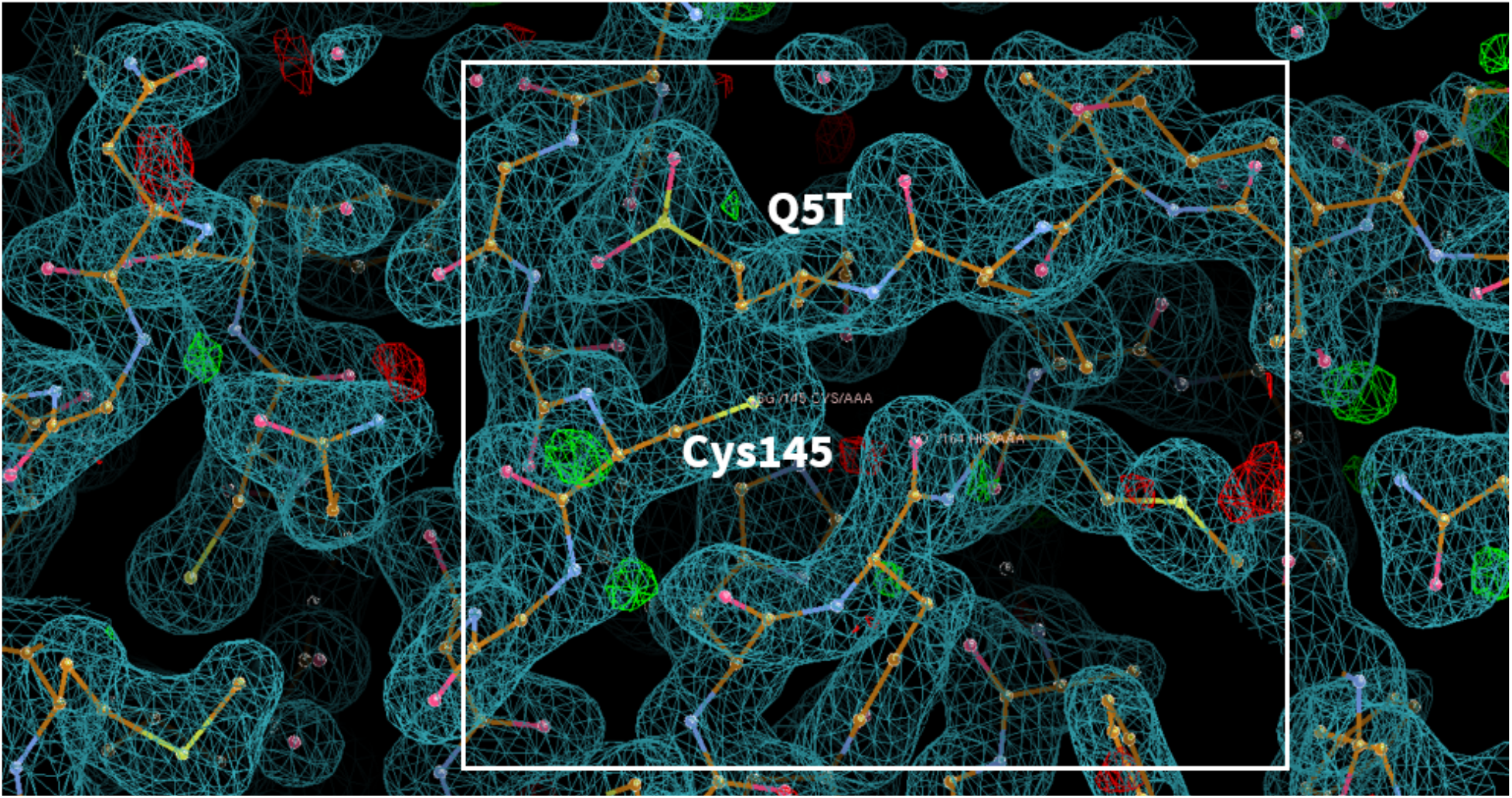
(covalent inhibitor) **PDB ID:** 6Z2E **Ligand:** biotin-PEG(4)-Abu-Tle-Leu-Gln-vinylsulfone(Q5T) **Resolution**: 1.70 Å **doi:** 10.2210/pdb6Z2E/pdb **Author(s)**: Zhang et al. **Deposited:** 2020 May 15 **Released:** 2020 June 7

**Fig. S4.23.**
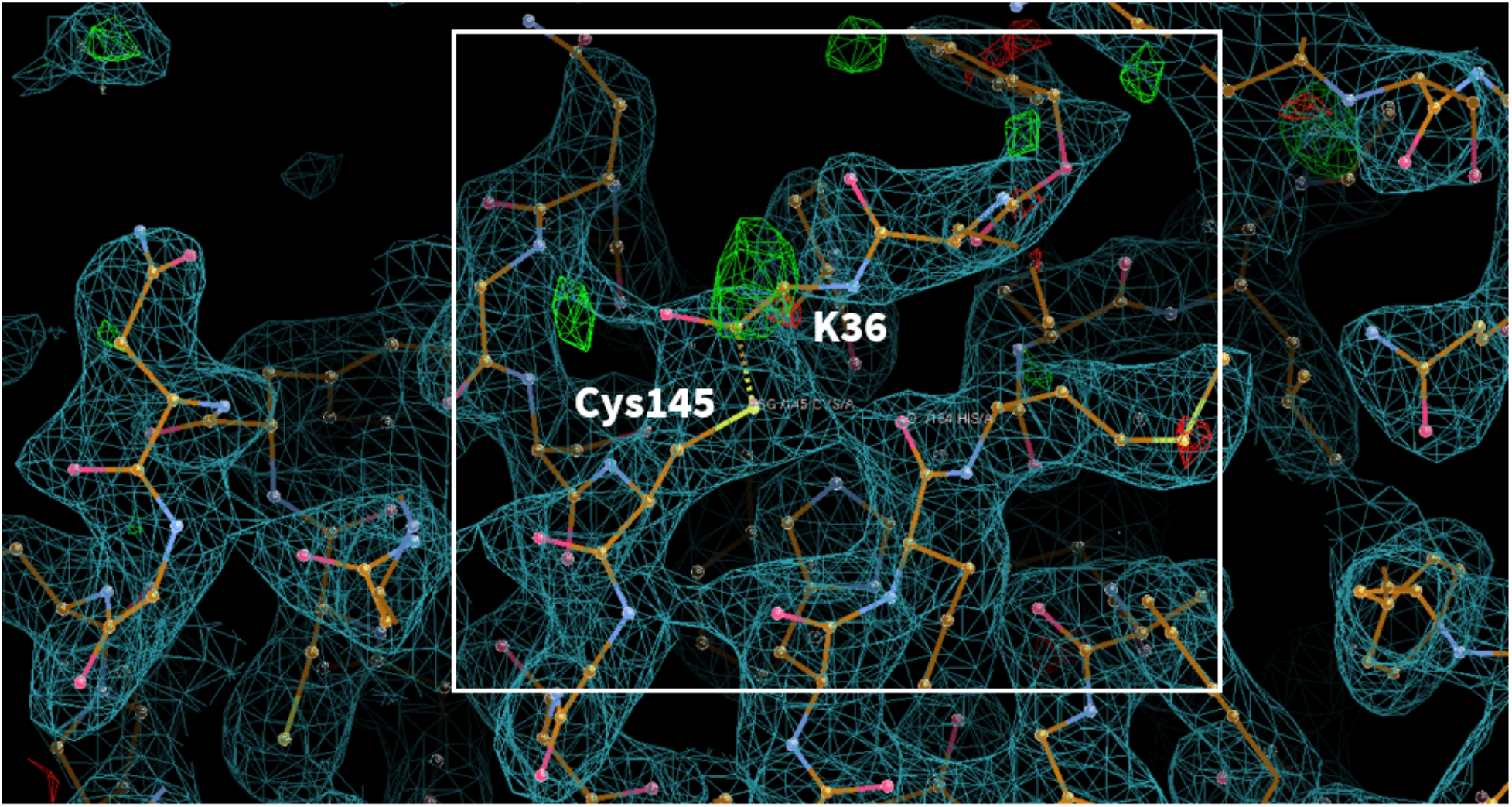
(covalent inhibitor) **PDB ID:** 7C8U **Ligand:** GC-376(K36) **Resolution**: 2.35 Å **doi:** 10.2210/pdb7C8U/pdb **Author(s)**: Zhang et al. **Deposited:** 2020 June 03 **Released:** 2020 June 24

**Fig. S4.24.**
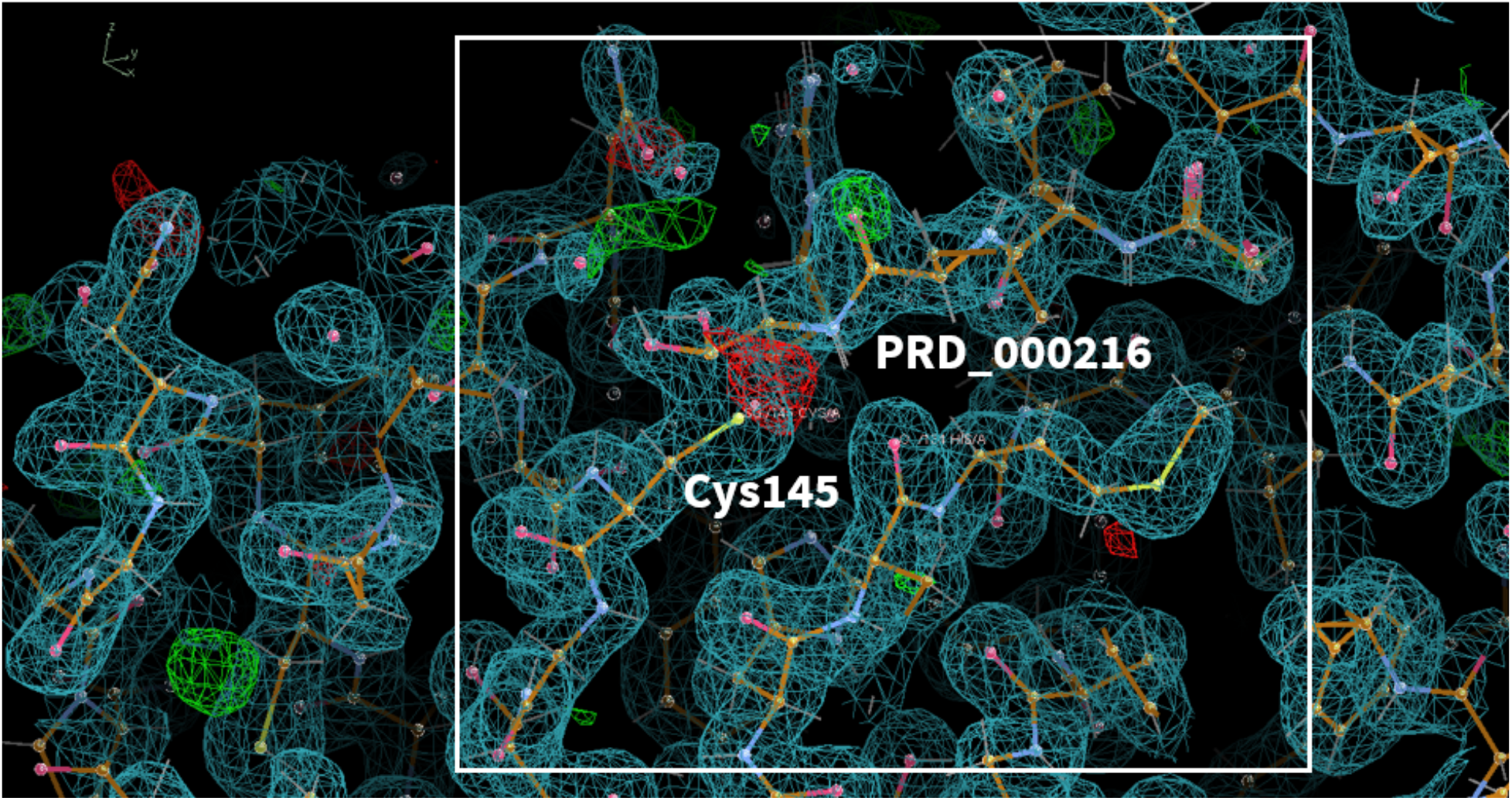
(covalent inhibitor) **PDB ID:** 6YZ6 **Ligand:** Leupeptin(PRD_000216) **Resolution:** 1.70 Å **doi:** 10.2210/pdb6YZ6/pdb **Author(s)**: Guenther et al. **Deposited:** 2020 May 06 **Released:** 2020 May 20

**Fig. S4.25.**
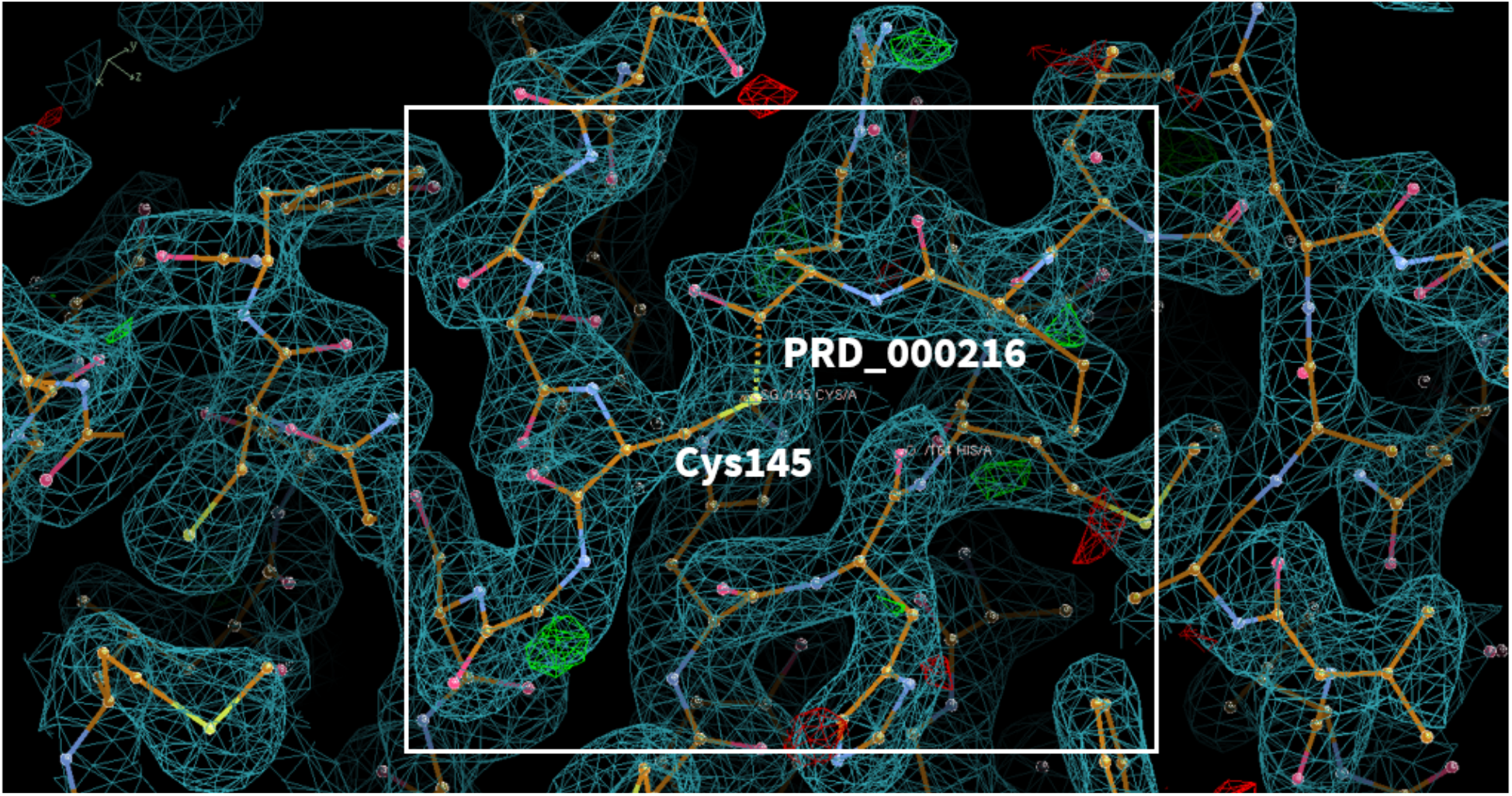
(covalent inhibitor) **PDB ID:** 6XCH **Ligand:** Leupeptin(PRD_000216) **Resolution:** 2.20 Å **doi:** 10.2210/pdb6XCH/pdb **Author(s)**: Kneller et al. **Deposited:** 2020 June 08 **Released:** 2020 June 17

**Fig. S4.26.**
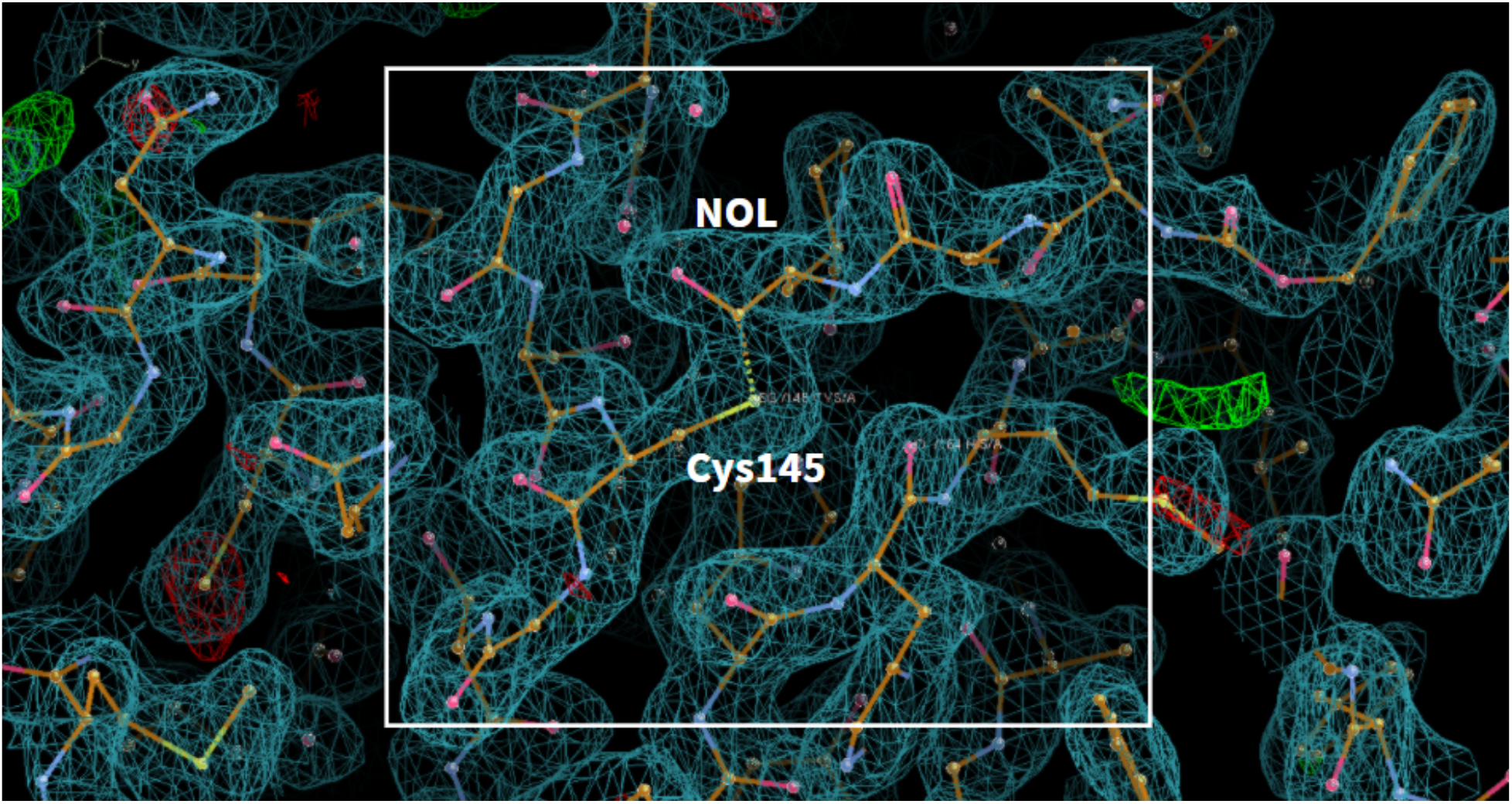
(covalent inhibitor) **PDB ID:** 7C8T **Ligand:** TG-0205221(NOL) **Resolution:** 2.05 Å **doi:** 10.2210/pdb7C8T/pdb **Author(s)**: Kuo et al. **Deposited:** 2020 June 03 **Released:** 2020 June 17

**Fig. S4.27.**
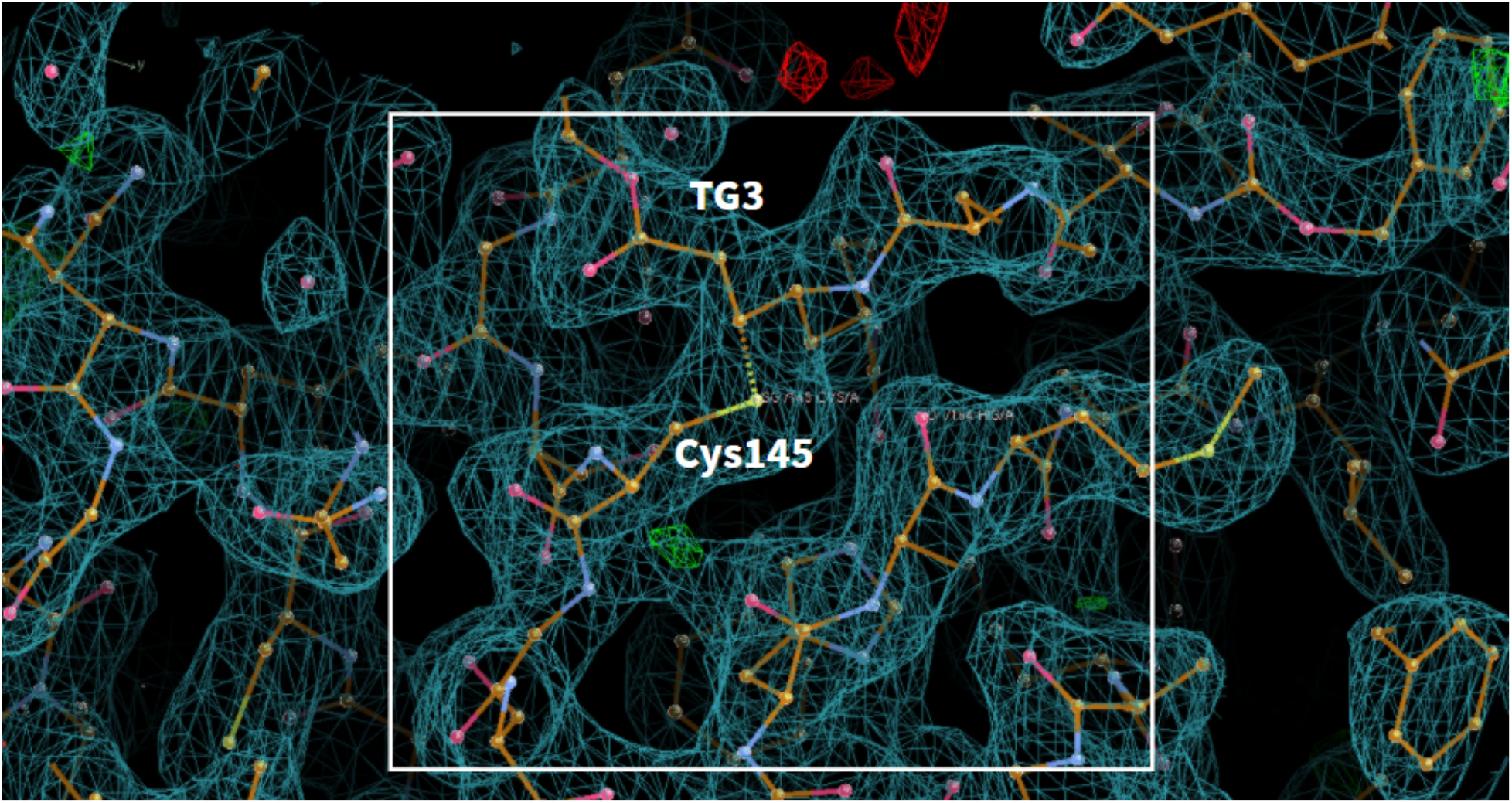
(covalent inhibitor) **PDB ID:** 7C8R **Ligand:** TG-0203770(TG3) **Resolution:** 2.30 Å **doi:** 10.2210/pdb7C8R/pdb **Author(s)**: Kuo et al. **Deposited:** 2020 June 03 **Released:** 2020 June 17

**Fig. S4.28.**
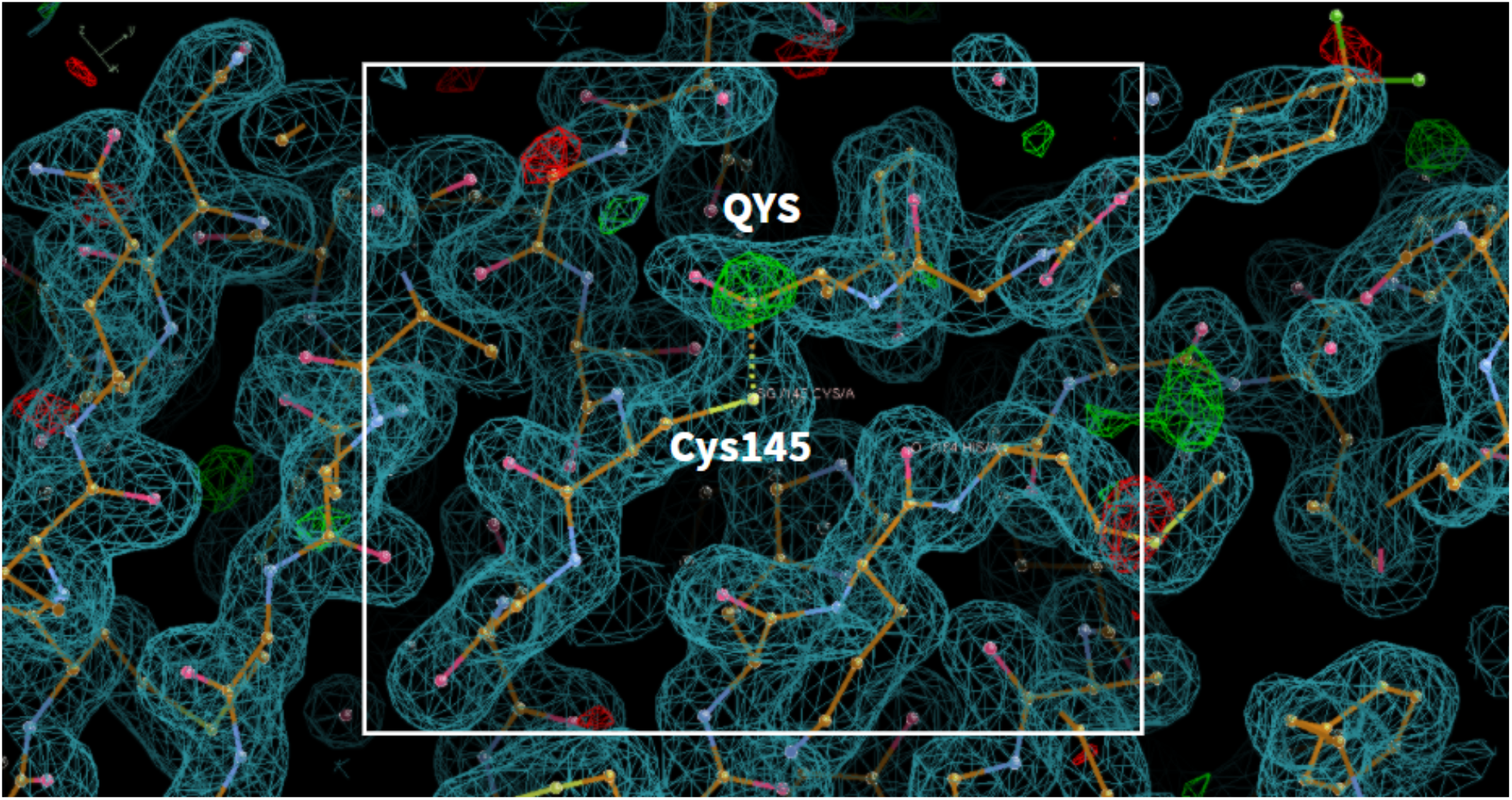
(covalent inhibitor) **PDB ID:** 6XMK **Ligand:** 7j(QYS) **Resolution:** 1.70 Å **doi:** 10.2210/pdb6XMK/pdb **Author(s)**: Rathnayake et al. **Deposited:** 2020 June 30 **Released:** 2020 July 08

**Fig. S4.29.**
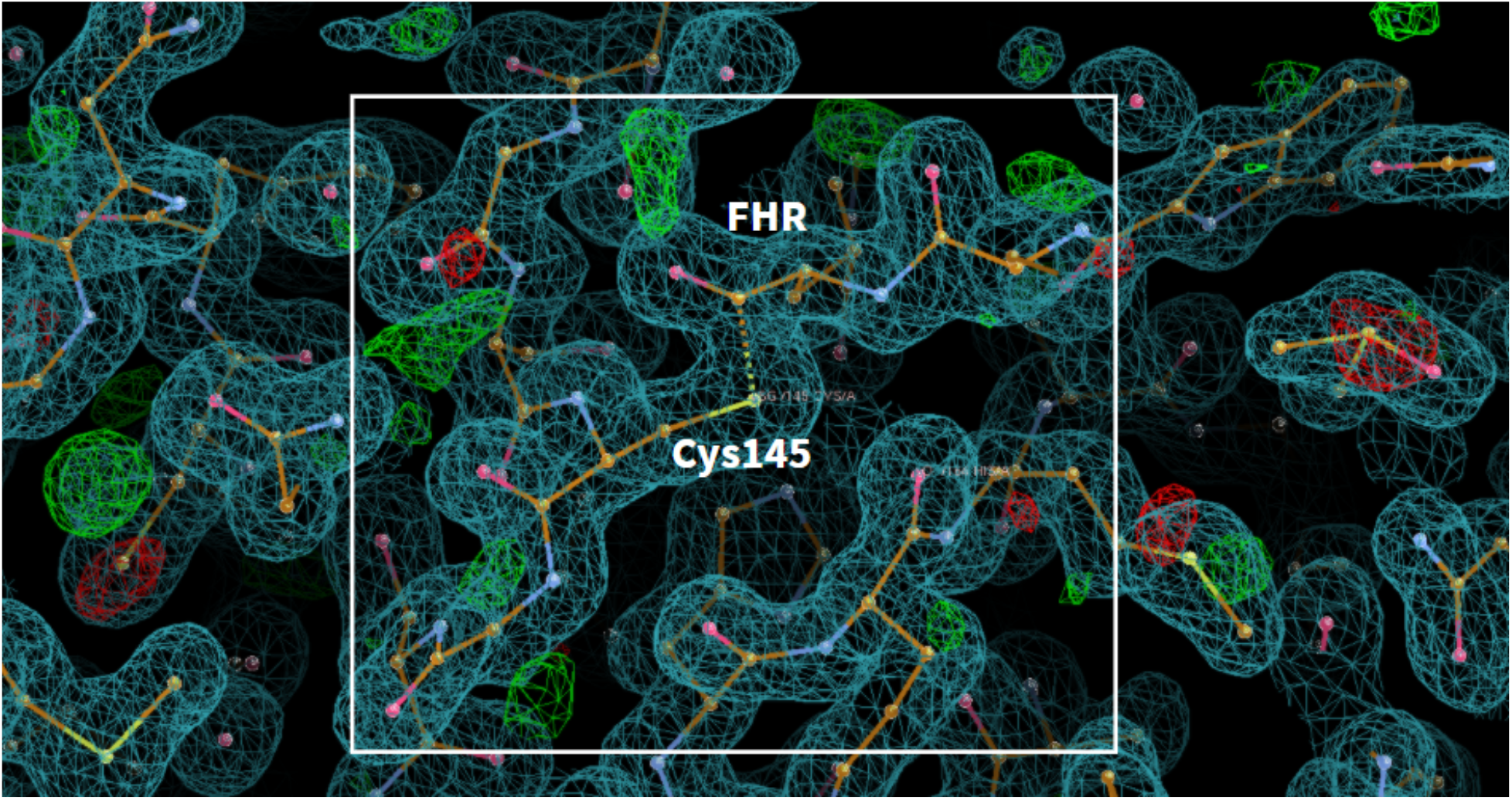
(covalent inhibitor) **PDB ID:** 6LZE **Ligand:** 11a(FHR) **Resolution:** 1.50 Å **doi:** 10.2210/pdb6LZE/pdb **Author(s)**: Dai et al. **Deposited:** 2020 February 19 **Released:** 2020 April 29

**Fig. S4.30.**
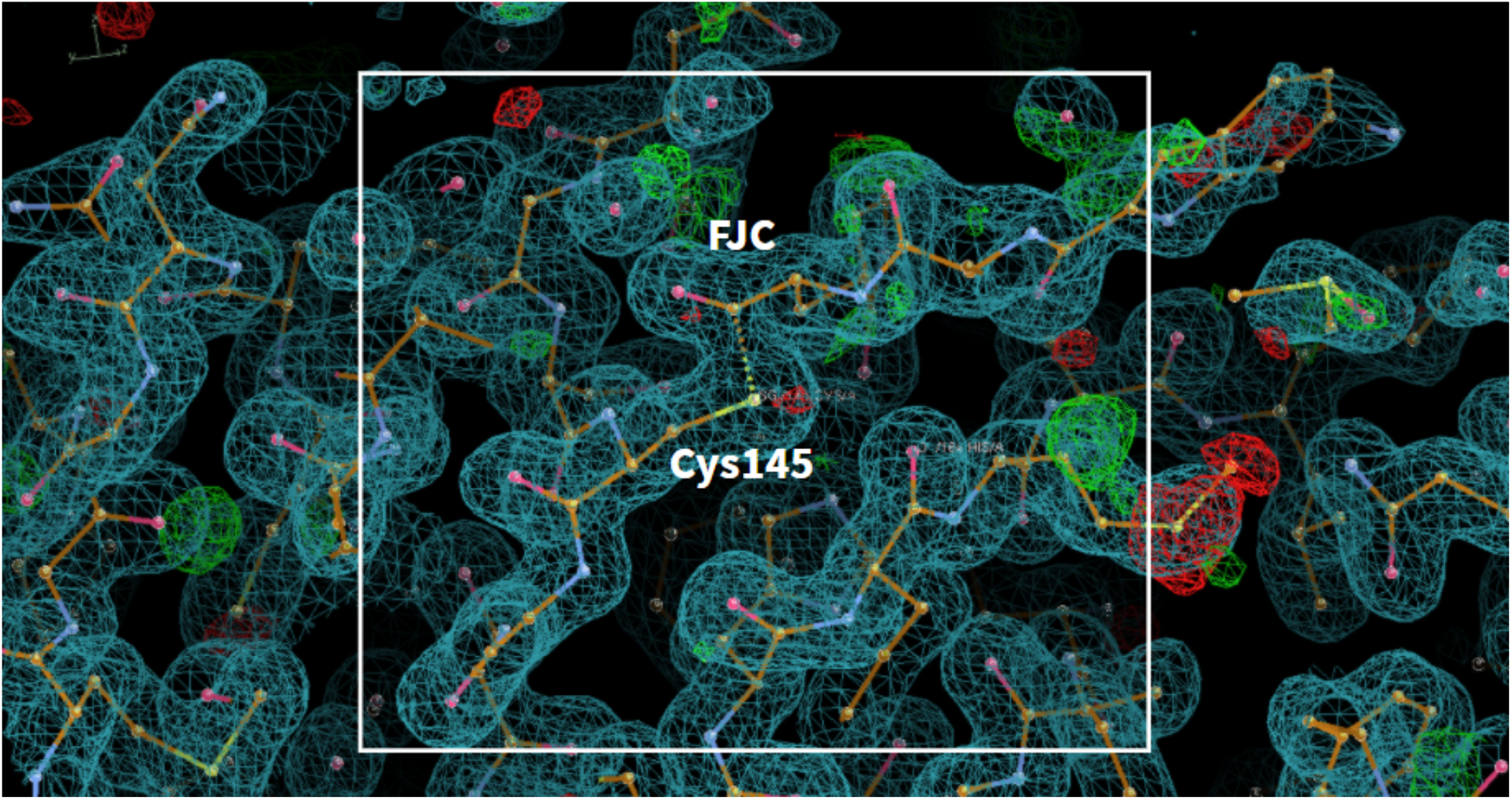
(covalent inhibitor) **PDB ID:** 6M0K **Ligand:** 11b(FJC) **Resolution:** 1.50 Å **doi:** 10.2210/pdb6M0K/pdb **Author(s)**: Dai et al. **Deposited:** 2020 February 22 **Released:** 2020 April 29

**Fig. S4.31.**
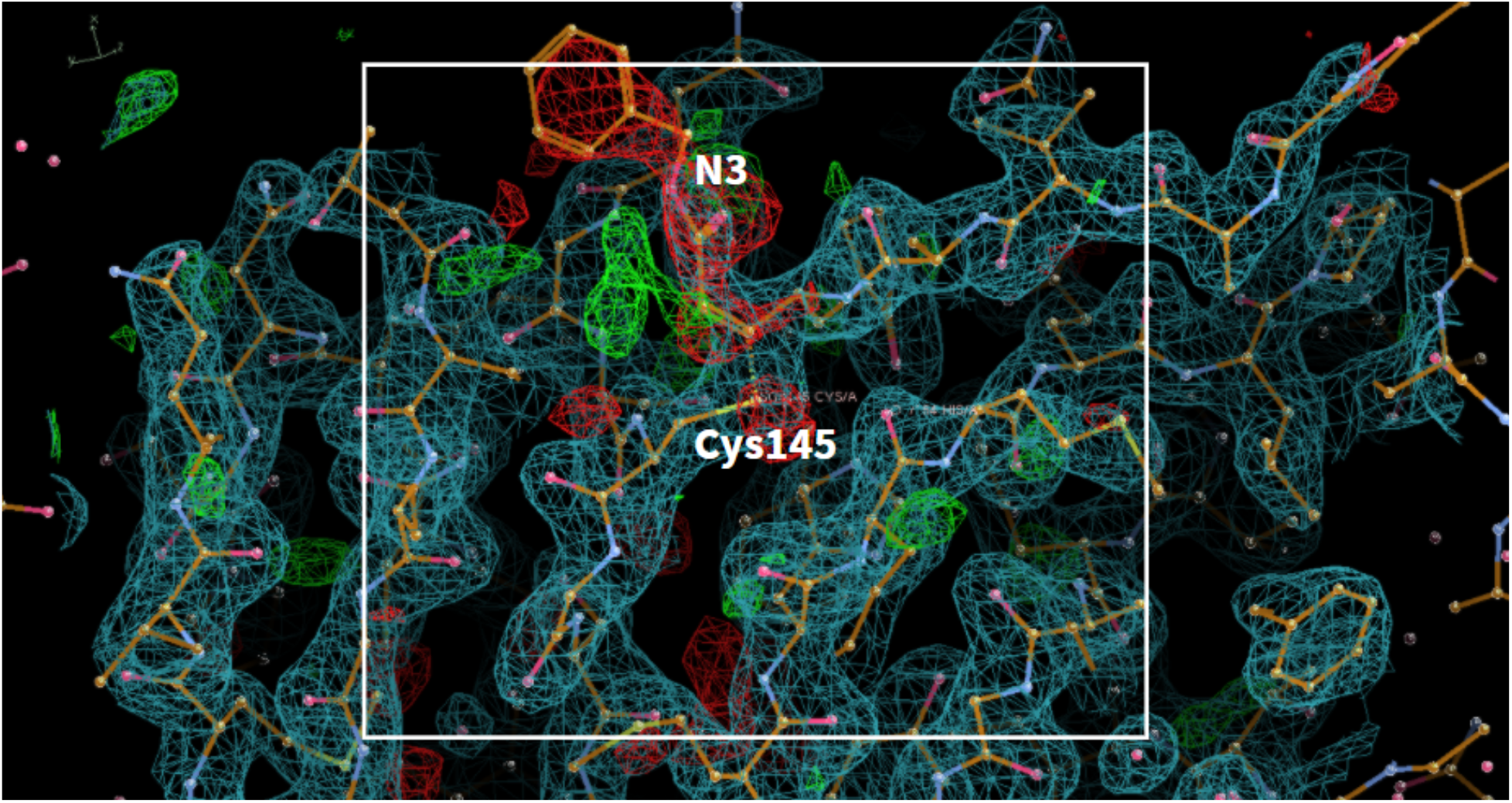
(covalent inhibitor) **PDB ID:** 6LU7 **Ligand:** N3 **Resolution:** 2.16 Å **doi:** 10.2210/pdb6LU7/pdb **Author(s)**: Jin et al. **Deposited:** 2020 January 26 **Released:** 2020 February 05

**Fig. S4.32.**
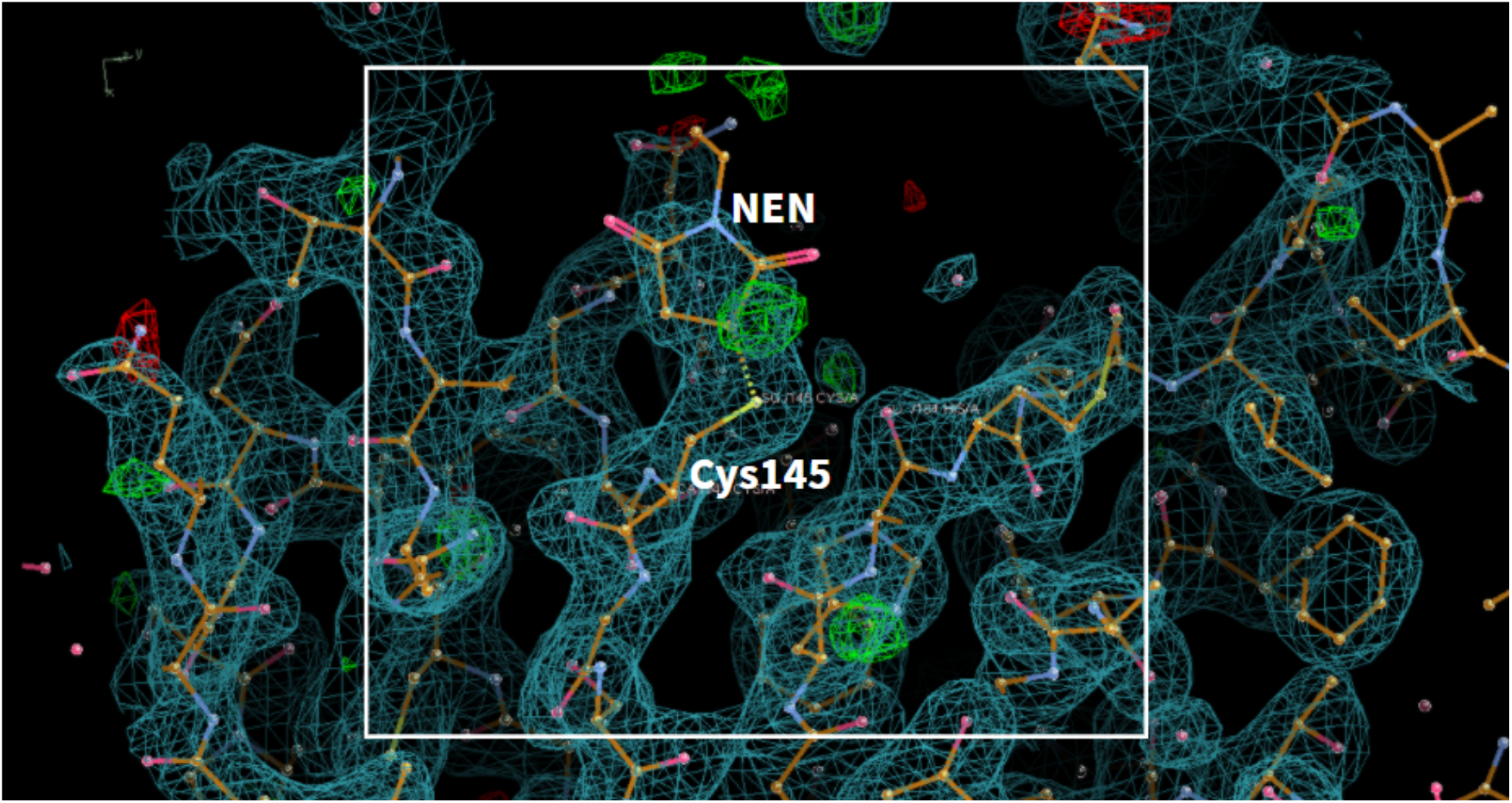
(covalent inhibitor) **PDB ID:** 6XB2 **Ligand:** 1-ETHYL-PYRROLIDINE-2,5-DIONE(NEN) **Resolution:** 2.12 A **doi:** 10.2210/pdb6XB2/pdb **Author(s)**: Kneller et al. **Deposited:** 2020 June 05 **Released:** 2020 June 17

**Fig. S4.33.**
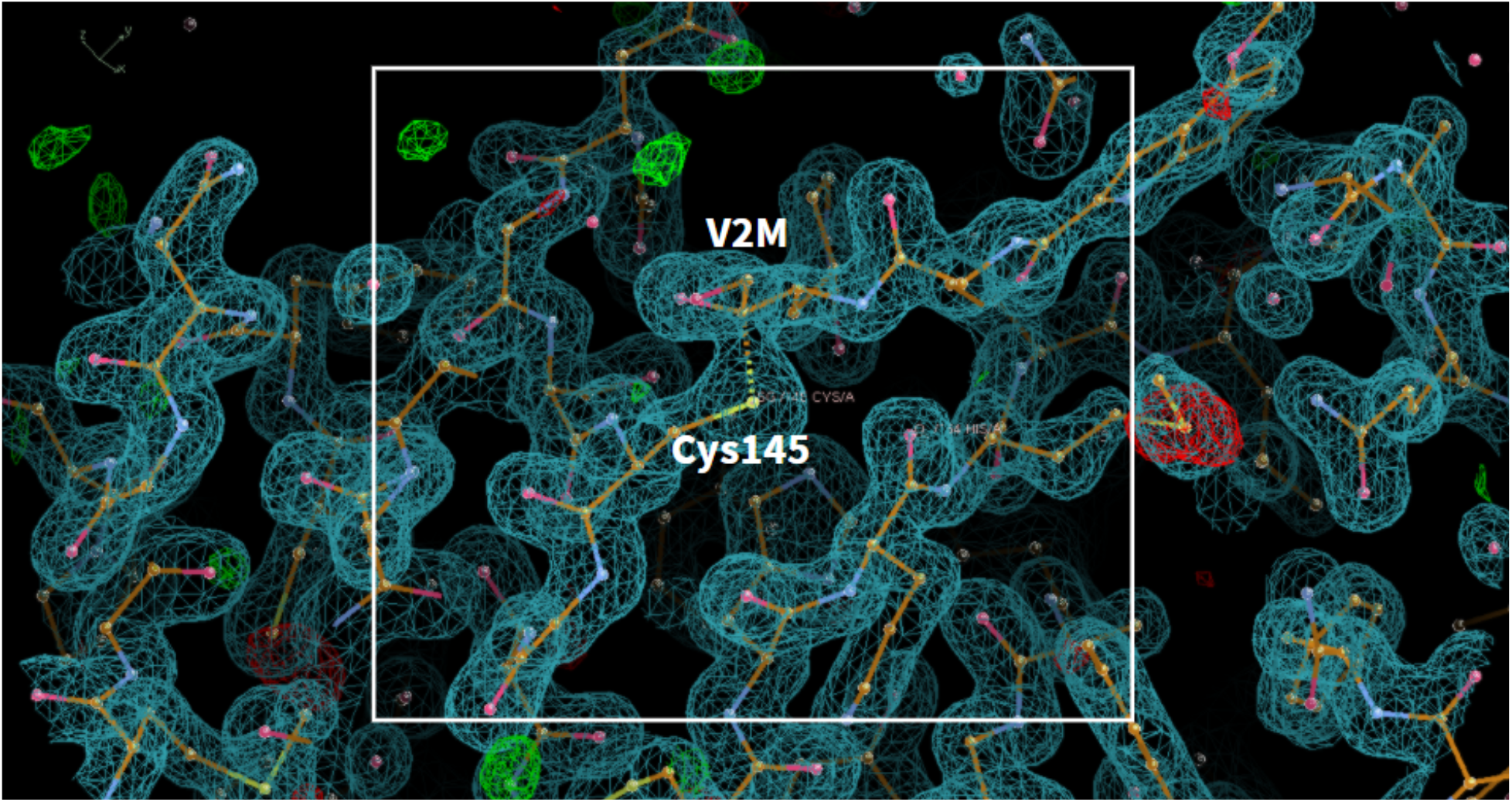
(covalent inhibitor) **PDB ID:** 6XHM **Ligand:** C24 H32 N4 O6(V2M) **Resolution:** 1.41 Å doi: 10.2210/pdb6XHM/pdb **Author(s)**: Hoffman et al. **Deposited:** 2020 June 19 **Released:** 2020 July 08

**Fig. S4.34.**
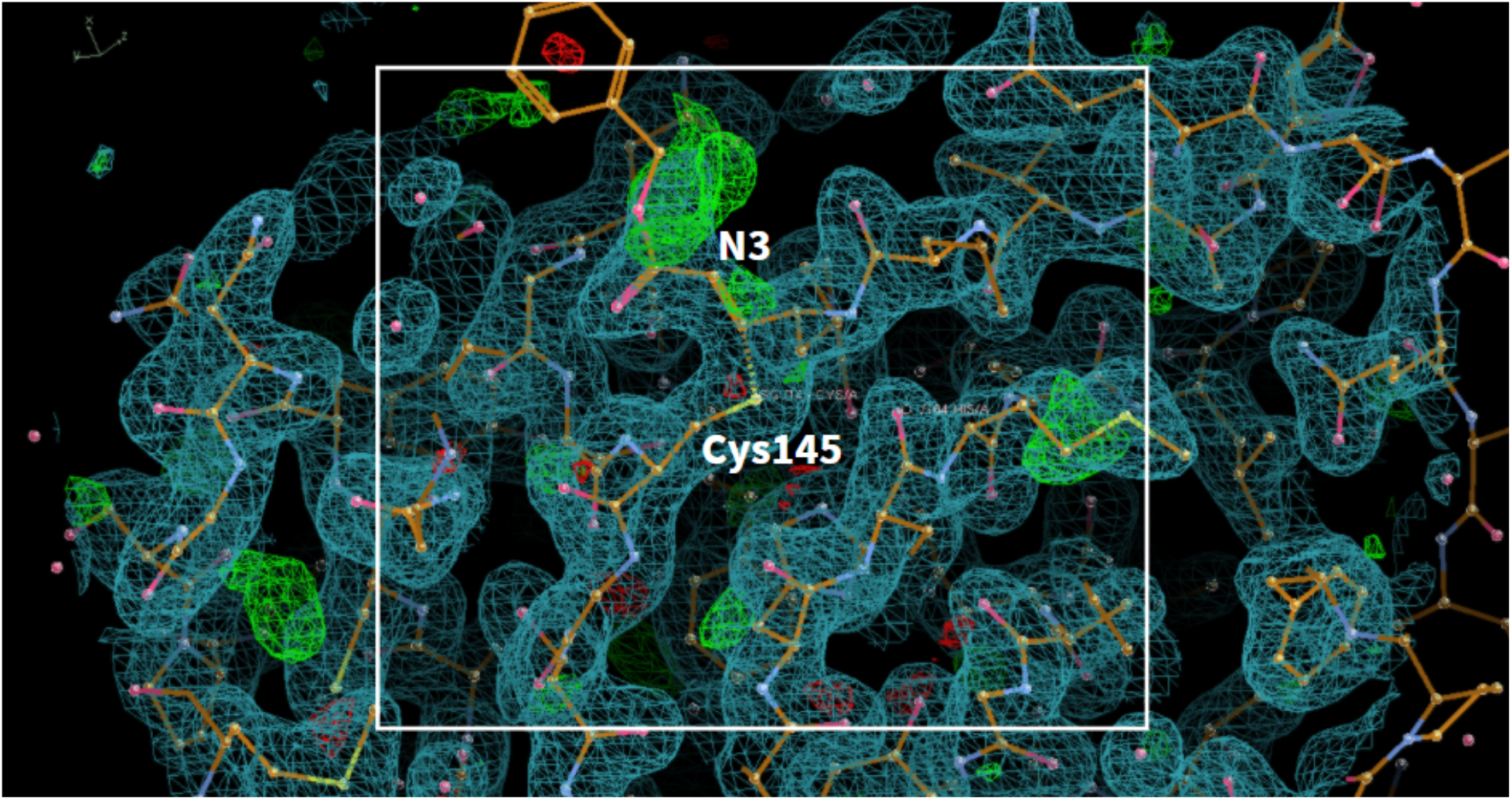
(covalent inhibitor) **PDB ID:** 7BQY **Ligand:** N3 **Resolution:** 1.70 Å doi: 10.2210/pdb7BQY/pdb **Author(s)**: Jin et al. **Deposited:** 2020 March 26 **Released:** 2020 April 22

**Fig. S4.35.**
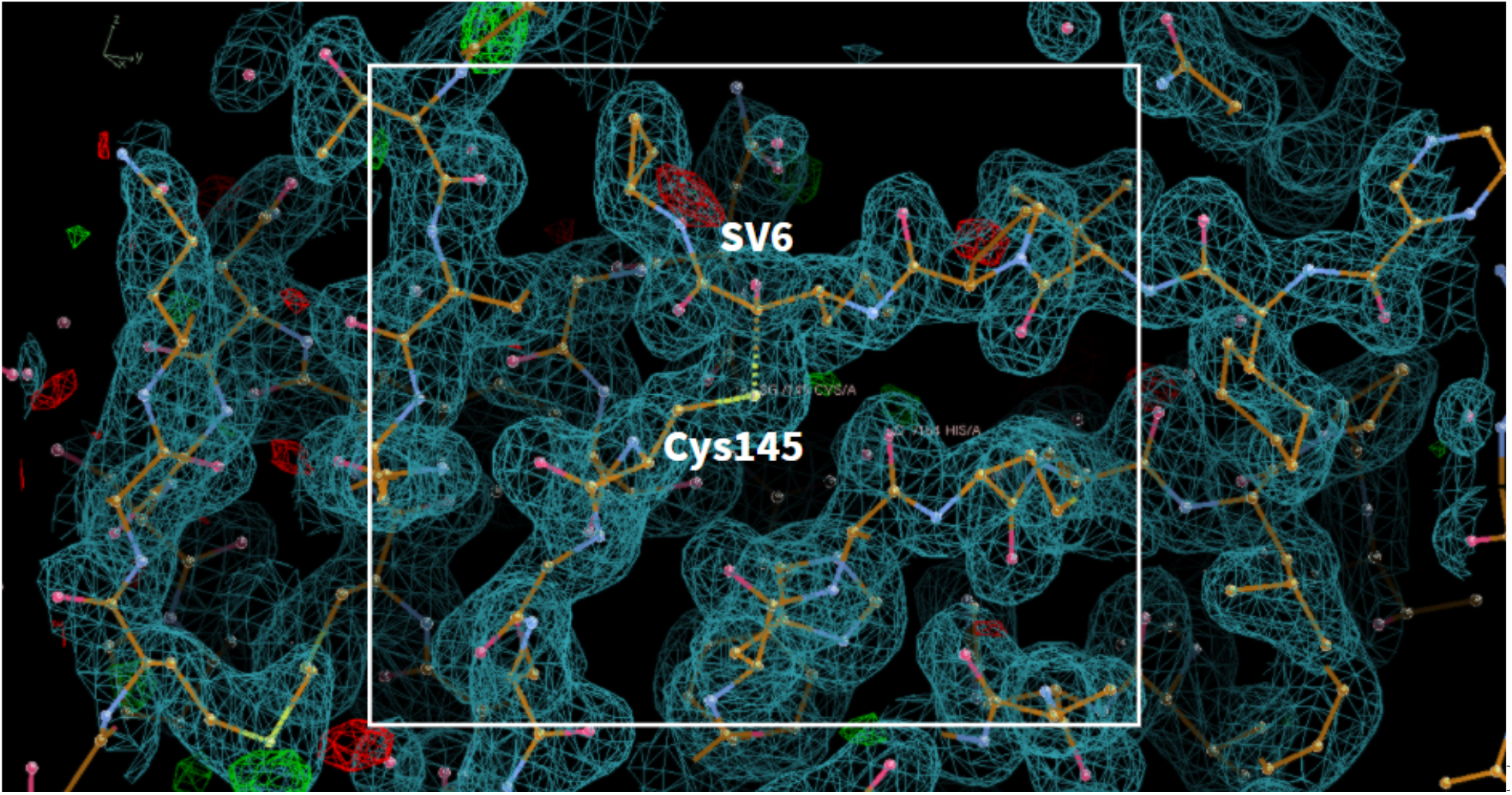
(covalent inhibitor) **PDB ID:** 6XQS **Ligand:**Telaprevir(SV6) **Resolution:** 1.90 Å doi: 10.2210/pdb6XQS/pdb **Author(s)**: Kneller et al. **Deposited:** 2020 July 10 **Released:** 2020 July 22

**Fig. S4.36.**
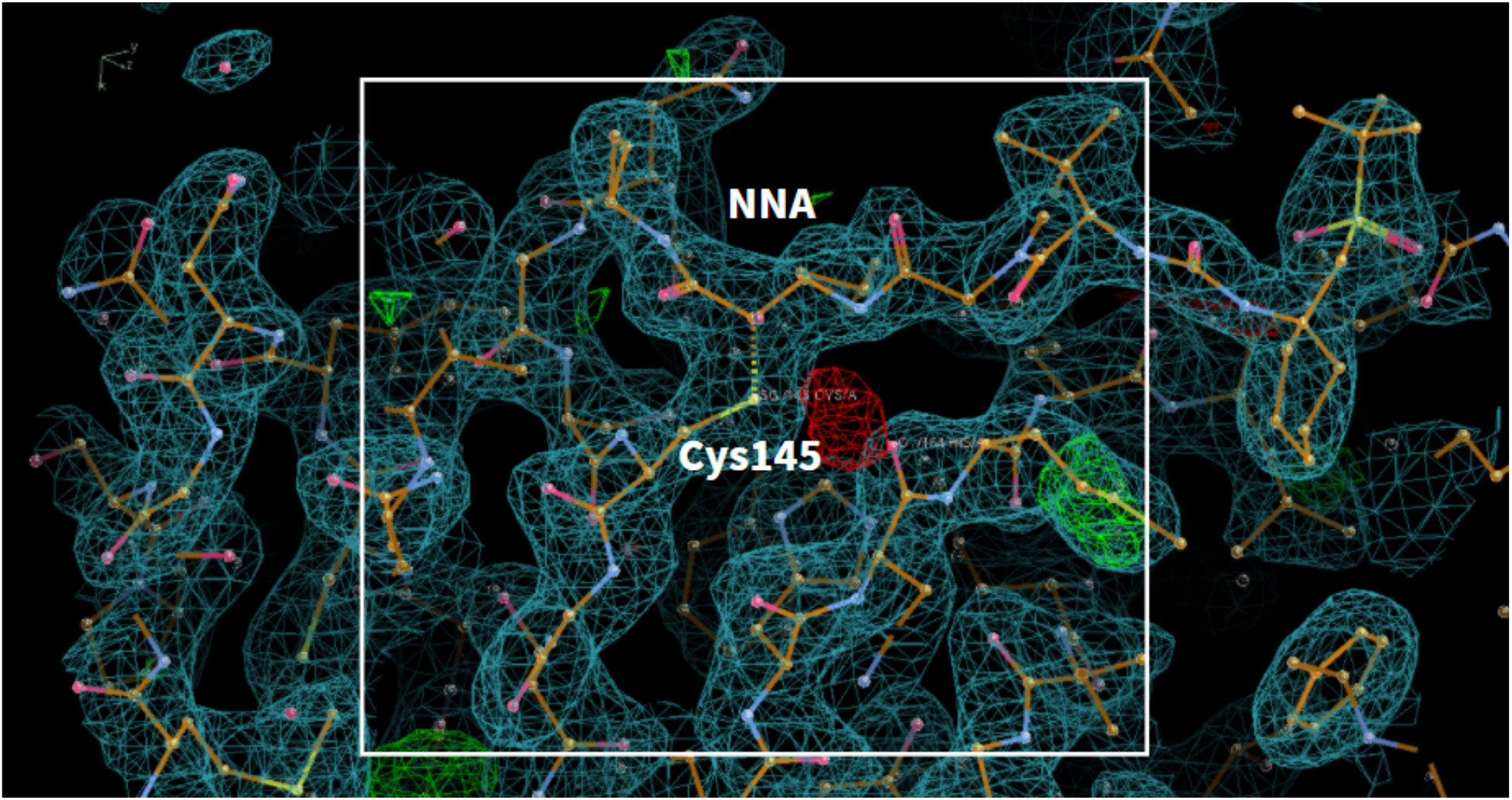
(covalent inhibitor) **PDB ID:** 6XQT **Ligand:** C36 H63 N5 O7 S(NNA) **Resolution:** 2.30 Å doi: 10.2210/pdb6XQT/pdb **Author(s)**: Kneller et al. **Deposited:** 2020 July 10 **Released:** 2020 July 22

**Fig. S4.37.**
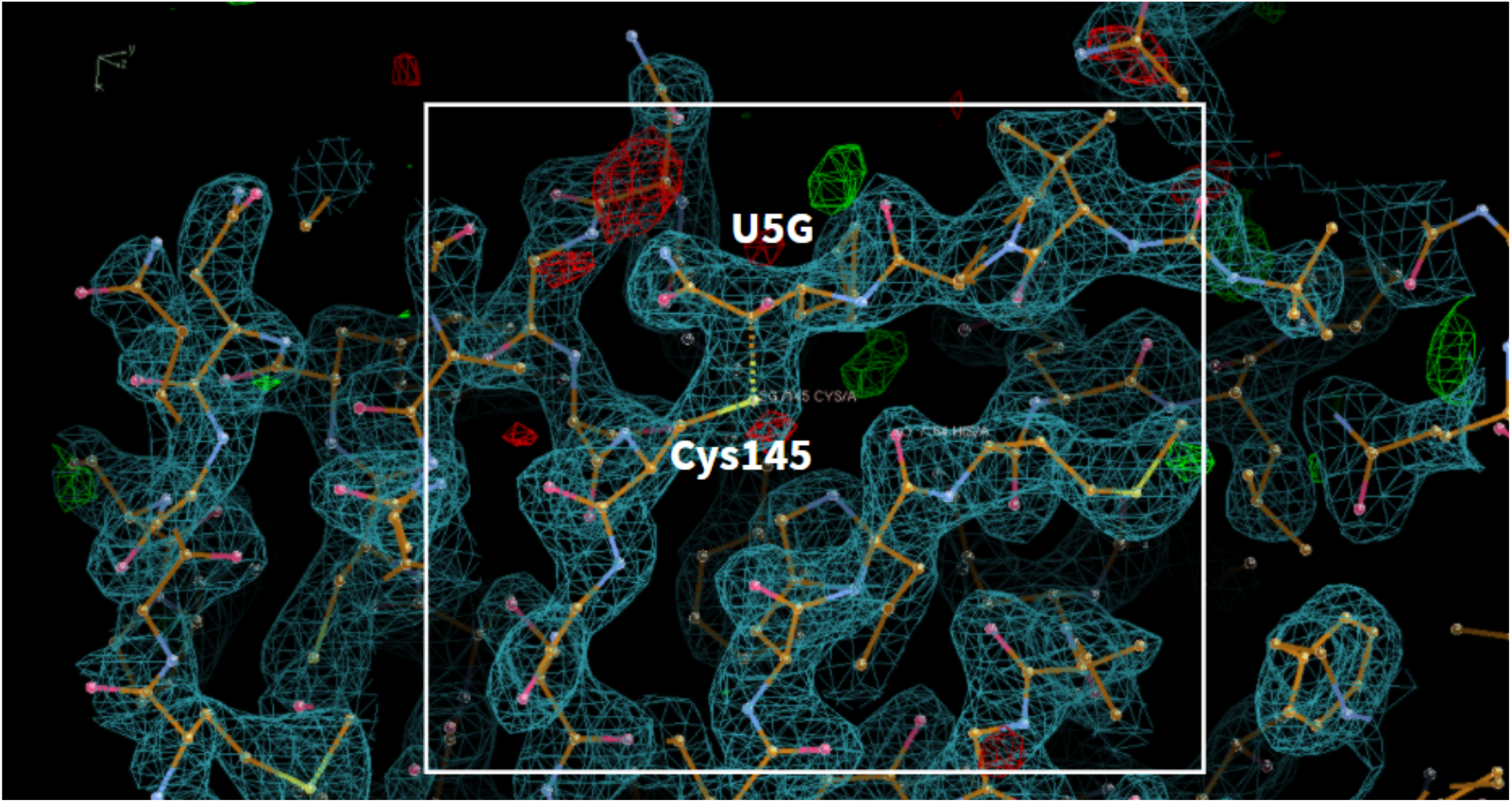
(covalent inhibitor) **PDB ID:** 6XQU **Ligand:** Boceprevir(U5G) **Resolution:** 2.20 Å doi: 10.2210/pdb6XQU/pdb **Author(s)**: Kneller et al. **Deposited:** 2020 July 10 **Released:** 2020 July 22

**Fig. S5.**
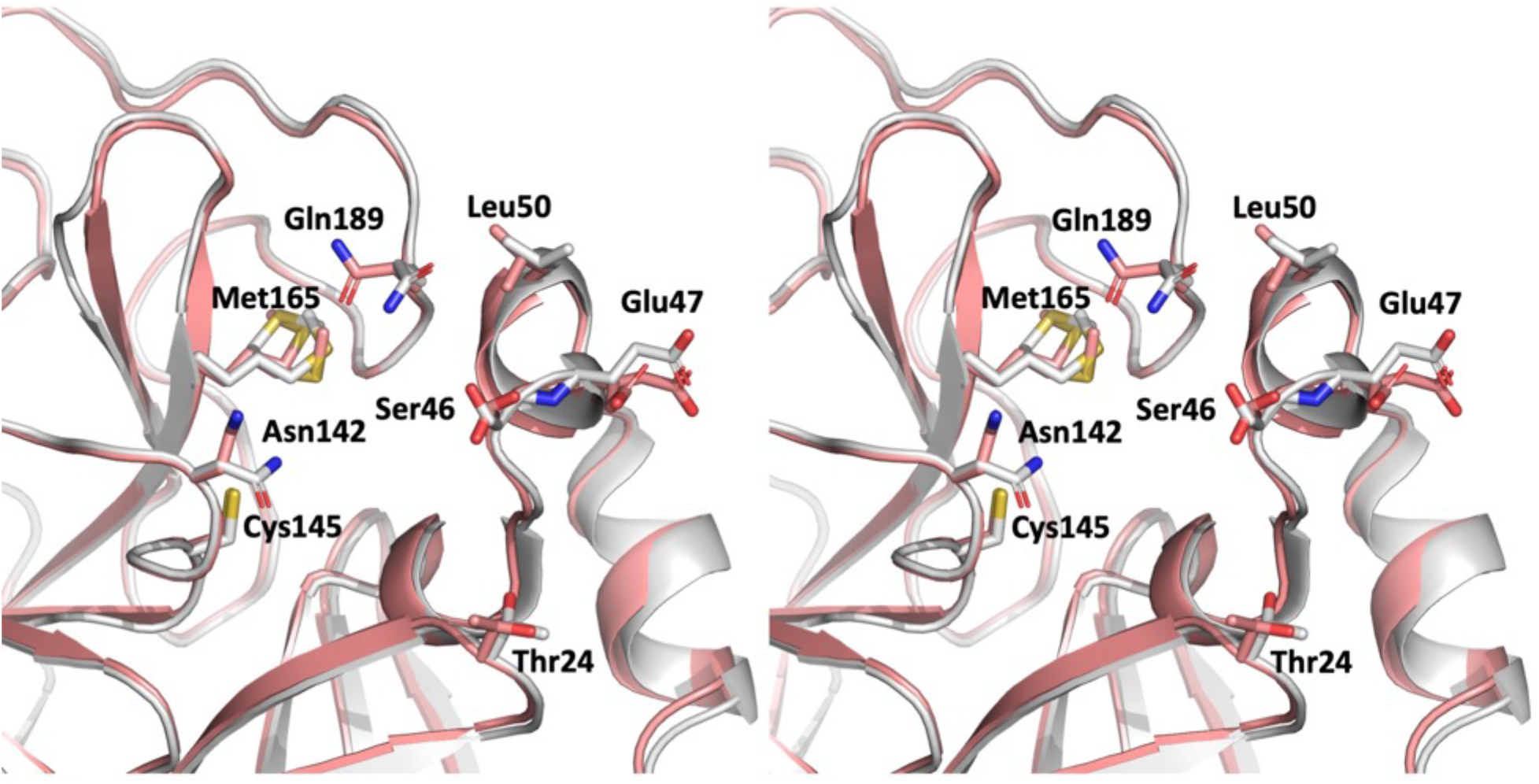
Wall-eye stereo image of the superposition of the active site of SFX structure (PDB ID:7CWB) with ambient-temperature structure (PDB ID: 6WQF). Significant conformational states are observed at the putative inhibitor binding pocket. Residues with altered conformations were indicated by sticks, labeled and their positions were indicated.

**Fig. S6.**
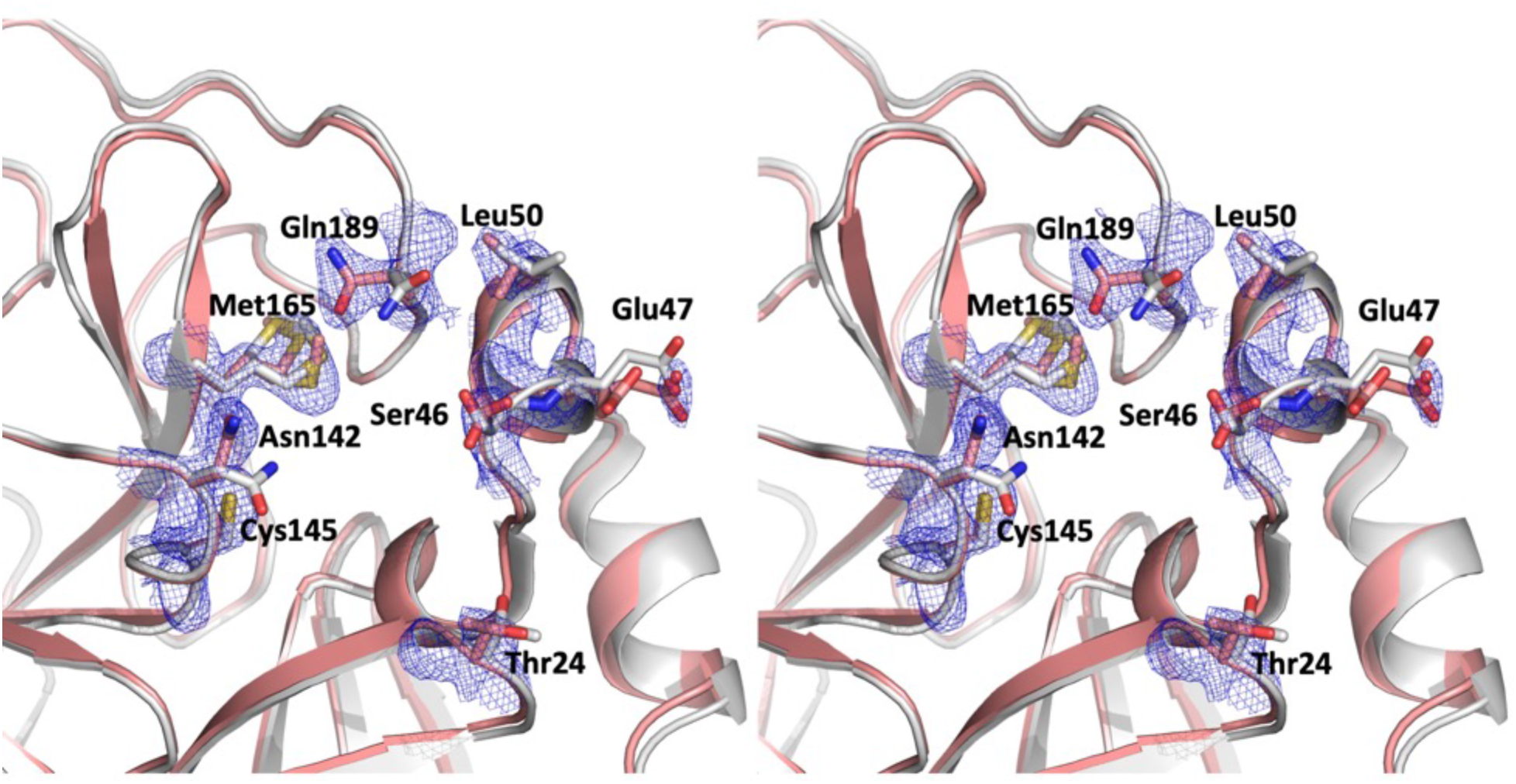
Wall-eye stereo image of composite omit electron density map of active site. Superposition the SFX structure (PDB ID: 7CWB) and ambient-temperature structure (PDB ID: 6WQF). The bias-free composite electron density belongs to residues with altered conformations of SARS-CoV-2 Mpro contoured at 1 sigma level and colored in slate.

**Fig. S7.**
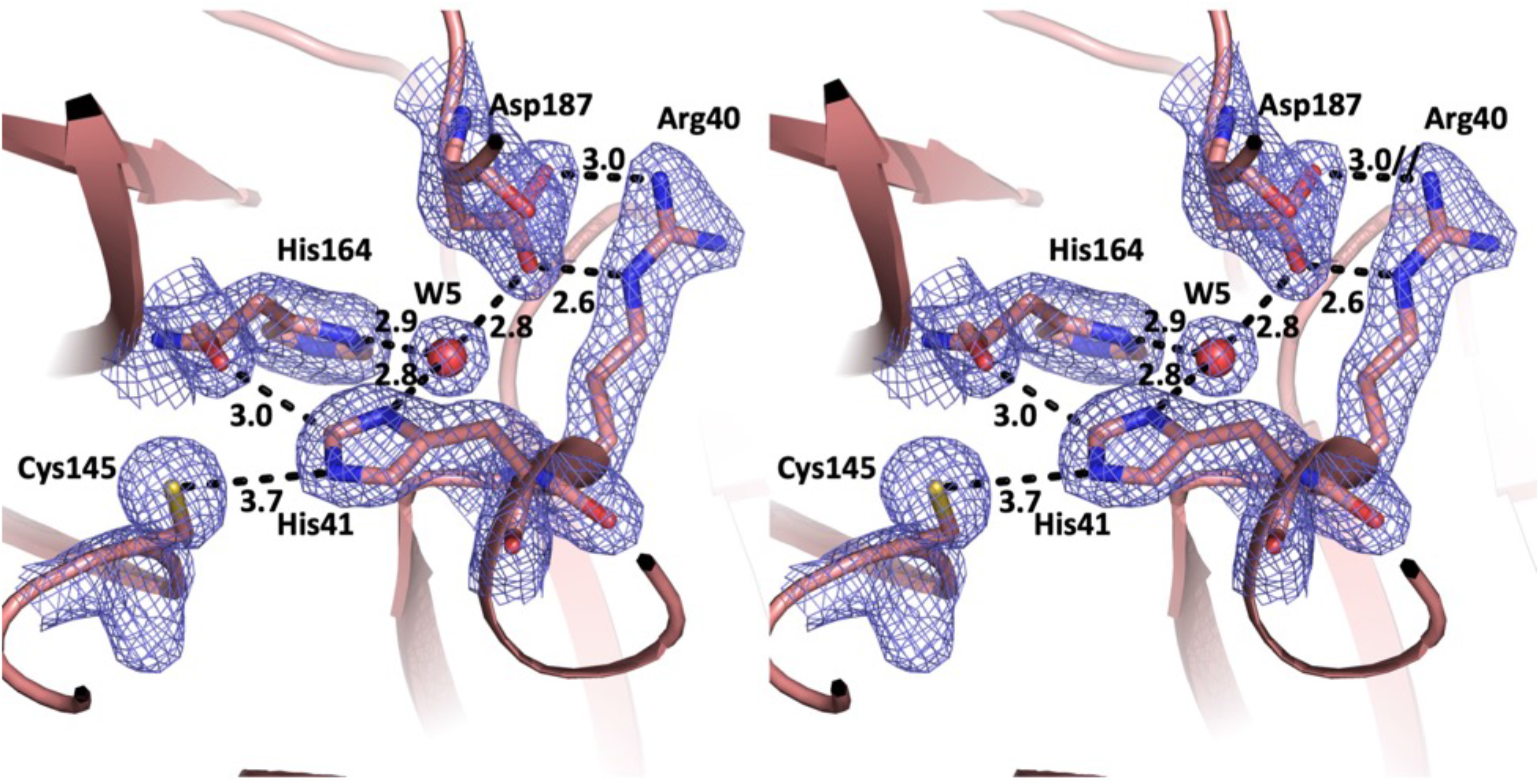
Wall-eye stereo image of SFX structure (PDB ID:7CWB) active site. 2Fo-Fc electron density belonging to the active site residues are contoured in 1 sigma level and colored in slate. H-bonds and other interactions are indicated by dashed lines and distances are given by Angstrom.

**Fig. S8.**
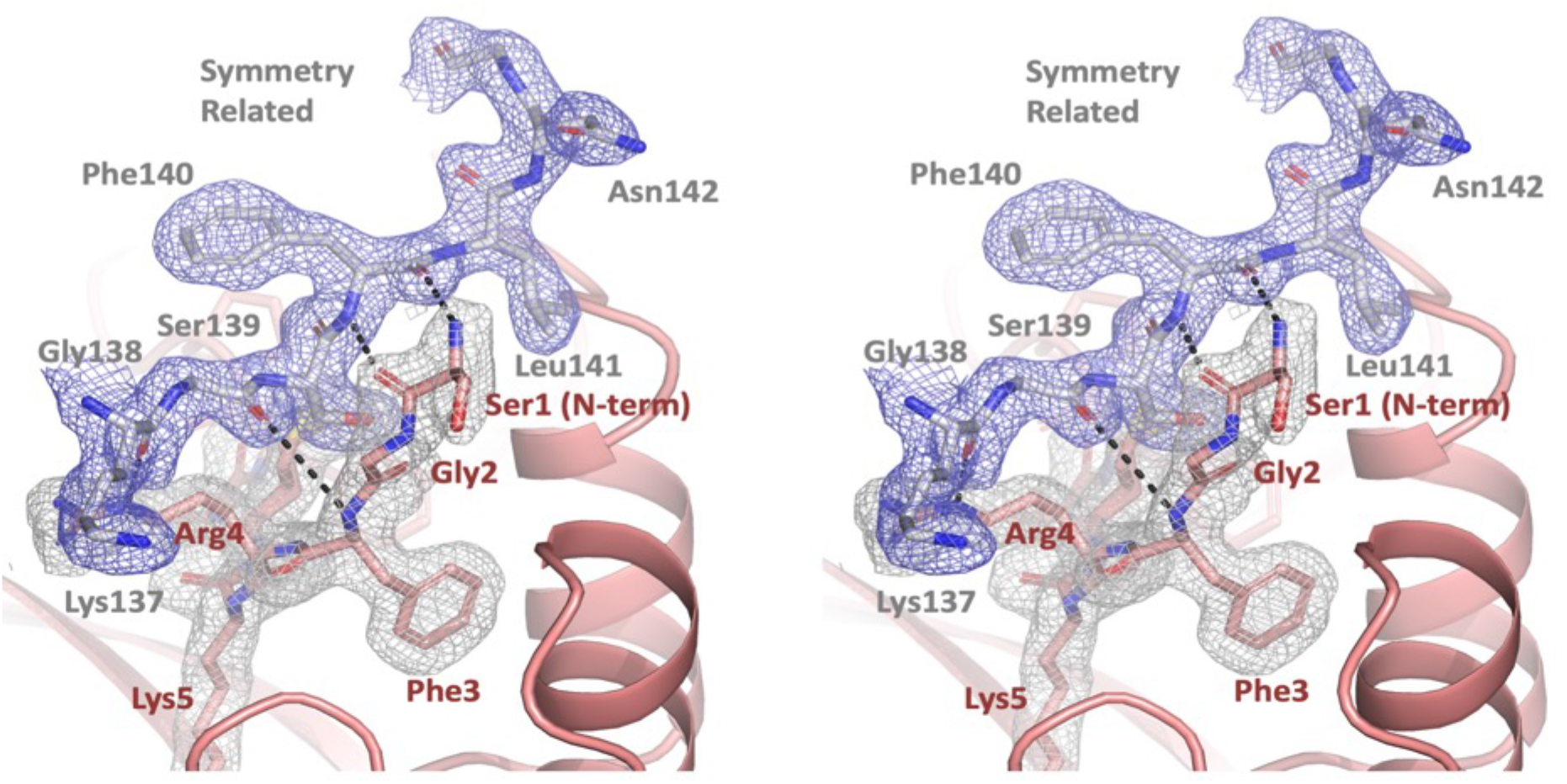
Wall-eye stereo image of key crystal contacts of C121 crystal form. The 2Fo-Fc electron density belongs to the N-terminal region of SARS-CoV-2 main protease contoured at 1 sigma level and colored in gray. Regions of the symmetry related molecule colored in gray, 2Fo-Fc electron density contoured in 1 sigma level and colored in slate. H-bonds are indicated by dashed lines. Elimination of these H-bonds were essential to obtain the second crystal form in P2_1_2_1_2_1_ space group with more open inhibitor binding pocket for future soaking studies of the putative main protease inhibitors.

**Fig. S9.**
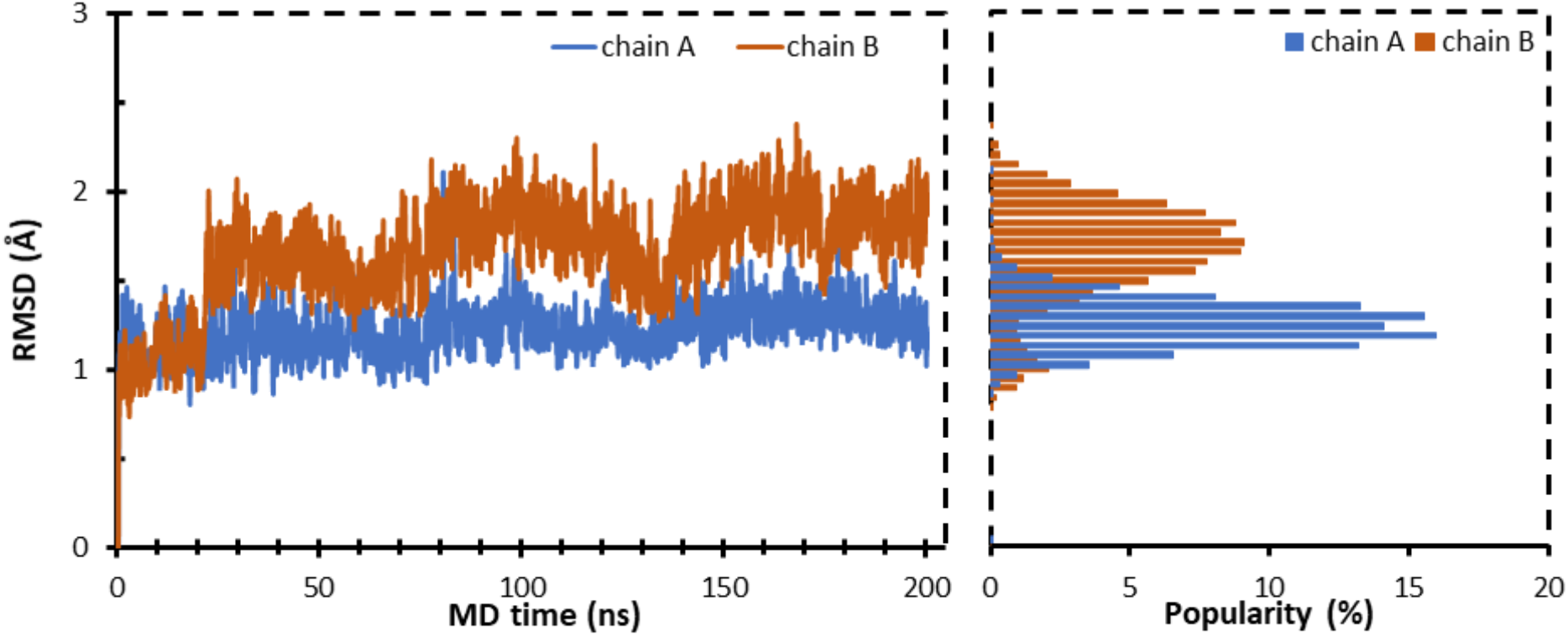
Changes in RMSD observed in Cα atoms during MD simulations with respect to initial trajectory frame for both chains of 7CWB.

**Fig. S10.**
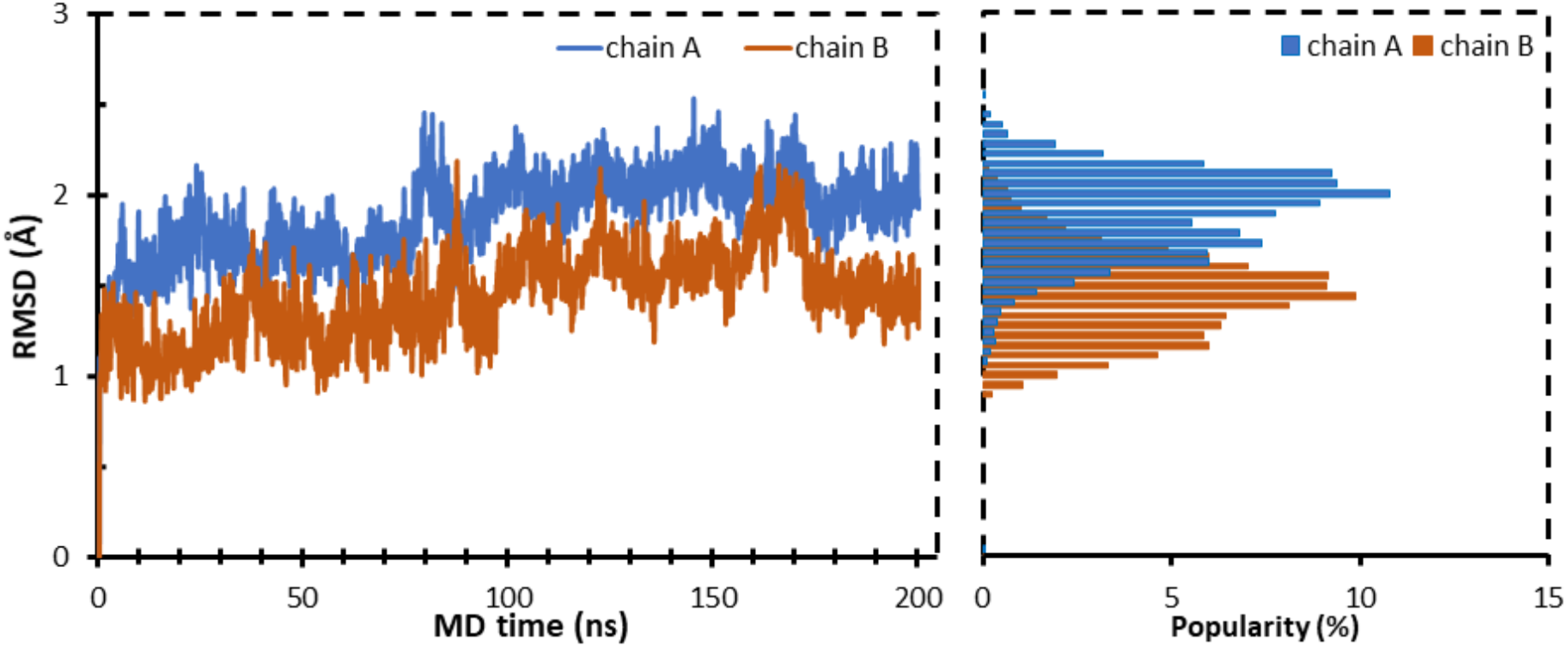
Changes in RMSD observed in Cα atoms during MD simulations with respect to initial trajectory frame for both chains of 7CWC.

**Fig. S11.**
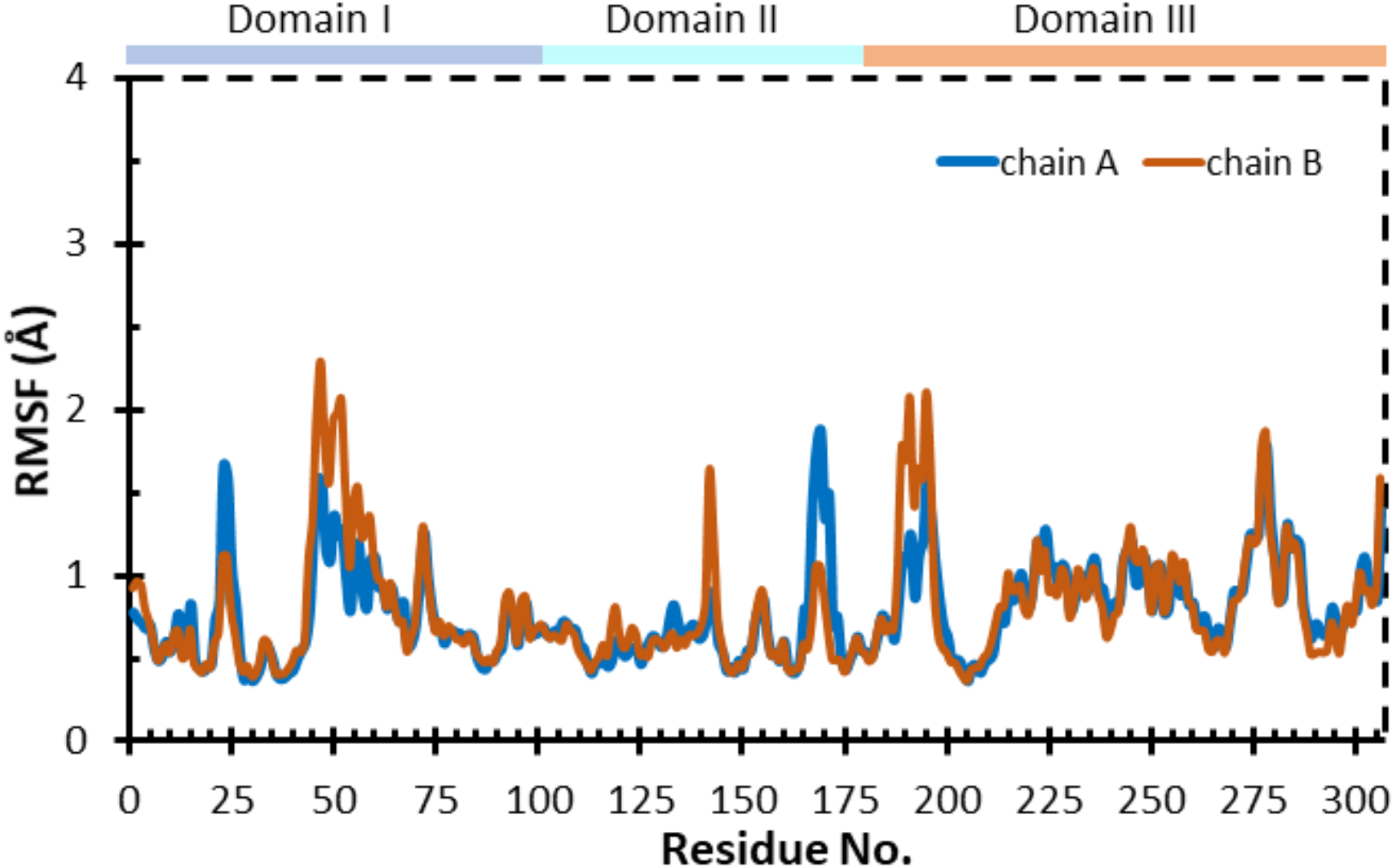
RMSF changes observed in Cα atoms during MD simulations for both chains of 7CWB with domains I, II and III also displayed above the graph.

**Fig. S12.**
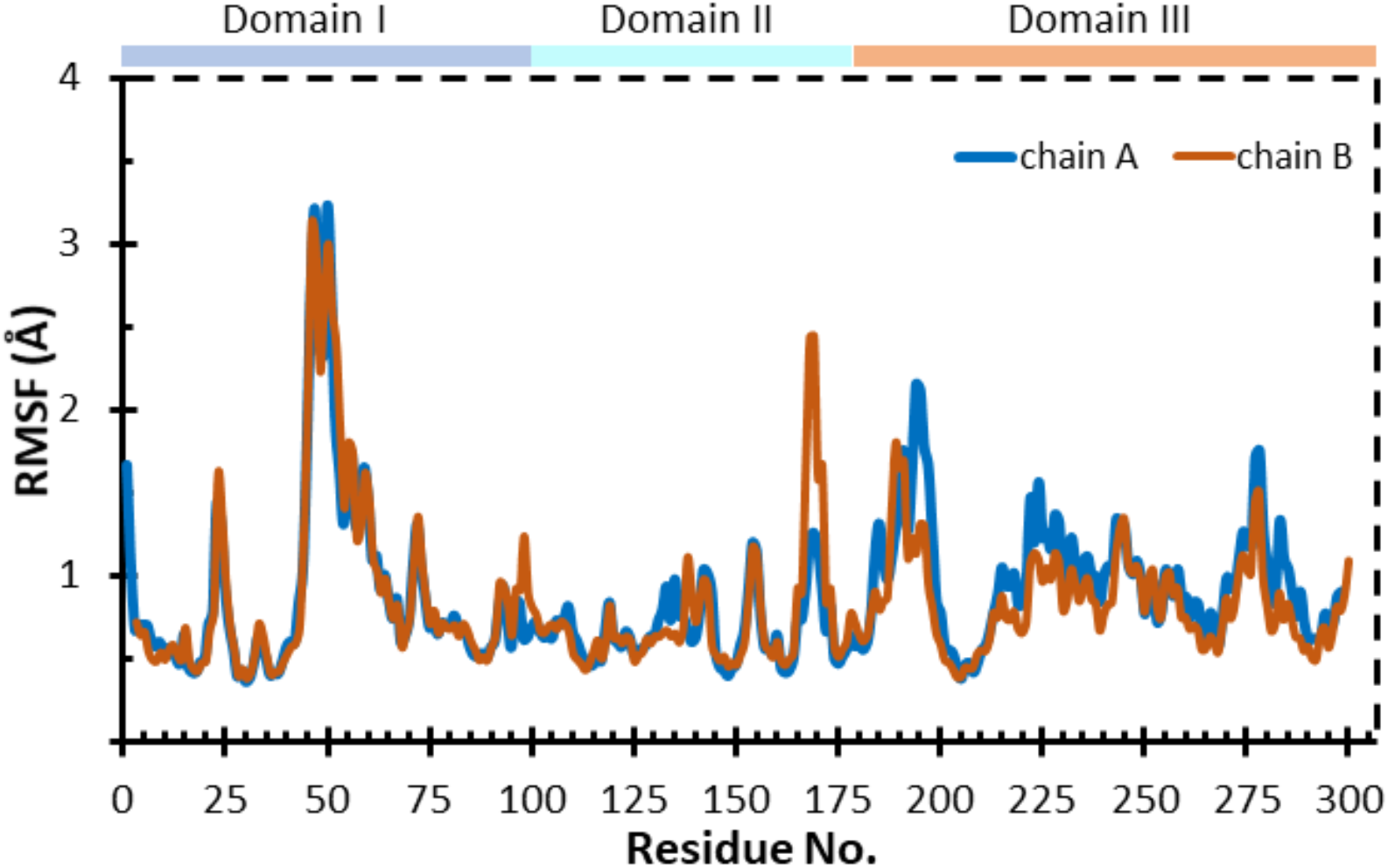
RMSF changes observed in Cα atoms during MD simulations for both chains of 7CWC with domains I, II and III also displayed above the graph.

**Fig. S13.**
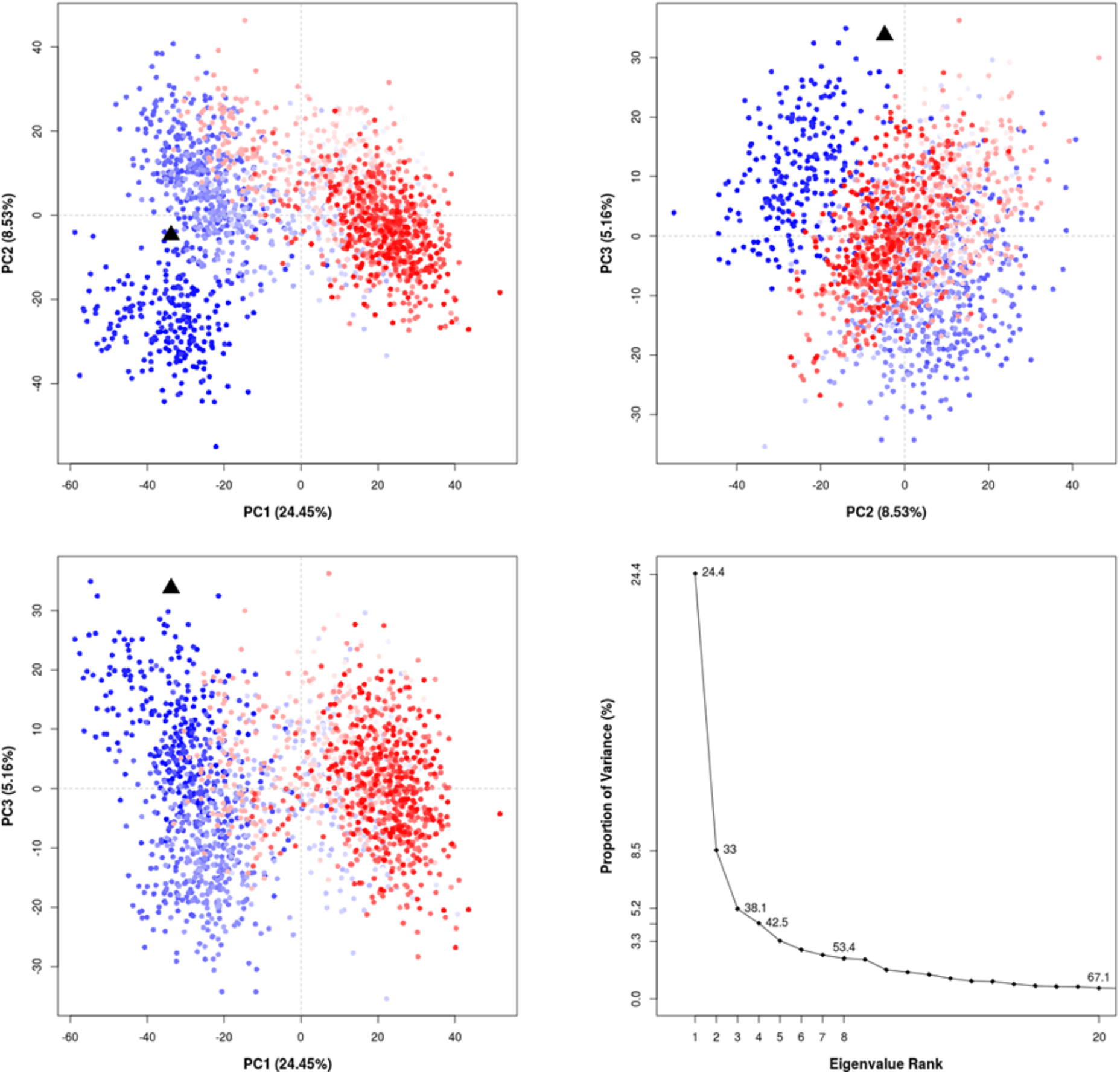
Projection of MD trajectory frames onto subspaces defined by the first three largest PCs as well as distribution of variance observed for eigenvalues for 7CWB. Instantaneous conformations (i.e. trajectory frames) colored from blue to red in order of trajectory time (200 ns). The black triangle represents the initial crystal structure conformation projected onto specified PCs.

**Fig. S14.**
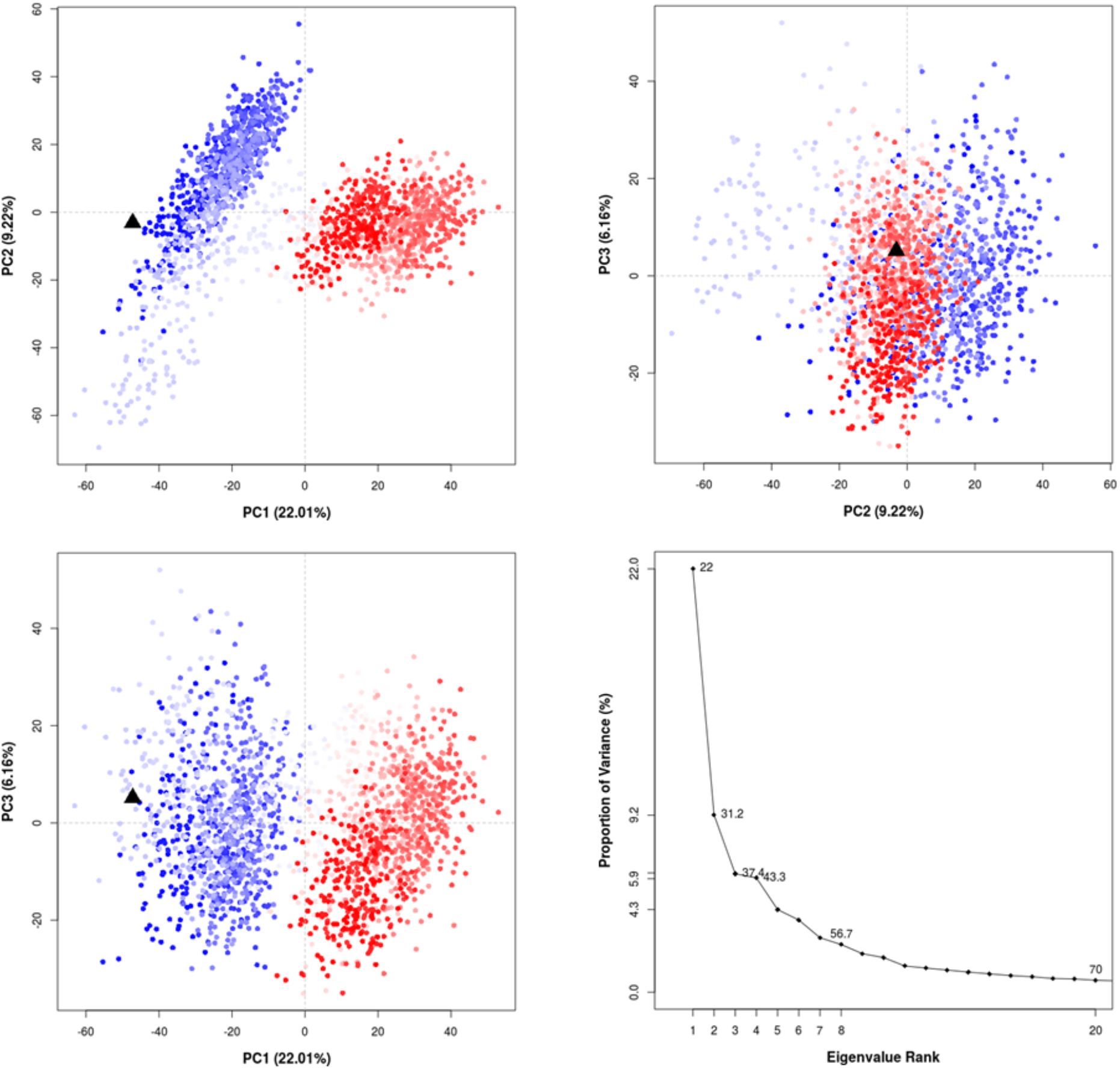
Projection of MD trajectory frames onto subspaces defined by the first three largest PCs as well as distribution of variance observed for eigenvalues for 7CWC. Instantaneous conformations (i.e. trajectory frames) colored from blue to red in order of trajectory time (200 ns). The black triangle represents the initial crystal structure conformation projected onto specified PCs.

**Fig. S15.**
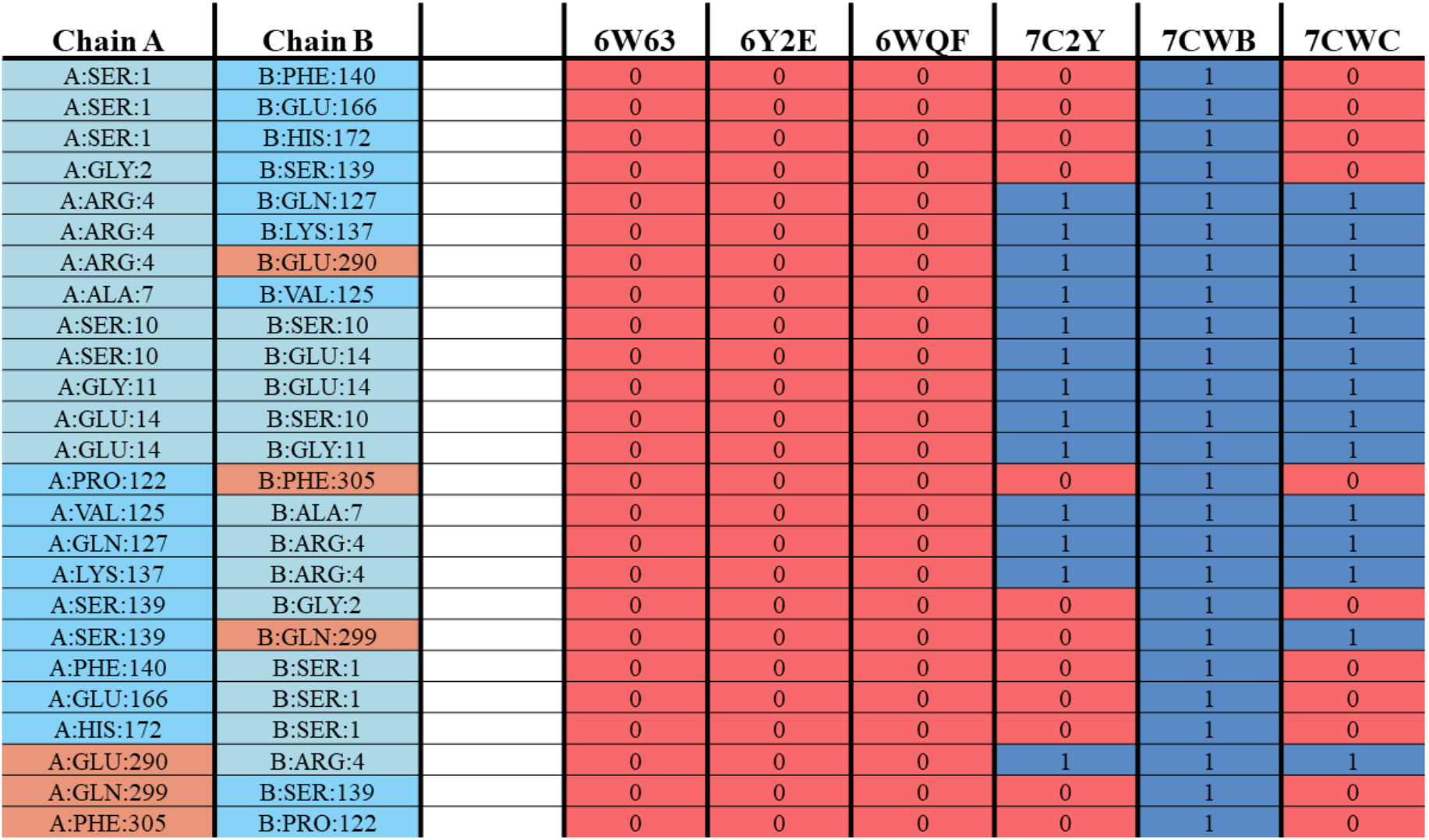
Comparison of the crystal structures of Mpro interfaces with respect to the H-bonds. Domain I residues depicted in light blue, domain II and domain III depicted in pale cyan and dark salmon, respectively. Blue color shows hydrogen bonding, lines with red ones lack hydrogen bonding.

**Fig. S16.**
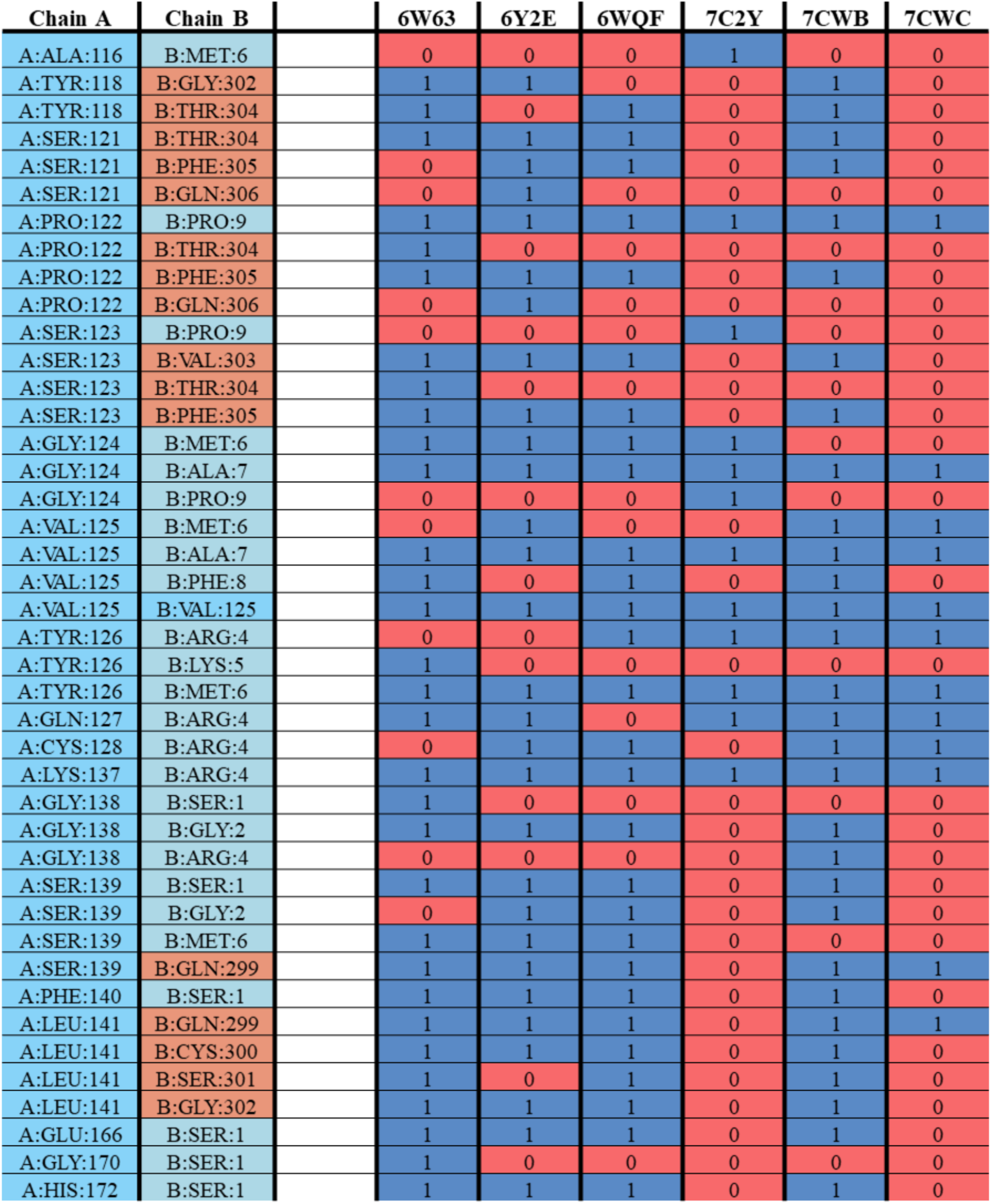
Evaluation of the crystal structures of Mpro interfaces with respect to the all possible interactions (salt bridges, pi-cation, pi-pi stacking, T-stacking, van der Waals (vdW), and H-bonds).

**Fig. S17.**
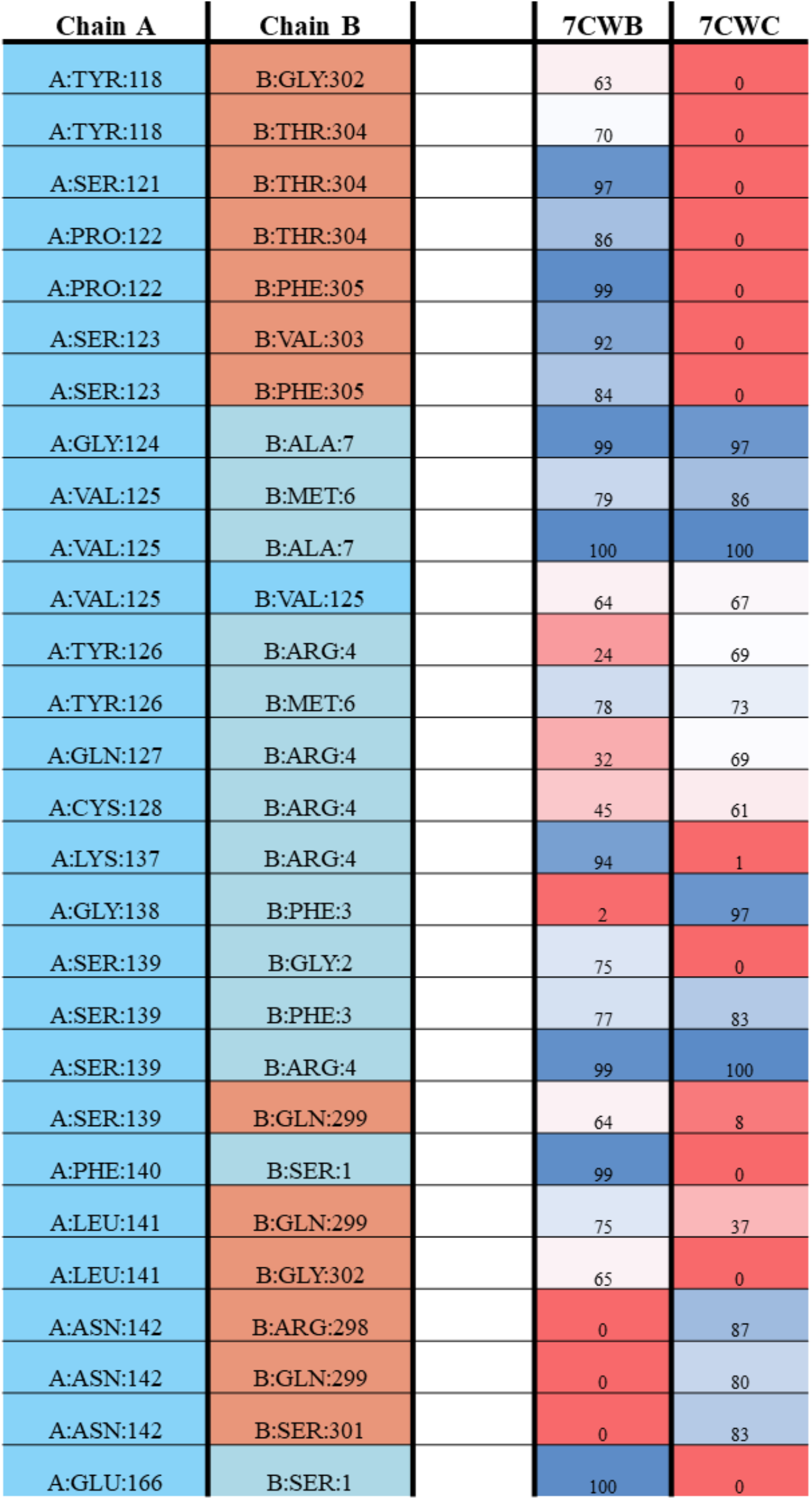
Dynamic valuation of the 7CWB and 7CWC dimer interfaces, all possible interactions were taken into account. Interaction propensity depicted as BWR color heatmap (percentages were given respect to their).

**Fig. S18.**
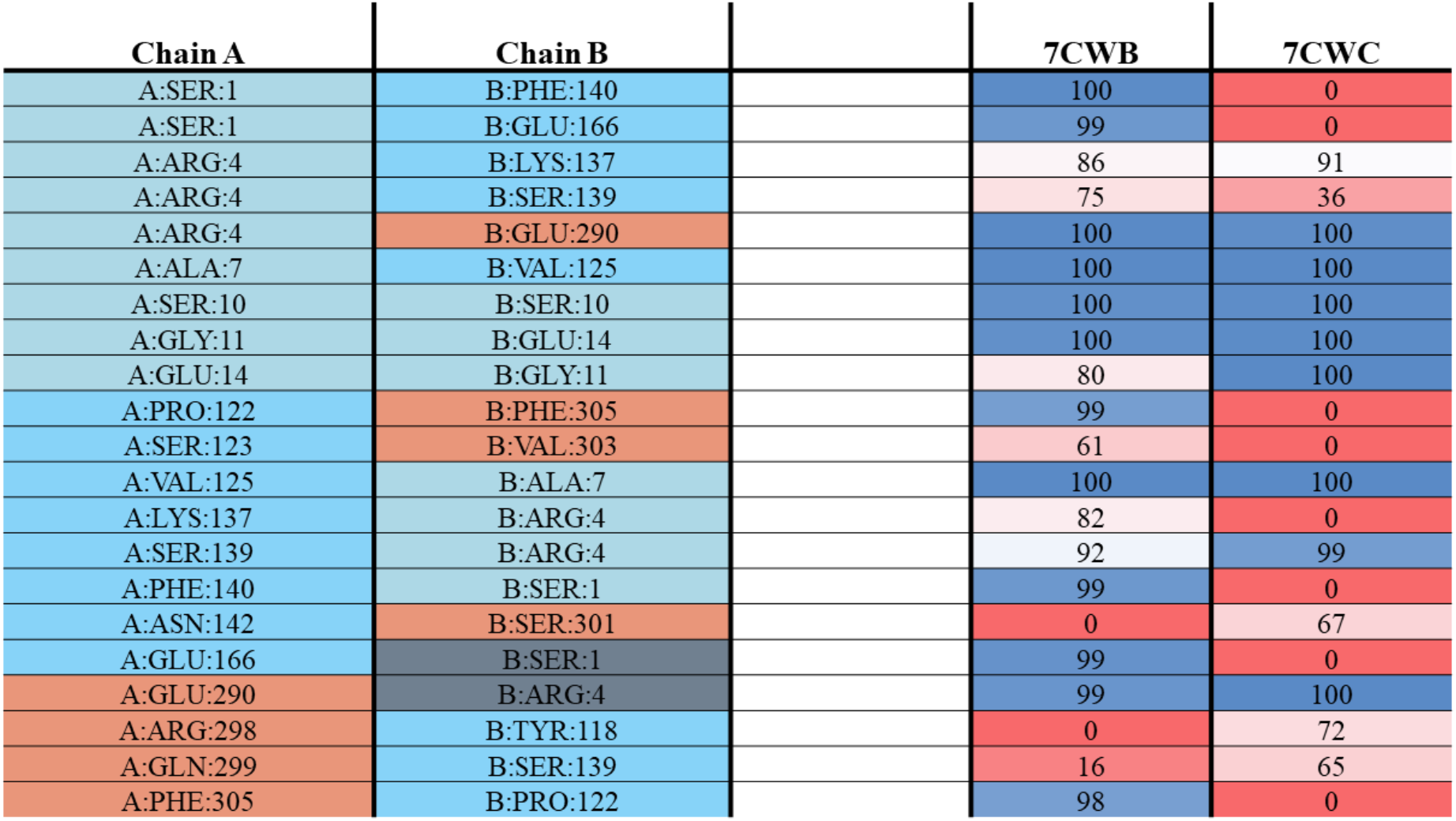
Dynamic evaluation of dimer interfaces of 7CWB, and 7CWC. Only H-bonds were taken into account (0 shows no interaction was observed during MD simulations).

**Fig. S19.**
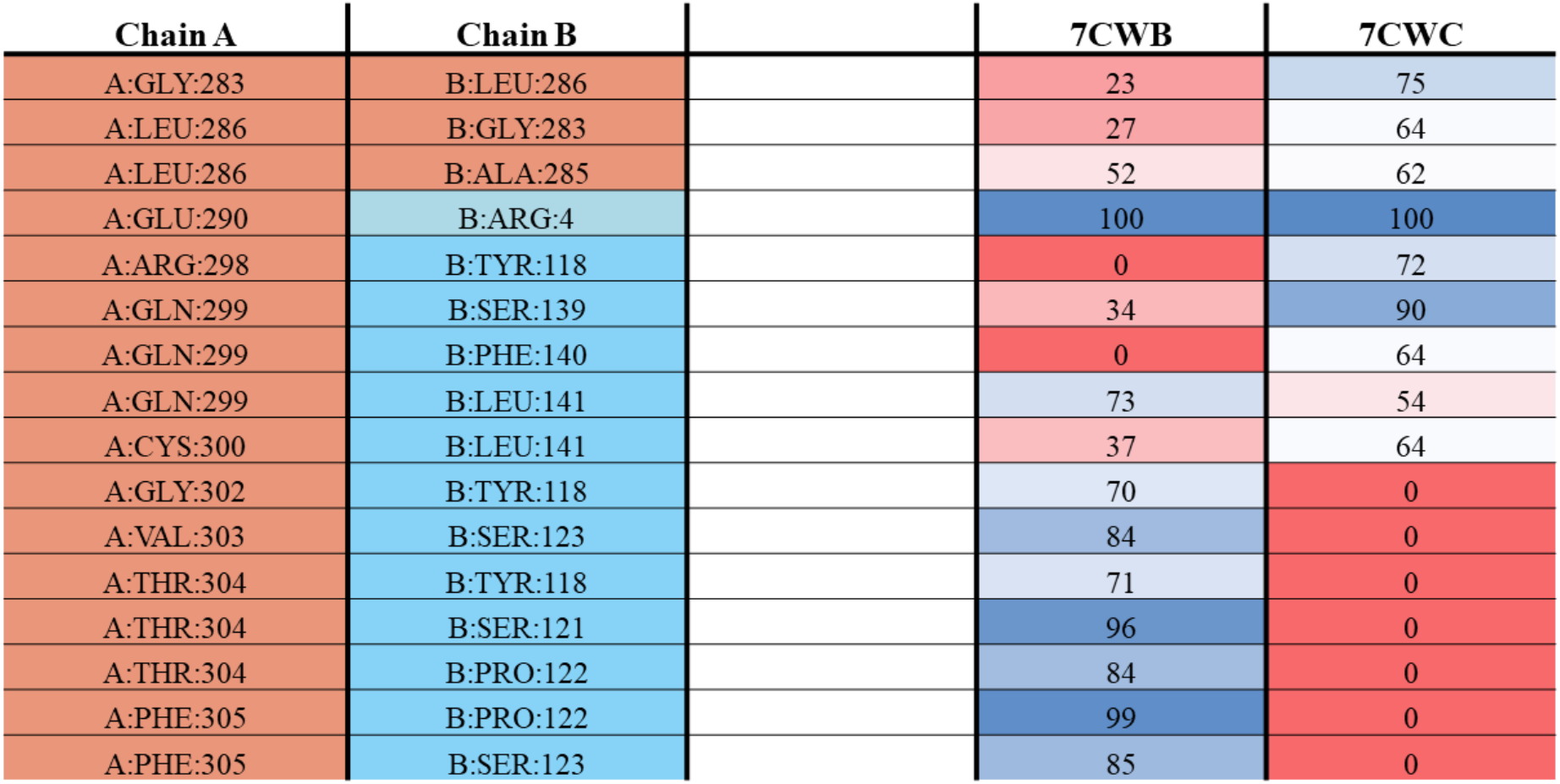
Dynamic evaluation of dimer interfaces of 7CWB, and 7CWC. All possible interactions were taken into account.

**Fig. S20.**
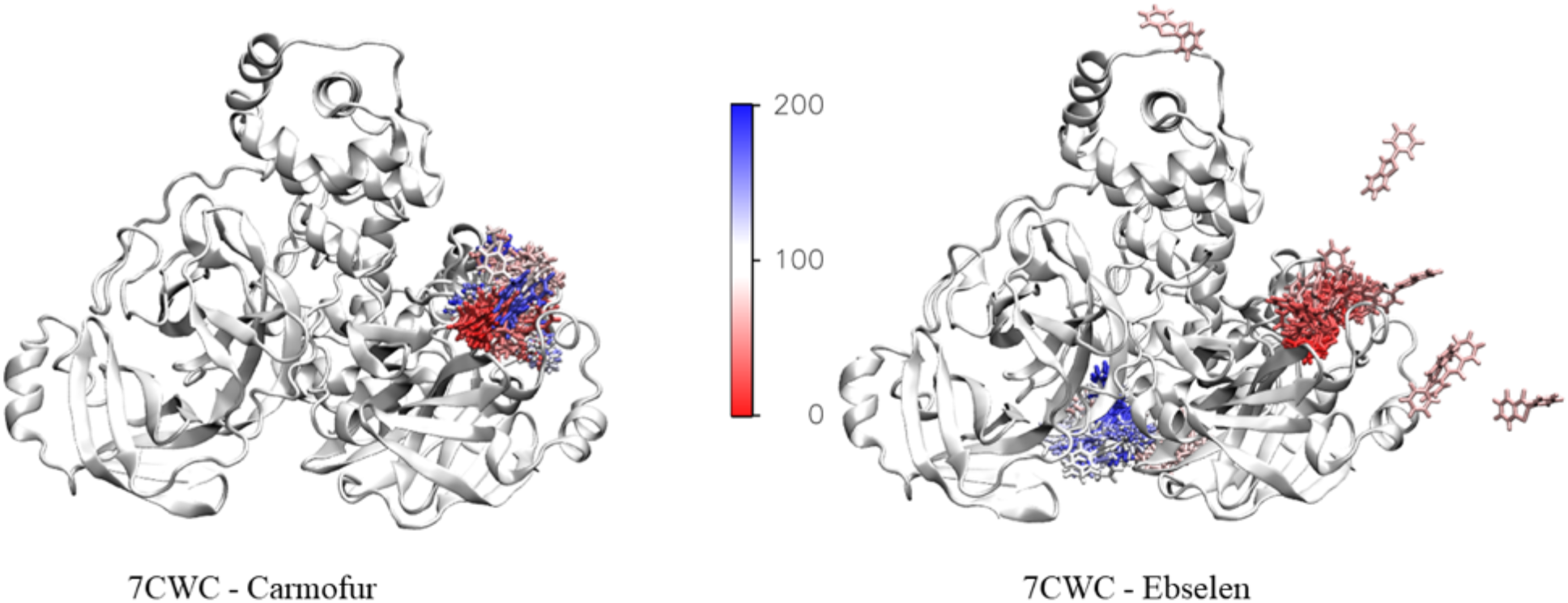
Binding poses of Carmofur (left) and Ebselen (right) at the 7CWC throughout the MD simulations.

**Fig. S21.**
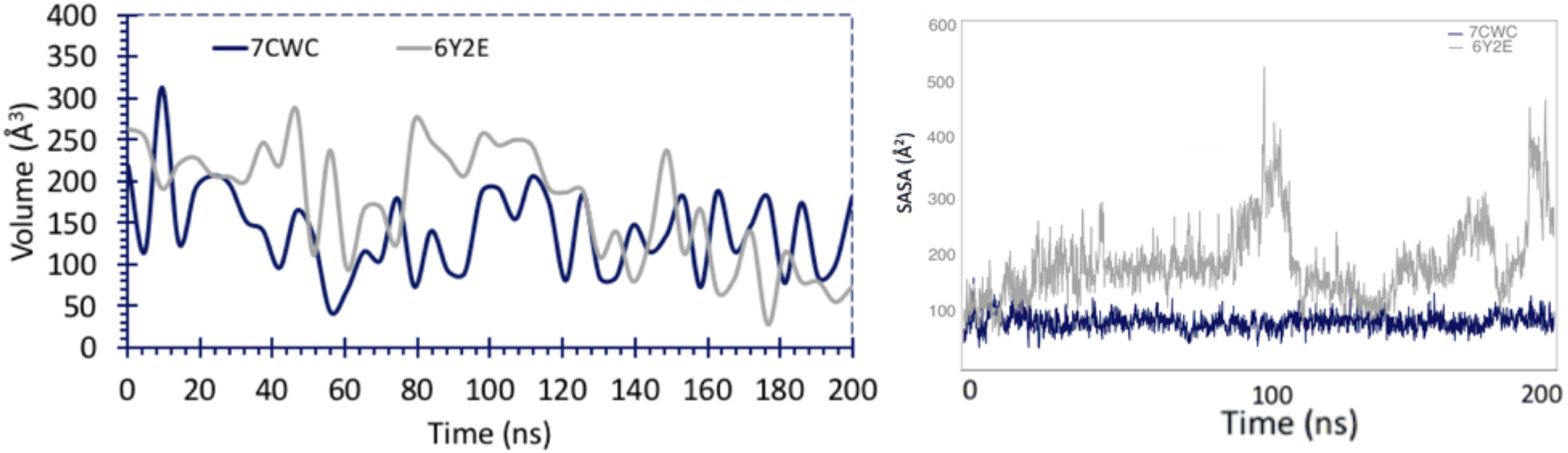
Binding pocket volume (left) and SASA (right) comparison of 7CWC and 6Y2E throughout the MD simulations of Tideglusib bound 7CWC and 6Y2E structures.

**Fig. S22.**
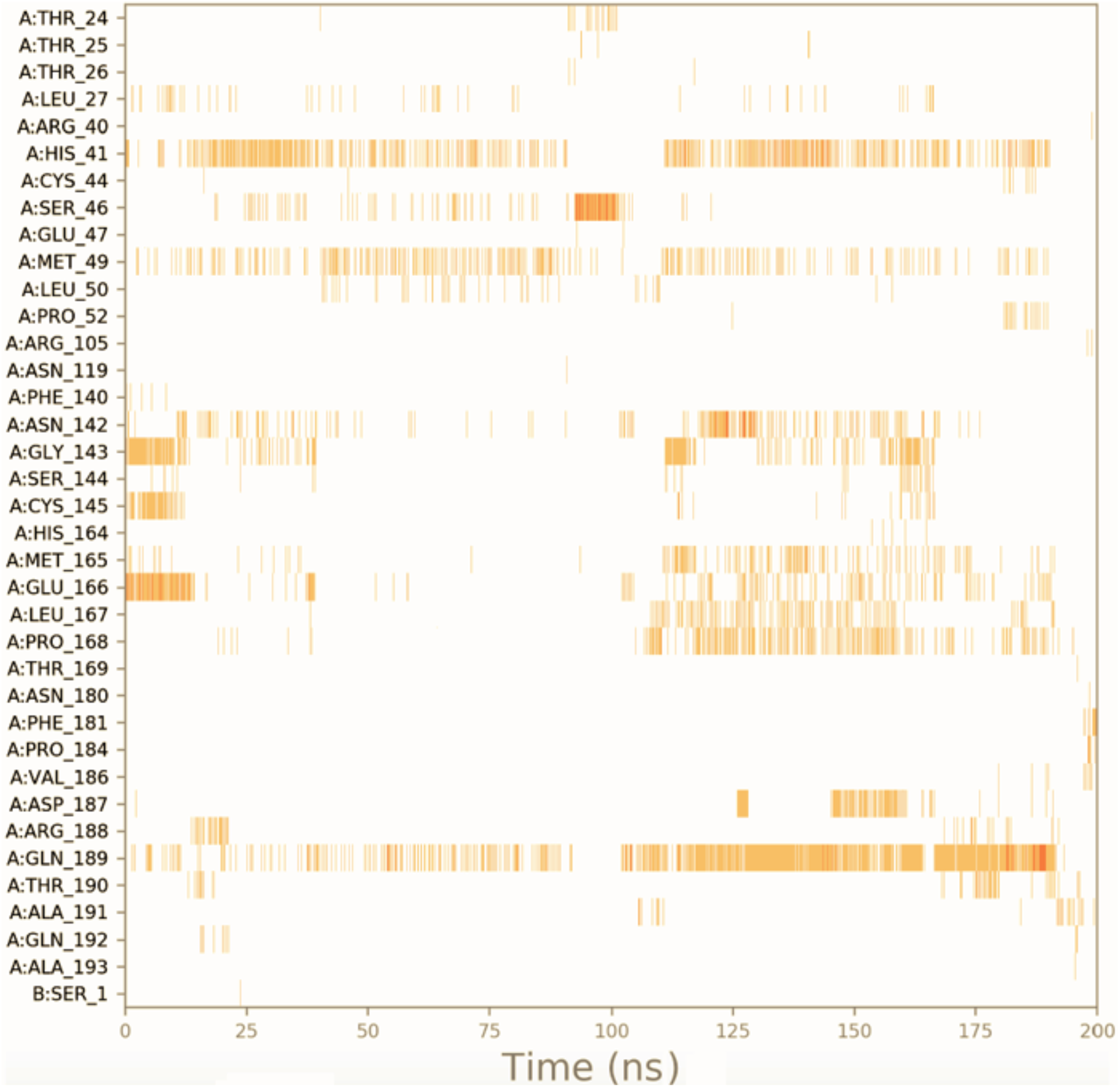
Timeline contacts of Tideglusib at the binding pocket of 6Y2E.

**Fig. S23.**
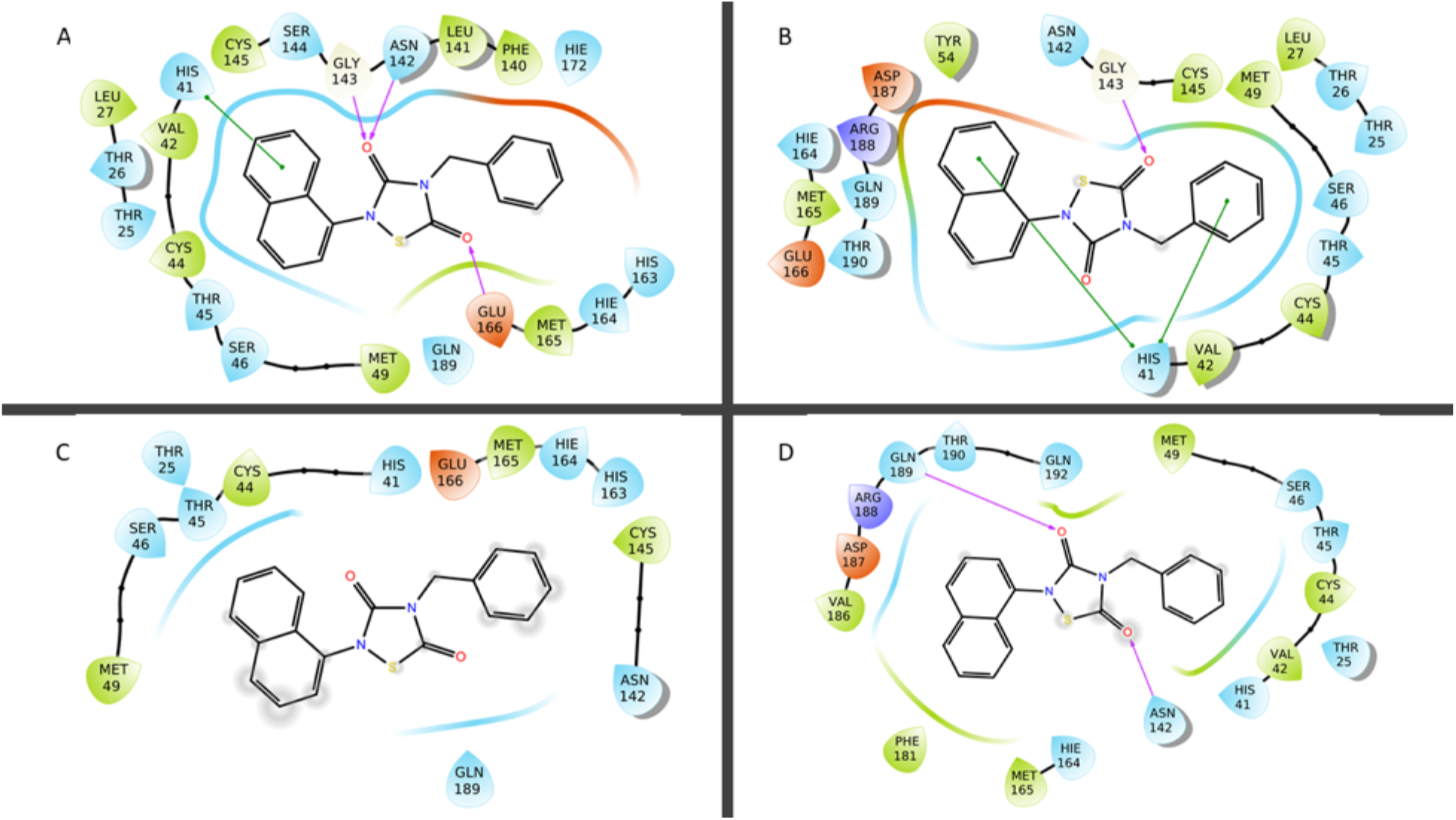
Comparison of the IFD poses of Tideglusib at the binding pocket of A) 7CWB and B) 6WQF. Representative poses that are obtained from MD simulations **C)** 7CWB and **D)** 6WQF.

**Fig. S24.**
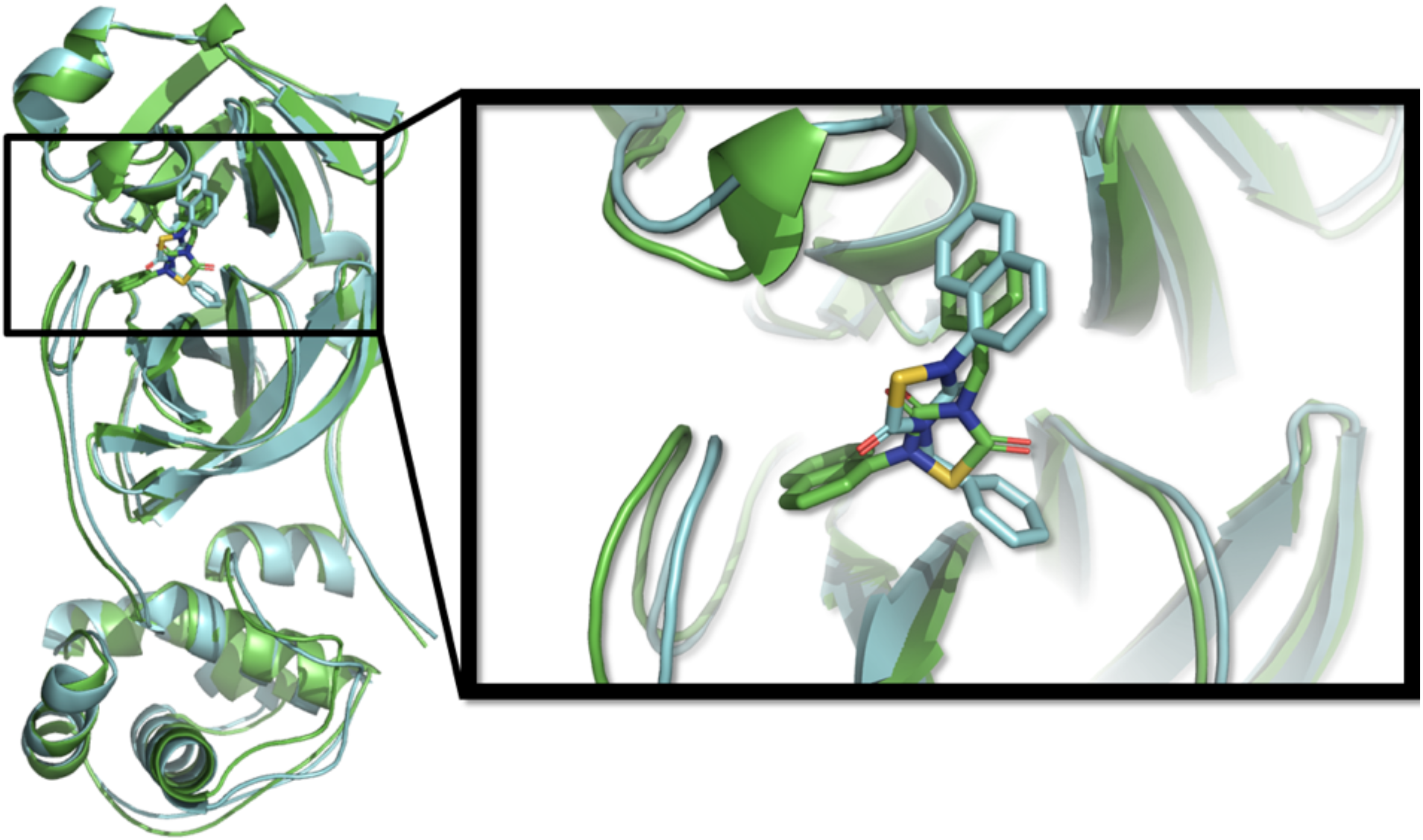
Alignment of the representative structures of Tideglusib complexes (7CWB, cyan, and 6WQF, green).

**Fig. S25.**
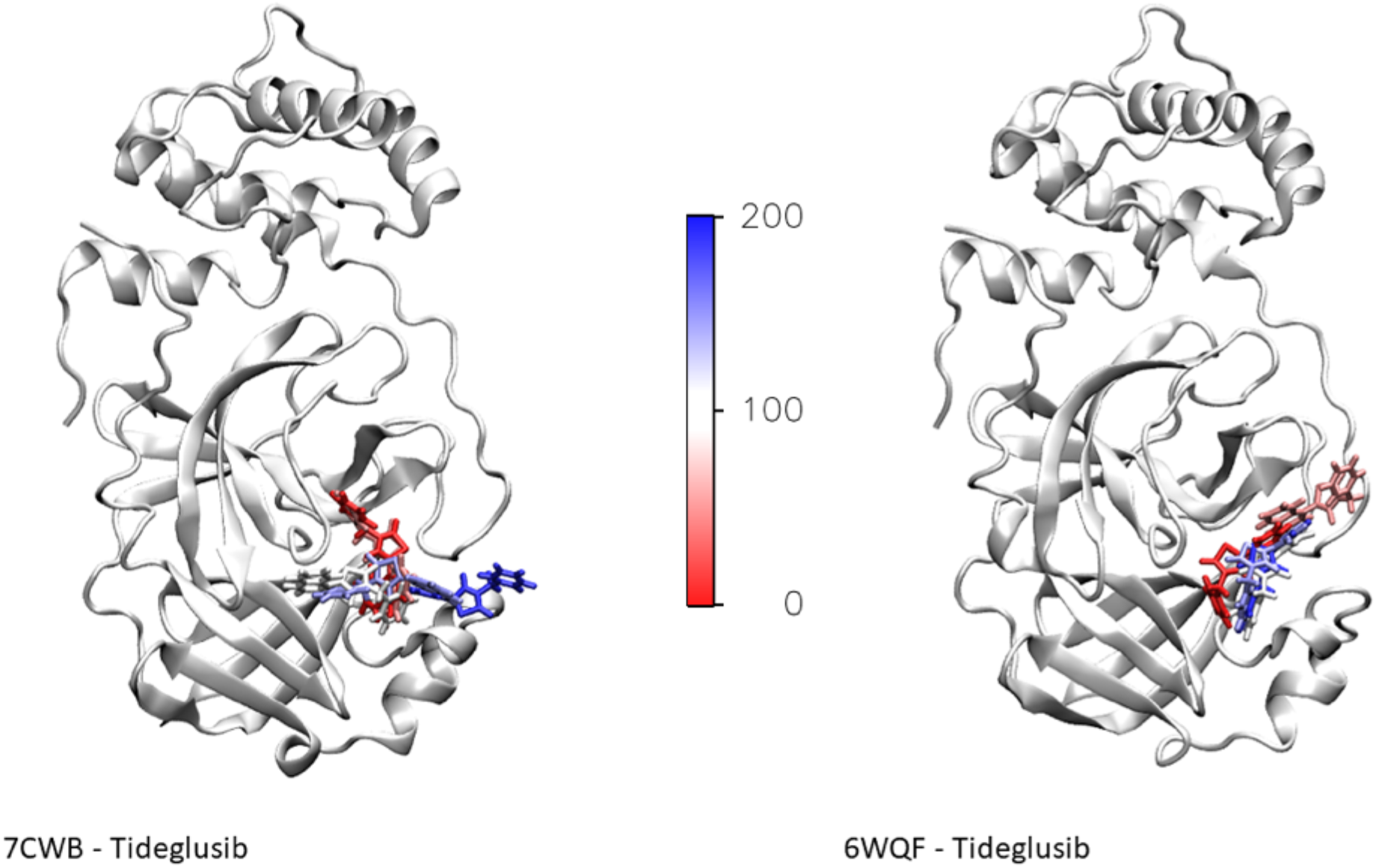
Tideglusib conformations explored during MD simulations on 7CWB (left) and 6WQF (right). Starting conformations and last frame depicted in red and blue color, respectively.

**Fig. S26.**
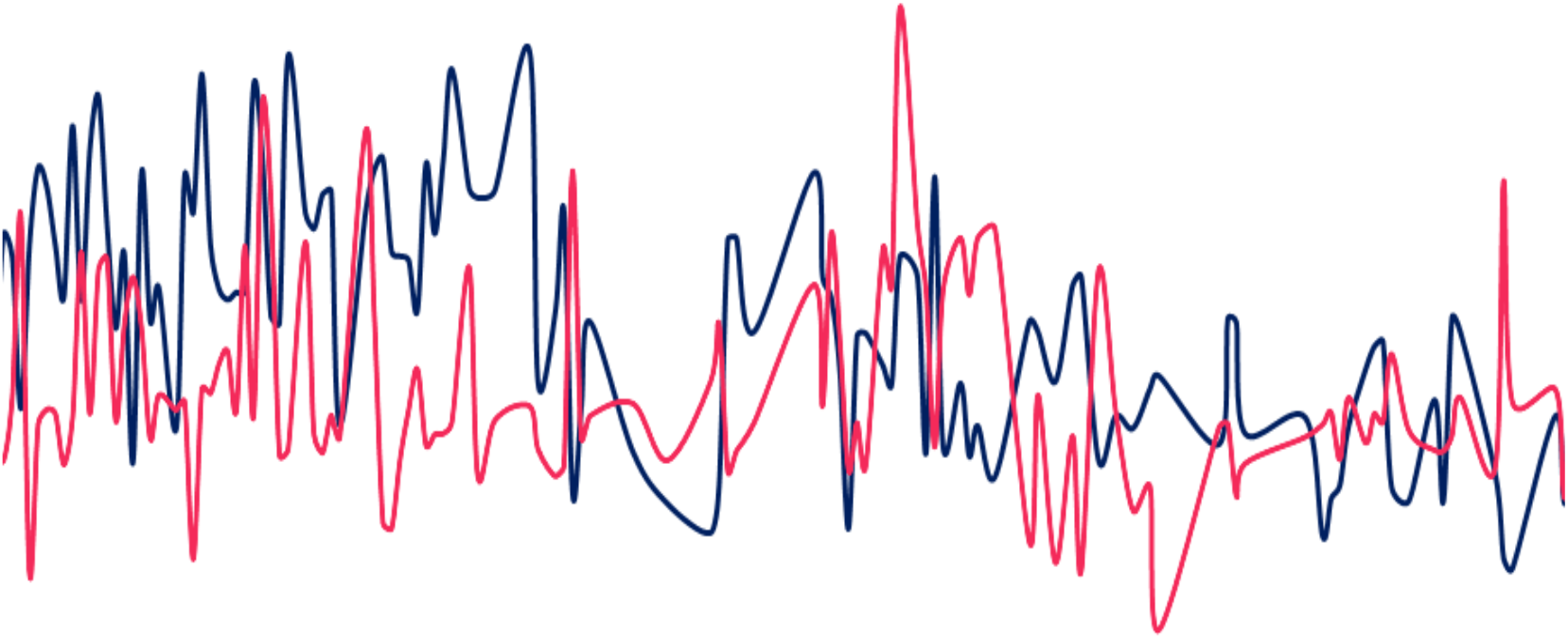
Binding pocket volume changes throughout the MD simulations.

**Fig. S27.**
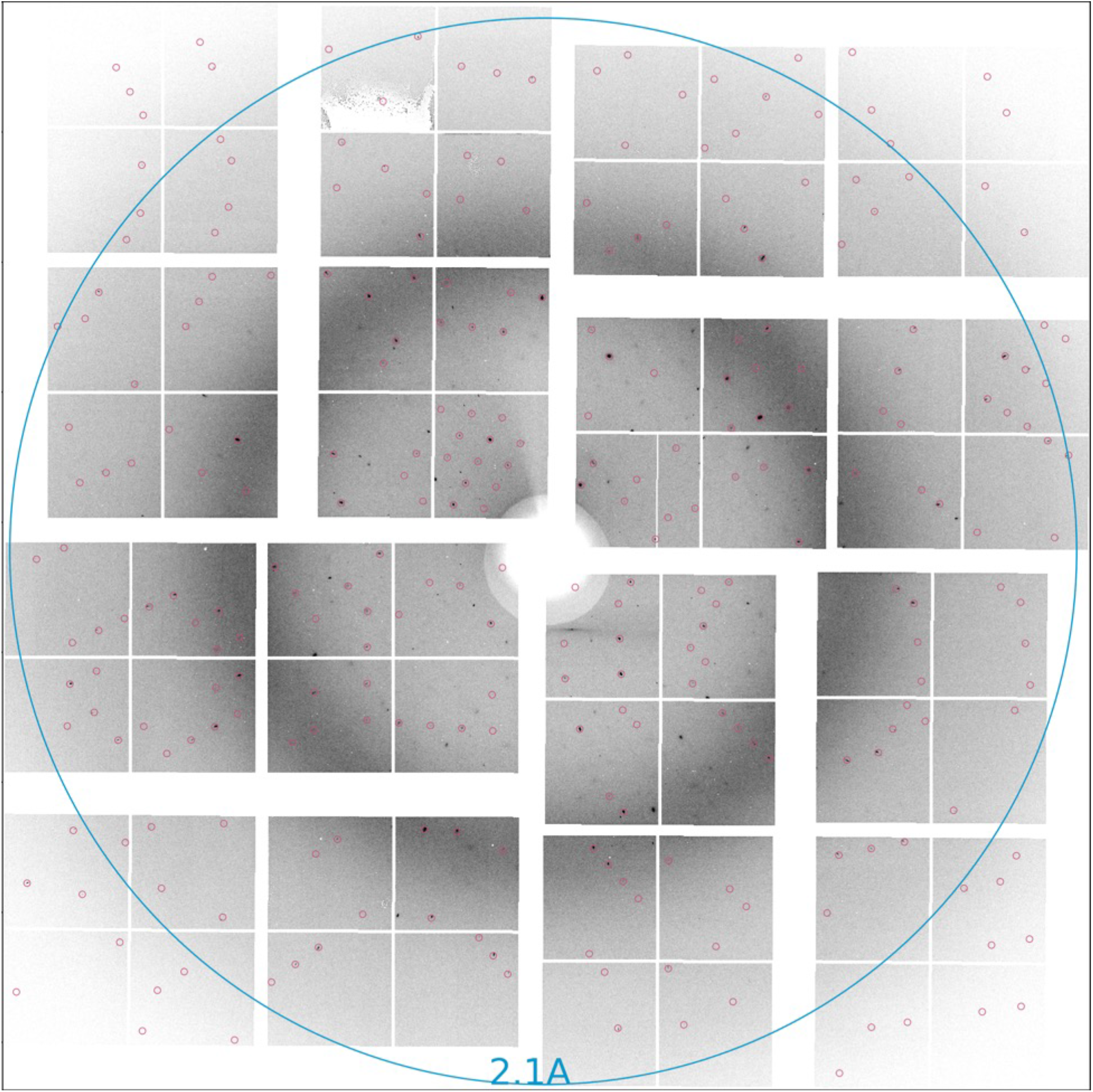
A diffraction pattern of monoclinic crystal form in space group C121 (PDB ID; 7CWB) with indexed spots.

## Supplementary Tables

**Supplementary Table S1.** Data collection and refinement statistics for X-ray crystallography

| Dataset | Monoclinic | Orthorhombic |
| --- | --- | --- |
| PDB ID | 7CWB | 7CWC |
| **Data collection<sup>1</sup>** |  |  |
| Beamline | LCLS (MFX) | LCLS (MFX) |
| Space group | C121 | P2<sub>1</sub>2<sub>1</sub>2<sub>1</sub> |
| Cell dimensions |  |  |
| *a*, *b*, *c* (Å) | 114.0, 53.5, 45.0 | 69.2, 104.3, 105.6 |
| $\alpha$, $\beta$, $\gamma$ (°) | 90, 102, 90 | 90, 90, 90 |
| Resolution (Å)<sup>2</sup> | 55.0-1.9<br>(1.98-1.90)<sup>2</sup> | 75.0-2.1<br>(2.14-2.10)<sup>2</sup> |
| *R*<sub>split</sub> | 0.63 (0.94) | 0.59 (3.33) |
| *CC*<sub>1/2</sub> | 0.996 (0.598) | 0.997 (0.634) |
| *I* / σ*I* | 10.4 (0.57) | 11.2 (0.33) |
| *CC* * | 0.999 (0.865) | 0.999 (0.678) |
| Completeness (%) | 100.0 (100.0) | 100.0 (100.0) |
| Redundancy | 825 (266) | 3105 (2025) |
| **Refinement** |  |  |
| Resolution (Å) | 34.0-1.9<br>(1.95-1.90) | 42.0-2.1<br>(2.15-2.10) |
| No. reflections | 21029 (1359) | 45238 (3044) |
| *R*<sub>work</sub> / *R*<sub>free</sub> | 0.22/0.26 (0.43/0.49) | 0.22/0.26 (0.37/0.44) |
| No. atoms |  |  |
| Protein | 2447 | 4710 |
| Ligand/Ion/Water | 113 | 166 |
| *B*-factors |  |  |
| Protein | 42.86 | 65.66 |
| Ligand/Ion/Water | 51.0 | 73.9 |
| Coordinate errors | 0.34 | 0.34 |
| R.m.s deviations |  |  |
| Bond lengths (Å) | 0.007 | 0.003 |
| Bond angles (°) | 0.869 | 0.619 |
| Ramachandran plot |  |  |
| Favored (%) | 294 (96.7) | 580 (97.5) |
| Allowed (%) | 9 (3.0) | 14 (2.4) |
| Disallowed (%) | 1 (0.3) | 1 (0.1) |

<sup>1</sup>Datasets are collected by serial crystallography
<sup>2</sup> The highest resolution shell is shown in parentheses

**Supplementary Table S2.** SFX data collection statistics

| Sample | Start Time | Stop Time | Total Run Time | Effective Run Time | Runs | Run Number |
| --- | --- | --- | --- | --- | --- | --- |
| 7CWB | 10:17:31 | 13:16:48 | 2h 59m 17s | 2h 47m 7s | 118-141 | 34 |
| 7CWC | 10:49:37 | 12:40:36 | 1h 50m 59s | 1h 36m 20s | 320-339 | 20 |

## References

A. Chauhan, S. Kalra, Identification of potent COVID-19 main protease (MPRO) inhibitors from flavonoids. Preprints (2020), doi:10.21203/rs.3.rs-34497/v1.

A. J. Tyrrell, M. L. Bynoe, Cultivation of a Novel Type of Common-cold Virus in Organ Cultures. Br. Med. J. (1965), doi:10.1136/bmj.1.5448.1467.

A. Khailany, M. Safdar, M. Ozaslan, Genomic characterization of a novel SARS-CoV-2. Gene Reports (2020), doi:10.1016/j.genrep.2020.100682.

A. M. Q. King, E. Lefkowitz, M. J. Adams, E. B. Carstens, “Order - Nidovirales” in Virus Taxonomy (Elsevier, 2012), pp. 784–794.

A. Mohd, J. A. Al-Tawfiq, Z. A. Memish, Middle East Respiratory Syndrome Coronavirus (MERS-CoV) origin and animal reservoir Susanna Lau. Virol. J. (2016), doi:10.1186/s12985-016-0544-0.

B. R. Beck, B. Shin, Y. Choi, S. Park, K. Kang, Predicting commercially available antiviral drugs that may act on the novel coronavirus (SARS-CoV-2) through a drug-target interaction deep learning model. Comput. Struct. Biotechnol. J. (2020), doi:10.1016/j.csbj.2020.03.025.

C. A. Menendez, F. Bylehn, G.R. Perez-Lemuz, W. Alvarado, J. J de Pablo, Molecular Characterization of Ebselen Binding Activity to SARS-CoV-2 Main Protease. Preprints (2020).

C. Ma, M. D. Sacco, B. Hurst, J. A. Townsend, Y. Hu, T. Szeto, X. Zhang, B. Tarbet, M. T. Marty, Y. Chen, J. Wang, Boceprevir, GC-376, and calpain inhibitors II, XII inhibit SARS-CoV-2 viral replication by targeting the viral main protease. Cell Res. (2020), doi:10.1038/s41422-020-0356-z.

C.R. Astell, R.A. Holt, S.J.M. Jones, M.A Marra, “Genome organization and structural aspects of the SARS-related virus” in Coronaviruses with Special Emphasis on First Insights Concerning SARS, A. Schmidt, O. Weber, M.H. Wolff, Eds. Birkhäuser Advances in Infectious Diseases BAID(2005), (Birkhäuser, Basel, 2005), pp 101–128, doi.org/10.1007/3-7643-7339-3_5.

C. Zhang, Z. Wu, J. W. Li, H. Zhao, G. Q. Wang, Cytokine release syndrome in severe COVID-19: interleukin-6 receptor antagonist tocilizumab may be the key to reduce mortality. Int. J. Antimicrob. Agents (2020), doi:10.1016/j.ijantimicag.2020.105954.

D. C. Dinesh, S. Tamilarasan, K. Rajaram, E. Bouřa, Antiviral Drug Targets of Single-Stranded RNA Viruses Causing Chronic Human Diseases. Curr. Drug Targets (2019), doi:10.2174/1389450119666190920153247.

D. Cavalla, Predictive methods in drug repurposing: Gold mine or just a bigger haystack? Drug Discov. Today (2013), doi:10.1016/j.drudis.2012.12.009.

D. E. Gordon, G. M. Jang, M. Bouhaddou, et al. A SARS-CoV-2 protein interaction map reveals targets for drug repurposing. Nature. (2020), doi:10.1038/s41586-020-2286-9.

D. Elmezayen, A. Al-Obaidi, A. T. Şahin, K. Yelekçi, Drug repurposing for coronavirus (COVID-19): in silico screening of known drugs against coronavirus 3CL hydrolase and protease enzymes. J. Biomol. Struct. Dyn. (2020), doi:10.1080/07391102.2020.1758791.

D. G. Ahn, H. J. Shin, M. H. Kim, S. Lee, H. S. Kim, J. Myoung, B. T. Kim, S. J. Kim, Current status of epidemiology, diagnosis, therapeutics, and vaccines for novel coronavirus disease 2019 (COVID-19). J. Microbiol. Biotechnol. (2020), doi:10.4014/jmb.2003.03011.

D. Kim, J. Y. Lee, J. S. Yang, J. W. Kim, V. N. Kim, H. Chang, The Architecture of SARS-CoV-2 Transcriptome. Cell (2020), doi:10.1016/j.cell.2020.04.011.

D. L. Heymann, G. Rodier, Global Surveillance, National Surveillance, and SARS. Emerg. Infect. Dis. (2004), doi:10.3201/eid1002.031038.

D. McGonagle, J. S. O’Donnell, K. Sharif, P. Emery, C. Bridgewood, Immune mechanisms of pulmonary intravascular coagulopathy in COVID-19 pneumonia. Lancet Rheumatol. (2020), doi:10.1016/S2665-9913(20)30121-1.

D. McGonagle, K. Sharif, A. O’Regan, C. Bridgewood, Interleukin-6 use in COVID-19 pneumonia related macrophage activation syndrome. Autoimmun. Rev. (2020), doi:10.1016/j.autrev.2020.102537.

D. Ragab, H. Salah Eldin, M. Taeimah, R. Khattab, R. Salem, The COVID-19 Cytokine Storm; What We Know So Far. Front. Immunol. (2020), doi:10.3389/fimmu.2020.01446.

D. Rathnayake, J. Zheng, Y. Kim, K. D. Perera, S. Mackin, D. K. Meyerholz, M. M. Kashipathy, K. P. Battaile, S. Lovell, S. Perlman, W. C. Groutas, K.-O. Chang, 3C-like protease inhibitors block coronavirus replication in vitro and improve survival in MERS-CoV–infected mice. Sci. Transl. Med. (2020), doi:10.1126/scitranslmed.abc5332.

D. Robertson, US FDA approves new class of HIV therapeutics. Nat. Biotechnol. (2003), doi:10.1038/nbt0503-470.

D. Suárez, N. Díaz, SARS-CoV 2 Main Protease: A Molecular Dynamics Study, J. Chem. Inf. Model. (2020), doi:10.1021/acs.jcim.0c00575.

D. W. Kneller, G. Phillips, H. M. O’Neill, R. Jedrzejczak, L. Stols, P. Langan, A. Joachimiak, L. Coates, A. Kovalevsky, Structural plasticity of SARS-CoV-2 3CL Mpro active site cavity revealed by room temperature X-ray crystallography. Nat. Commun. (2020), doi:10.1038/s41467-020-16954-7.

di Mauro Gabriella, S. Cristina, R. Concetta, R. Francesco, C. Annalisa, SARS-Cov-2 infection: Response of human immune system and possible implications for the rapid test and treatment. Int. Immunopharmacol. (2020), doi:10.1016/j.intimp.2020.106519.

E. Gorbalenya, E. J. Snijder, Viral cysteine proteinases. Perspect. Drug Discov. Des. (1996), doi:10.1007/BF02174046.

E. J. T. Aguila, I. H. Y. Cua, Repurposed GI Drugs in the Treatment of COVID-19. Dig. Dis. Sci. (2020), doi:10.1007/s10620-020-06430-z.

E. Petersen, M. Koopmans, U. Go, D. H. Hamer, N. Petrosillo, F. Castelli, M. Storgaard, S. Al Khalili, L. Simonsen, Comparing SARS-CoV-2 with SARS-CoV and influenza pandemics. Lancet Infect. Dis. (2020), doi:10.1016/S1473-3099(20)30484-9.

E. Pihan, L. Colliandre, J. F. Guichou, D. Douguet, E-Drug3D: 3D structure collections dedicated to drug repurposing and fragment-based drug design. Bioinformatics (2012), doi: 10.1093/bioinformatics/bts186.

E. W. Su, T. M. Sanger: Systematic drug repositioning through mining adverse event data in ClinicalTrials. gov. Peer J (2017), doi: 10.7717/peerj.3154

E. Węglarz-Tomczak, J. M. Tomczak, M. Talma, S. Brul, Ebselen as a highly active inhibitor of PLProCoV2. bioRxiv (2020), doi:10.1101/2020.05.17.100768.

F. Braun, M. Lütgehetmann, S. Pfefferle, M. N. Wong, A. Carsten, M. T. Lindenmeyer, D. Nörz, F. Heinrich, K. Meißner, D. Wichmann, S. Kluge, O. Gross, K. Pueschel, A. S. Schröder, C. Edler, M. Aepfelbacher, V. G. Puelles, T. B. Huber, SARS-CoV-2 renal tropism associates with acute kidney injury. Lancet (2020), doi:10.1016/s0140-6736(20)31759-1.

F. Hikmet, L. Méar, Å. Edvinsson, P. Micke, M. Uhlén, C. Lindskog, The protein expression profile of ACE2 in human tissues. Mol. Syst. Biol. (2020), doi:10.15252/msb.20209610.

F. Huang, C. Zhang, Q. Liu, Y. Zhao, Y. Zhang, Y. Qin, X. Li, C. Li, C. Zhou, N. Jin, C. Jiang, Identification of amitriptyline HCl, flavin adenine dinucleotide, azacitidine and calcitriol as repurposing drugs for influenza A H5N1 virus-induced lung injury. PLoS Pathog. (2020), doi:10.1371/journal.ppat.1008341.

F. Wu, S. Zhao, B. Yu, Y. M. Chen, W. Wang, Z. G. Song, Y. Hu, Z. W. Tao, J. H. Tian, Y. Y. Pei, M. L. Yuan, Y. L. Zhang, F. H. Dai, Y. Liu, Q. M. Wang, J. J. Zheng, L. Xu, E. C. Holmes, Y. Z. Zhang, A new coronavirus associated with human respiratory disease in China. Nature (2020), doi:10.1038/s41586-020-2008-3.

G. Andersen, A. Rambaut, W. I. Lipkin, E. C. Holmes, R. F. Garry, The proximal origin of SARS-CoV-2. Nat. Med. (2020), doi:10.1038/s41591-020-0820-9.

G. Blaj, A. Dragone, C. J. Kenney, F. Abu-Nimeh, P. Caragiulo, D. Doering, M. Kwiatkowski, B. Markovic, J. Pines, M. Weaver, S. Boutet, G. Carini, C. E. Chang, P. Hart, J. Hasi, M. Hayes, R. Herbst, J. Koglin, K. Nakahara, J. Segal, G. Haller, Performance of ePix10K, a high dynamic range, gain auto-ranging pixel detector for FELs, in AIP Conference Proceedings (2019), doi:10.1063/1.5084693.

G. G. Chang. “Quaternary Structure of the SARS Coronavirus Main Protease.” In Molecular Biology of the SARS-Coronavirus S.K Lal Eds. (Springer, Berlin, Heidelberg, 2010), pp 115–128 (2010), doi:10.1007/978-3-642-03683-5_8.

G. Sierra, A. Batyuk, Z. Sun, A. Aquila, M. S. Hunter, T. J. Lane, M. Liang, C. H. Yoon, R. Alonso-Mori, R. Armenta, J. C. Castagna, M. Hollenbeck, T. O. Osier, M. Hayes, J. Aldrich, R. Curtis, J. E. Koglin, T. Rendahl, E. Rodriguez, S. Carbajo, S. Guillet, R. Paul, P. Hart, K. Nakahara, G. Carini, H. Demirci, E. H. Dao, B. M. Hayes, Y. P. Rao, M. Chollet, Y. Feng, F. D. Fuller, C. Kupitz, T. Sato, M. H. Seaberg, S. Song, T. B. Van Driel, H. Yavas, D. Zhu, A. E. Cohen, S. Wakatsuki, S. Boutet, The macromolecular femtosecond crystallography instrument at the linac coherent light source. J. Synchrotron Radiat. (2019), doi:10.1107/S1600577519001577.

H. Chen, F. Cheng, J. Li, IDrug: Integration of drug repositioning and drug-target prediction via cross-network embedding. PLoS Comput. Biol. (2020), doi:10.1371/journal.pcbi.1008040.

H. M. Mengist, X. Fan, T. Jin, Designing of improved drugs for COVID-19: Crystal structure of SARS-CoV-2 main protease Mpro. Signal Transduct. Target. Ther. (2020), doi:10.1038/s41392-020-0178-y.

H. Sies, M. J. Parnham, Potential therapeutic use of ebselen for COVID-19 and other respiratory viral infections. Free Radic. Biol. Med. (2020), doi:10.1016/j.freeradbiomed.2020.06.032.

H. Wang, S. He, W. Deng, Y. Zhang, G. Li, J. Sun, W. Zhao, Y. Guo, Z. Yin, D. Li, L. Shang, Comprehensive Insights into the Catalytic Mechanism of Middle East Respiratory Syndrome 3C-Like Protease and Severe Acute Respiratory Syndrome 3C-Like Protease. ACS Catal. (2020), doi:10.1021/acscatal.0c00110.

I. Bahar, T. R. Lezon, A. Bakan and I. H. Shrivastava, Normal mode analysis of biomolecular structures: functional mechanisms of membrane proteins, Chem. Rev. (2010), doi:10.1021/cr900095e.

J. O’Shea, A. Ma, P. Lipsky, Cytokines and autoimmunity. Nat. Rev. Immunol. (2002), doi:10.1038/nri702.

J. M. Bumb, F. Enning and F. M. Leweke, Drug repurposing and emerging adjunctive treatments for schizophrenia, Expert Opinion on Pharmacotherapy (2020), doi:10.1517/14656566.2015.1032248

J. S. Kahn, K. McIntosh, History and Recent Advances in Coronavirus Discovery. Pediatr. Infect. Dis. J. (2005), doi:10.1097/01.inf.0000188166.17324.60.

J. Wang, Fast Identification of Possible Drug Treatment of Coronavirus Disease-19 (COVID-19) through Computational Drug Repurposing Study. J. Chem. Inf. Model. (2020), doi:10.1021/acs.jcim.0c00179.

J. Ziebuhr, E. J. Snijder, A. E. Gorbalenya, Virus-encoded proteinases and proteolytic processing in the Nidovirales. J. Gen. Virol. (2000), doi:10.1099/0022-1317-81-4-853.

K. Hu, W. jie Guan, Y. Bi, W. Zhang, L. Li, B. Zhang, Q. Liu, Y. Song, X. Li, Z. Duan, Q. Zheng, Z. Yang, J. Liang, M. Han, L. Ruan, C. Wu, Y. Zhang, Z. hua Jia, N. shan Zhong, Efficacy and safety of Lianhuaqingwen capsules, a repurposed Chinese herb, in patients with coronavirus disease 2019: A multicenter, prospective, randomized controlled trial. Phytomedicine (2020), doi:10.1016/j.phymed.2020.153242.

K. Ratia, A. Kilianski, Y. M. Baez-Santos, S. C. Baker, A. Mesecar, Structural Basis for the Ubiquitin-Linkage Specificity and deISGylating Activity of SARS-CoV Papain-Like Protease. PLoS Pathog. (2014), doi:10.1371/journal.ppat.1004113.

L. Zhang, D. Lin, X. Sun, U. Curth, C. Drosten, L. Sauerhering, S. Becker, K. Rox, R. Hilgenfeld, Crystal structure of SARS-CoV-2 main protease provides a basis for design of improved a-ketoamide inhibitors. Science (80-.). (2020), doi:10.1126/science.abb3405.

M. Madjid, P. Safavi-Naeini, S. D. Solomon, O. Vardeny, Potential Effects of Coronaviruses on the Cardiovascular System: A Review. JAMA Cardiol. (2020), doi:10.1001/jamacardio.2020.1286.

M. M. Ghahremanpour, J. Tirado-Rives, M. Deshmukh, J. A. Ippolito, C. Zhang, I. Cabeza de Vaca, M. Liosi, K. S. Anderson, W. L. Jorgensen. Identification of 14 Known Drugs as Inhibitors of the Main Protease of SARS-CoV-2. bioRxiv (2020), doi:10.1101/2020.08.28.271957.

M. Martin-Garcia, C. E. Conrad, J. Coe, S. Roy-Chowdhury, P. Fromme, Serial femtosecond crystallography: A revolution in structural biology. Arch. Biochem. Biophys. (2016), doi:10.1016/j.abb.2016.03.036.

M. Zaki, S. Van Boheemen, T. M. Bestebroer, A. D. M. E. Osterhaus, R. A. M. Fouchier, Isolation of a novel coronavirus from a man with pneumonia in Saudi Arabia. N. Engl. J. Med. (2012), doi:10.1056/NEJMoa1211721.

M. Zmudzinski, W. Rut, K. Olech, J. Granda, M. Giurg, M. Burda-Grabowska, L. Zhang, X. Sun, Z. Lv, D. Nayak, M. Kesik-Brodacka, S. K. Olsen, R. Hilgenfeld, M. Drag, Ebselen derivatives are very potent dual inhibitors of SARS-CoV-2 proteases - PLpro and Mpro in vitro studies. bioRxiv (2020), doi.org/10.1101/2020.08.30.273979.

N. Muralidharan, R. Sakthivel, D. Velmurugan, M. M. Gromiha, Computational studies of drug repurposing and synergism of lopinavir, oseltamivir and ritonavir binding with SARS-CoV-2 protease against COVID-19. J. Biomol. Struct. Dyn. (2020), doi:10.1080/07391102.2020.1752802.

N. Zhong, S. Zhang, P. Zou, J. Chen, X. Kang, Z. Li, C. Liang, C. Jin, B. Xia, Without Its N-Finger, the Main Protease of Severe Acute Respiratory Syndrome Coronavirus Can Form a Novel Dimer through Its C-Terminal Domain. J. Virol. (2008), doi:10.1128/j

O. S. Choo, D. Yoon, Y. Choi, S. Jo, H. M. Jung, J. Y. An, Y. H. Choung, Drugs for hyperlipidaemia may slow down the progression of hearing loss in the elderly: A drug repurposing study. Basic Clin. Pharmacol. Toxicol. (2019), doi:10.1111/bcpt.13150.

P. Anand, A. Puranik, M. Aravamudan, A. J. Venkatakrishnan, V. Soundararajan, SARS-CoV-2 strategically mimics proteolytic activation of human ENaC. Elife (2020), doi:10.7554/eLife.58603.

R. J. Khan, R. K. Jha, G. M. Amera, M. Jain, E. Singh, A. Pathak, R. P. Singh, J. Muthukumaran, A. K. Singh, Targeting SARS-CoV-2: a systematic drug repurposing approach to identify promising inhibitors against 3C-like proteinase and 2’-O-ribose methyltransferase. J. Biomol. Struct. Dyn. (2020), doi:10.1080/07391102.2020.1753577.

R. Root-Bernstein, Human immunodeficiency virus proteins mimic human T cell receptors inducing cross-reactive antibodies. Int. J. Mol. Sci. (2017), doi:10.3390/ijms18102091.

S. Abdul-Rasool, B. C. Fielding, Understanding Human Coronavirus HCoV-NL63. Open Virol. J. (2010).

S. Adem, V. Eyupoglu, I. Sarfraz, A. Rasul, M. Ali Identification of Potent COVID-19 Main Protease (Mpro) Inhibitors from Natural Polyphenols: An in Silico Strategy Unveils a Hope against CORONA. Preprints (2020), doi: 10.20944/preprints202003.0333.v1.

S. Amamuddy, G. M. Verkhivker and Ö. T. Bishop, Impact of emerging mutations on the dynamic properties the SARS-CoV-2 main protease: an in silico investigation, Preprints (2020), https://doi.org/10.1101/2020.05.29.123190

S. Belouzard, J. K. Millet, B. N. Licitra, G. R. Whittaker, Mechanisms of coronavirus cell entry mediated by the viral spike protein. Viruses (2012), doi:10.3390/v4061011.

S. Choudhary, Y. S. Malik, S. Tomar, Identification of SARS-CoV-2 Cell Entry Inhibitors by Drug Repurposing Using in silico Structure-Based Virtual Screening Approach. Front. Immunol. (2020), doi: 10.3389/fimmu.2020.01664.

S. Durdagi, B. Aksoydan, B. Dogan, K. Sahin, A. Shahraki, Screening of Clinically Approved and Investigation Drugs as Potential Inhibitors of COVID-19 Main Protease: A Virtual Drug Repurposing Study. ChemRxiv (2020), doi:10.26434/chemrxiv.12032712.v1.

S. Dutta, P. Sengupta, SARS-CoV-2 and Male Infertility: Possible Multifaceted Pathology. Reprod. Sci. (2020), doi:10.1007/s43032-020-00261-z.

S. G. Viveiros Rosa, W. C. Santos, Clinical trials on drug repositioning for COVID-19 treatment. Rev. Panam. Salud Publica/Pan Am. J. Public Heal. (2020), doi:10.26633/RPSP.2020.40.

S. Isabel, L. Graña-Miraglia, J. M. Gutierrez, C. Bundalovic-Torma, H. E. Groves, M. R. Isabel, A. R. Eshaghi, S. N. Patel, J. B. Gubbay, T. Poutanen, D. S. Guttman, S. M. Poutanen, Evolutionary and structural analyses of SARS-CoV-2 D614G spike protein mutation now documented worldwide. Sci. Rep. (2020), doi:10.1038/s41598-020-70827-z.

S. K. P. Lau, P. C. Y. Woo, C. C. Y. Yip, H. Tse, H. W. Tsoi, V. C. C. Cheng, P. Lee, B. S. F. Tang, C. H. Y. Cheung, R. A. Lee, L. Y. So, Y. L. Lau, K. H. Chan, K. Y. Yuen, Coronavirus HKU1 and other coronavirus infections in Hong Kong. J. Clin. Microbiol. (2006), doi:10.1128/JCM.02614-05.

S. Khaerunnisa, H. Kurniawan, R. Awaluddin, S. Suhartati, Potential Inhibitor of COVID-19 Main Protease (M pro) from Several Medicinal Plant Compounds by Molecular Docking Study. Preprints (2020), doi:10.20944/preprints202003.0226.v1.

S. L. Maude, D. Barrett, D. T. Teachey, S. A. Grupp, Managing cytokine release syndrome associated with novel T cell-engaging therapies. Cancer J. (United States) (2014), doi:10.1097/PPO.0000000000000035.

S. Laha, J. Chakraborty, S. Das, S. K. Manna, S. Biswas, R. Chatterjee, Characterizations of SARS-CoV-2 mutational profile, spike protein stability and viral transmission. Infect. Genet. Evol. (2020), doi:10.1016/j.meegid.2020.104445.

S. M. Strittmatter, Overcoming drug development bottlenecks with repurposing: Old drugs learn new tricks. Nat. Med. (2014), doi:10.1038/nm.3595.

S. Pushpakom, F. Iorio, P. A. Eyers, K. J. Escott, S. Hopper, A. Wells, A. Doig, T. Guilliams, J. Latimer, C. McNamee, A. Norris, P. Sanseau, D. Cavalla, M. Pirmohamed, Drug repurposing: Progress, challenges and recommendations. Nat. Rev. Drug Discov. (2018), doi:10.1038/nrd.2018.168.

S. Ullrich, C. Nitsche, The SARS-CoV-2 main protease as drug target. Bioorganic Med. Chem. Lett. (2020), doi:10.1016/j.bmcl.2020.127377.

T. B. Van Driel, S. Nelson, R. Armenta, G. Blaj, S. Boo, S. Boutet, D. Doering, A. Dragone, P. Hart, G. Haller, C. Kenney, M. Kwaitowski, L. Manger, M. McKelvey, K. Nakahara, M. Oriunno, T. Sato, M. Weaver, The ePix10k 2-megapixel hard X-ray detector at LCLS. J. Synchrotron Radiat. (2020), doi:10.1107/S1600577520004257.

T. Joshi, T. Joshi, P. Sharma, S. Mathpal, H. Pundir, V. Bhatt, S. Chandra, In silico screening of natural compounds against COVID-19 by targeting Mpro and ACE2 using molecular docking. Eur. Rev. Med. Pharmacol. Sci. (2020), doi:10.26355/eurrev_202004_21036.

T. N. Jarada, J. G. Rokne, R. Alhajj, A review of computational drug repositioning: Strategies, approaches, opportunities, challenges, and directions. J. Cheminform. (2020), doi:10.1186/s13321-020-00450-7.

T. Pillaiyar, M. Manickam, V. Namasivayam, Y. Hayashi, S. H. Jung, An overview of severe acute respiratory syndrome-coronavirus (SARS-CoV) 3CL protease inhibitors: Peptidomimetics and small molecule chemotherapy. J. Med. Chem. (2016), doi:10.1021/acs.jmedchem.5b01461.

T. T. Ashburn, K. B. Thor, Drug repositioning: Identifying and developing new uses for existing drugs. Nat. Rev. Drug Discov. (2004), doi:10.1038/nrd1468.

T. Ton, F. Gentile, M. Hsing, F. Ban, A. Cherkasov, Rapid Identification of Potential Inhibitors of SARS-CoV-2 Main Protease by Deep Docking of 1.3 Billion Compounds. Mol. Inform. (2020), doi:10.1002/minf.202000028.

V. Thiel, K. A. Ivanov, Á. Putics, T. Hertzig, B. Schelle, S. Bayer, B. Weißbrich, E. J. Snijder, H. Rabenau, H. W. Doerr, A. E. Gorbalenya, J. Ziebuhr, Mechanisms and enzymes involved in SARS coronavirus genome expression. J. Gen. Virol. (2003), doi:10.1099/vir.0.19424-0.

W. Dai, B. Zhang, X. M. Jiang, H. Su, J. Li, Y. Zhao, X. Xie, Z. Jin, J. Peng, F. Liu, C. Li, Y. Li, F. Bai, H. Wang, X. Cheng, X. Cen, S. Hu, X. Yang, J. Wang, X. Liu, G. Xiao, H. Jiang, Z. Rao, L. K. Zhang, Y. Xu, H. Yang, H. Liu, Structure-based design of antiviral drug candidates targeting the SARS-CoV-2 main protease. Science (80-.). (2020), doi:10.1126/science.abb4489.

W. Zhang, Y. Zhao, F. Zhang, Q. Wang, T. Li, Z. Liu, J. Wang, Y. Qin, X. Zhang, X. Yan, X. Zeng, S. Zhang, The use of anti-inflammatory drugs in the treatment of people with severe coronavirus disease 2019 (COVID-19): The experience of clinical immunologists from China. Clin. Immunol. (2020), doi:10.1016/j.clim.2020.108393.

World Health Organization, (WHO), “Timeline: WHO’s COVID-19 response” (Publication WHO, 2020; www.who.int/emergencies/diseases/novel-coronavirus-2019/interactive-timeline [the easiest access to this source is via the URL].

Y. Chen, Q. Liu, D. Guo, Emerging coronaviruses: Genome structure, replication, and pathogenesis. J. Med. Virol. (2020), doi:10.1002/jmv.25681.

Y. Jia, G. Shen, Y. Zhang, K.-S. Huang, H.-Y. Ho, W.-S. Hor, C.-H. Yang, C. Li, W.-L. Wang, Analysis of the mutation dynamics of SARS-CoV-2 reveals the spread history and emergence of RBD mutant with lower ACE2 binding affinity. bioRxiv (2020), doi:10.1101/2020.04.09.034942.

Y. Kumar, H. Singh, C. N. Patel, In silico prediction of potential inhibitors for the Main protease of SARS-CoV-2 using molecular docking and dynamics simulation based drug-repurposing. J. Infect. Public Health (2020), doi:10.1016/j.jiph.2020.06.016.

Y. W. Chen, C.-P. B. Yiu, K.-Y. Wong, Prediction of the SARS-CoV-2 (2019-nCoV) 3C-like protease (3CLpro) structure: virtual screening reveals velpatasvir, ledipasvir, and other drug repurposing candidates. F1000Research (2020), doi:10.12688/f1000research.22457.1.

Y. Zhou, Y. Hou, J. Shen, Y. Huang, W. Martin, F. Cheng, Network-based drug repurposing for novel coronavirus 2019-nCoV/SARS-CoV-2. Cell Discov. (2020), doi:10.1038/s41421-020-0153-3.

Z. Jin, X. Du, Y. Xu, Y. Deng, M. Liu, Y. Zhao, B. Zhang, X. Li, L. Zhang, C. Peng, Y. Duan, J. Yu, L. Wang, K. Yang, F. Liu, R. Jiang, X. Yang, T. You, X. Liu, X. Yang, F. Bai, H. Liu, X. Liu, L. W. Guddat, W. Xu, G. Xiao, C. Qin, Z. Shi, H. Jiang, Z. Rao, H. Yang, Structure of Mpro from SARS-CoV-2 and discovery of its inhibitors. Nature (2020), doi:10.1038/s41586-020-2223-y.

## Supplementary References

A. Barty, R. A. Kirian, F. R. N. C. Maia, M. Hantke, C. H. Yoon, T. A. White, H. Chapman, Cheetah: Software for high-throughput reduction and analysis of serial femtosecond X-ray diffraction data. J. Appl. Crystallogr. (2014), doi:10.1107/S1600576714007626.

A. J. McCoy, R. W. Grosse-Kunstleve, P. D. Adams, M. D. Winn, L. C. Storoni, R. J. Read, Phaser crystallographic software. J. Appl. Crystallogr. (2007), doi:10.1107/S0021889807021206.

D. C. Bas, D. M. Rogers, J. H. Jensen, Very fast prediction and rationalization of pKa values for protein-ligand complexes. Proteins Struct. Funct. Genet. (2008), doi:10.1002/prot.22102.

D. Damiani, M. Dubrovin, I. Gaponenko, W. Kroeger, T. J. Lane, A. Mitra, C. P. O’Grady, A. Salnikov, A. Sanchez-Gonzalez, D. Schneider, C. H. Yoon, Linac Coherent Light Source data analysis using psana. J. Appl. Crystallogr. (2016), doi:10.1107/S1600576716004349.

E. Harder, W. Damm, J. Maple, C. Wu, M. Reboul, J. Y. Xiang, L. Wang, D. Lupyan, M. K. Dahlgren, J. L. Knight, J. W. Kaus, D. S. Cerutti, G. Krilov, W. L. Jorgensen, R. Abel, R. A. Friesner, OPLS3: A Force Field Providing Broad Coverage of Drug-like Small Molecules and Proteins. J. Chem. Theory Comput. (2016), doi:10.1021/acs.jctc.5b00864.

G. J. Martyna, D. J. Tobias, Klein, Constant pressure molecular dynamics algorithms. J. Chem. Phys. (1994), doi: 10.1063/1.467468.

G. Madhavi Sastry, M. Adzhigirey, T. Day, R. Annabhimoju, W. Sherman, Protein and ligand preparation: Parameters, protocols, and influence on virtual screening enrichments. J. Comput. Aided. Mol. Des. (2013), doi:10.1007/s10822-013-9644-8.

H. J. C. Berendsen, J. R. Grigera, T. P. Straatsma, The missing term in effective pair potentials. Journal of Physical Chemistry (1987).

H. R. Powell, O. Johnson, A. G. W. Leslie, Autoindexing diffraction images with iMosflm. Acta Crystallogr. Sect. D Biol. Crystallogr. (2013), doi:10.1107/S0907444912048524.

J. A. McCammon, S. C. Harvey, Dynamics of Proteins and Nucleic Acids. (Cambridge: Cambridge University Press.,1987), doi:10.1017/CBO9781139167864.

J. C. Shelley, A. Cholleti, L. L. Frye, J. R. Greenwood, M. R. Timlin, M. Uchimaya, Epik: A software program for pKa prediction and protonation state generation for drug-like molecules. J. Comput. Aided. Mol. Des. (2007), doi:10.1007/s10822-007-9133-z.

J. Grant, A. P. C. Rodrigues, K. M. ElSawy, J. A. McCammon, L. S. D. Caves, Bio3d: An R package for the comparative analysis of protein structures. Bioinformatics (2006), doi: 10.1093/bioinformatics/btl461.

J. M. Duisenberg, Indexing in single-crystal diffractometry with an obstinate list of reflections. J. Appl. Crystallogr. (1992), doi:10.1107/S0021889891010634.

J. Thayer, D. Damiani, C. Ford, M. Dubrovin, I. Gaponenko, C. P. O’Grady, W. Kroeger, J. Pines, T. J. Lane, A. Salnikov, D. Schneider, T. Tookey, M. Weaver, C. H. Yoon, A. Perazzo, Data systems for the Linac coherent light source. Adv. Struct Chem Imaging. (2017), doi:10.1186/s40679-016-0037-7.

K. J. Bowers, E. Chow, H. Xu, R. O. Dror, M. P. Eastwood, B. A. Gregersen, J. L. Klepeis, I. Kolossvary, M. A. Moraes, F. D. Sacerdoti, J. K. Salmon, Y. Shan, D. E. Shaw, in Proceedings of the 2006 ACM/IEEE Conference on Supercomputing, SC’06 (2006).

M. D. Winn, C. C. Ballard, K. D. Cowtan, E. J. Dodson, P. Emsley, P. R. Evans, R. M. Keegan, E. B. Krissinel, A. G. W. Leslie, A. McCoy, S. J. McNicholas, G. N. Murshudov, N. S. Pannu, E. A. Potterton, H. R. Powell, R. J. Read, A. Vagin, K. S. Wilson, Overview of the CCP4 suite and current developments. Acta Crystallogr. Sect. D Biol. Crystallogr. (2011), doi:10.1107/S0907444910045749.

M. P. Jacobson, D. L. Pincus, C. S. Rapp, T. J. F. Day, B. Honig, D. E. Shaw, R. A. Friesner, A Hierarchical Approach to All-Atom Protein Loop Prediction. Proteins Struct. Funct. Genet. (2004), doi:10.1002/prot.10613.

P. A. Karplus, K. Diederichs, Linking crystallographic model and data quality. Science (80-.). (2012), doi:10.1126/science.1218231.

P. D. Adams, P. V. Afonine, G. Bunkóczi, V. B. Chen, I. W. Davis, N. Echols, J. J. Headd, L. W. Hung, G. J. Kapral, R. W. Grosse-Kunstleve, A. J. McCoy, N. W. Moriarty, R. Oeffner, R. J. Read, D. C. Richardson, J. S. Richardson, T. C. Terwilliger, P. H. Zwart, PHENIX: A comprehensive Python-based system for macromolecular structure solution. Acta Crystallogr. Sect. D Biol. Crystallogr. (2010), doi:10.1107/S0907444909052925.

P. Emsley, K. Cowtan, Coot: Model-building tools for molecular graphics. Acta Crystallogr. Sect. D Biol. Crystallogr. (2004), doi:10.1107/S0907444904019158.

R. A. Friesner, J. L. Banks, R. B. Murphy, T. A. Halgren, J. J. Klicic, D. T. Mainz, M. P. Repasky, E. H. Knoll, M. Shelley, J. K. Perry, D. E. Shaw, P. Francis, P. S. Shenkin, Glide: A New Approach for Rapid, Accurate Docking and Scoring. 1. Method and Assessment of Docking Accuracy. J. Med. Chem. (2004), doi:10.1021/jm0306430.

R. A. Friesner, R. B. Murphy, M. P. Repasky, L. L. Frye, J. R. Greenwood, T. A. Halgren, P. C. Sanschagrin, D. T. Mainz, Extra precision glide: Docking and scoring incorporating a model of hydrophobic enclosure for protein-ligand complexes. J. Med. Chem. (2006), doi:10.1021/jm051256o.

R. G. Sierra, C. Gati, H. Laksmono, E. H. Dao, S. Gul, F. Fuller, J. Kern, R. Chatterjee, M. Ibrahim, A. S. Brewster, I. D. Young, T. Michels-Clark, A. Aquila, M. Liang, M. S. Hunter, J. E. Koglin, S. Boutet, E. A. Junco, B. Hayes, M. J. Bogan, C. Y. Hampton, E. V. Puglisi, N. K. Sauter, C. A. Stan, A. Zouni, J. Yano, V. K. Yachandra, S. M. Soltis, J. D. Puglisi, H. DeMirci, Concentric-flow electrokinetic injector enables serial crystallography of ribosome and photosystem II. Nat. Methods (2016), doi:10.1038/nmeth.3667.

R. G. Sierra, H. Laksmono, J. Kern, R. Tran, J. Hattne, R. Alonso-Mori, B. Lassalle-Kaiser, C. Glöckner, J. Hellmich, D. W. Schafer, N. Echols, R. J. Gildea, R. W. Grosse-Kunstleve, J. Sellberg, T. A. McQueen, A. R. Fry, M. M. Messerschmidt, A. Miahnahri, M. M. Seibert, C. Y. Hampton, D. Starodub, N. D. Loh, D. Sokaras, T. C. Weng, P. H. Zwart, P. Glatzel, D. Milathianaki, W. E. White, P. D. Adams, G. J. Williams, S. Boutet, A. Zouni, J. Messinger, N. K. Sauter, U. Bergmann, J. Yano, V. K. Yachandra, M. J. Bogan, Nanoflow electrospinning serial femtosecond crystallography. Acta Crystallogr. Sect. D Biol. Crystallogr. (2012), doi:10.1107/S0907444912038152.

S. Nosé, A unified formulation of the constant temperature molecular dynamics methods. J. Chem. Phys. (1984), doi: 10.1063/1.447334

T. A. Halgren, R. B. Murphy, R. A. Friesner, H. S. Beard, L. L. Frye, W. T. Pollard, J. L. Banks, Glide: A New Approach for Rapid, Accurate Docking and Scoring. 2. Enrichment Factors in Database Screening. J. Med. Chem. (2004), doi:10.1021/jm030644s.

T. A. White, Processing serial crystallography data with crystFEL: A step-by-step guide. Acta Crystallogr. Sect. D Struct. Biol. (2019), doi:10.1107/S205979831801238X.

T. A. White, R. A. Kirian, A. V. Martin, A. Aquila, K. Nass, A. Barty, H. N. Chapman, CrystFEL: A software suite for snapshot serial crystallography. J. Appl. Crystallogr. (2012), doi:10.1107/S0021889812002312.

T. A. White, R. A. Kirian, K. R. Beyerlein, A. Aquila, A. V. Martin, L. Galli, C. H. Yoon, K. Nass, N. A. Zatsepin, A. Barty, C. Gati, F. Chervinskii, A. Tolstikova, W. Brehm, V. Mariani, P. de Waal, T. Nakane, K. Yamashita, O. Yefanov, S. Aplin, H. M. Ginn, T. Grant, M. Suzuki, N. Riebesel, Y. Gevorkov, O. Mor, CrystFEL - Release notes for version 0.9.0. 2020.

T. A. White, V. Mariani, W. Brehm, O. Yefanov, A. Barty, K. R. Beyerlein, F. Chervinskii, L. Galli, C. Gati, T. Nakane, A. Tolstikova, K. Yamashita, C. H. Yoon, K. Diederichs, H. N. Chapman, Recent developments in CrystFEL. J. Appl. Crystallogr. (2016), doi:10.1107/S1600576716004751.

T. Ichiye, M. Karplus, Collective motions in proteins: A covariance analysis of atomic fluctuations in molecular dynamics and normal mode simulations. Proteins Struct. Funct. Bioinforma. (1991), doi:10.1002/prot.340110305.

T. B. van Driel, S. Nelson, R. Armenta, G. Blaj, S. Boo, S. Boutet, D. Doering, A. Dragone, P. Hart, G. Haller, C. Kenney, M. Kwaitowski, L. Manger, M. McKelvey, K. Nakahara, M. Oriunno, T. Sato, M. Weaver, The ePix10k 2-megapixel hard X-ray detector at LCLS. J. Synchrotron Rad. (2020), doi:10.1107/S1600577520004257.

U Essmann, L. Perera, M. L. Berkowitz, T. Darden, H. Lee, L. G. Pedersen, A smooth particle mesh Ewald method. J. Chem. Phys. (1995), doi: 10.1063/1.470117.

V. Mariani, A. Morgan, C. H. Yoon, T. J. Lane, T. A. White, C. O’Grady, M. Kuhn, S. Aplin, J. Koglin, A. Barty, H. N. Chapman, OnDA: online data analysis and feedback for serial X-ray imaging. J. Appl. Crystallogr. (2016), doi:10.1107/S1600576716007469.

W. G. Hoover, Canonical dynamics: equilibrium phase-space distributions. Phys. Rev. A. doi: 10.1103/PhysRevA.31.1695

W. Kabsch, Xds. Acta Crystallogr D Biol Crystallogr., 66 (Pt 2), 125–32 (2010).

X. Q. Yao, B. J. Grant, Domain-opening and dynamic coupling in the α-subunit of heterotrimeric G proteins. Biophys. J. (2013), doi:10.1016/j.bpj.2013.06.006.

Y. Gevorkov, O. Yefanov, A. Barty, T. A. White, V. Mariani, W. Brehm, A. Tolstikova, R.-R. Grigat, H. N. Chapman, Acta Cryst. A75, 694–704 (2019).

